# The parasitic nematode *Strongyloides ratti* exists as populations of long-lived asexual lineages

**DOI:** 10.1101/2021.05.26.445462

**Authors:** Rebecca Cole, Nancy Holroyd, Alan Tracey, Matt Berriman, Mark Viney

## Abstract

Nematodes are important parasites of people and animals and in natural ecosystems they are a major ecological force. *Strongyloides ratti* is a common parasitic nematode of wild rats and we have investigated its population genetics using single worm, whole genome sequencing. We find that *S. ratti* populations in the UK consist of mixtures of asexual lineages that are widely dispersed across a host population. These parasite lineages are likely very old and may have originated in Asia from where rats originated. Genes that underly the parasitic phase of the parasite’s life cycle are hyperdiverse, compared with the rest of the genome, and this may allow the parasites to maximise their fitness in a diverse host population. These patterns of parasitic nematode population genetics have not been found before and may also apply to *Strongyloides* spp. that infect people, which will affect how we should approach their control.

## Introduction

Parasitic nematodes are important parasites of humans, livestock and other animals. In humans, parasitic nematodes are responsible for four of the twenty World Health Organization-defined Neglected Tropical Diseases (WHO 2021). In natural ecosystems parasitic nematodes are highly abundant and so a major force affecting host populations (Lafferty *et al.,* 2005; Kuris *et al*., 2008), so understanding parasites’ biology is critical in understanding wider ecological patterns and processes. Study of the population genetics of parasitic nematodes can give important insights into their biology and patterns of transmission in host populations. This has been studied extensively in nematodes parasitizing livestock, for example finding that they exist with very large effective population sizes, showing limited population genetic substructure, likely due to the high rate of livestock movement (Blouin *et al*. 1985; Redman *et al*., 2015; Sallé *et al*., 2019). The population genetics of parasitic nematodes infecting humans has also been investigated to understand their host range and zoonotic potential (Criscione *et al*., 2007; Thiele *et al*., 2018). In contrast, there has been much more limited study of the population genetics and genomics of nematodes infecting natural, unmanaged species (Cole and Viney, 2018).

*Strongyloides* spp. are a genus of parasitic nematodes, with two species – *S. stercoralis* and *S. fuelleborni* – infecting some 100-600 million people worldwide (Bethany *et al.,* 2006; Buonfrate *et al*. 2020). *S. ratti* is a common parasite of rats, *Rattus norvegicus* (Fisher and Viney, 1998). In the *Strongyloides* spp. life cycle hosts are infected by parasitic female worms only that reproduce parthenogenetically (Viney, 1994), producing eggs that pass out of the host in its faeces. Outside of the hosts, larvae can develop directly into infective larvae that then infect new hosts. Alternatively, and facultatively, larvae outside of the host can develop into a single generation of free-living adult males and females that reproduce sexually (Viney *et al*., 1993), with their progeny then also developing into infective larvae to infect new hosts. Genetically, this means that *Strongyloides* reproduces by obligatory mitotic parthenogenesis inside the host and by facultative sexual reproduction outside of the host. The choice between asexual, direct development and sexual, indirect development is affected by environmental conditions (particularly the host immune response and the temperature outside of the host), but genotypes also differ in their propensity for these two developmental routes (Viney *et al*. 1992; Harvey *et al*. 2000).

*Strongyloides’* mode of reproduction is very likely to affect its population genetics. If reproduction is exclusively by mitotic parthenogenesis, then the only source of genetic variation is mutation, and such a population would consist of an assemblage of different genetic lineages. In this scenario the mutation rate of a species is key in determining the extent of variation within a population. The occurrence of some sexual reproduction would allow genetic lineages of parasites to recombine, though the extent to which this will happen depends on the frequency with which sexual reproduction occurs. A three locus study of *S. ratti* in the UK found that it consists of one interbreeding population, likely mainly reproducing by direct, asexual reproduction (Fisher and Viney, 1998).

*S. ratti* has a compact 43 Mbp genome, consisting of 2 autosomes and an X chromosome, and its genome assembly is among the most contiguous for any nematode. This facilitates the population genomic analysis of wild *S. ratti.* Further, genomic, transcriptomic and proteomic analyses have identified genes that are putatively critical to *Strongyloides’* parasitic lifestyle. These were characterised in two ways: genes whose expression was significantly greater in the parasitic female stage compared with the free-living female stage; and genomic clusters of genes belonging to one of three families (encoding astacin-like metallopeptidases, CAP domain-containing proteins, acetylcholinesterases), which are gene families that have expanded as *Strongyloides* evolved to become parasites (Hunt *et al*., 2016).

In many, but not all, host-parasite systems, parasites can locally adapt to their host population, which enhances the fitness of those parasite genotypes (Greischar *et al.,* 2007). The genes and gene products underlying parasite local adaption are not well known. In *Strongyloides* the genes that have been shown to be central to its parasitism are at the interface between the parasite and its host and therefore may play a role in such adaptation. In such a scenario these genes could be under different selection pressures compared with the rest of the genome and so may have population genetic patterns that differ from the rest of the genome.

Here we report the whole-genome, fine-scale population genomics of *S. ratti* in a wild rat population, describing how parasite genotypes are distributed among individual hosts. We find that *S. ratti* populations in the UK exist as a mixture of asexual lineages and that these lineages are likely very long lived. Comparison of these lineages with historical, geographically dispersed samples shows that some of these lineages may be very widely spatially dispersed. We find that genes and gene clusters critical to the parasitic phase of the *S. ratti* life cycle are genetically hyperdiverse, compared with the rest of the genome. This hyperdiversity may contribute to *S. ratti*’s local adaption to its hosts within the context of it existing as long-lived asexual lineages.

## Results

### *S. ratti* is a common parasite of rats

We sampled rat faecal pellets from three sites in the southwest UK (**Figure 1**), from which we isolated 10,471 *S. ratti* infective larvae from 114 pellets (from a sample of 308). The proportion of infected faecal pellets significantly differed among the three sites (13, 47 and 62 % at sites CA, AM and LA respectively; χ^2^ = 48.9, *df* = 2, P < 0.0001) (**Figure 1; Supplementary Table 1**), but is consistent with a previous report of a high prevalence of *S. ratti* in wild rats in the UK (Fisher and Viney, 1998). The number of *S. ratti* larvae per pellet ranged from 1 – 1,730 (**Supplementary Figure 1**). While culturing these faeces for *S. ratti* we did not observe any sexual free-living adults. We genotyped rat faecal pellets to assign these to individual rats finding that 132 genotyped pellets belonged to 112 rats.

**Figure 1.**
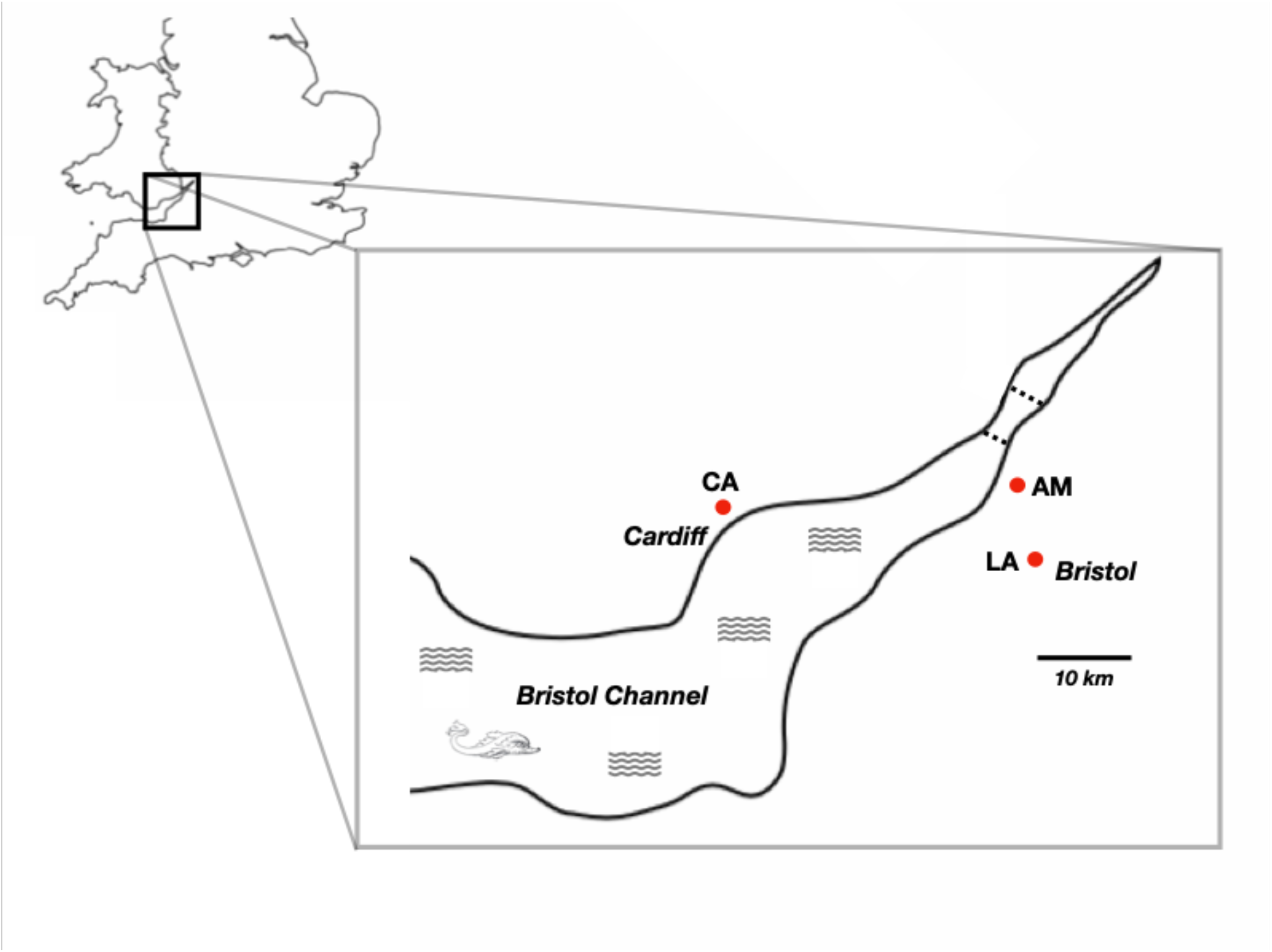
Map of sampling sites. Showing site LA near Bristol, AM on the English coast of the Bristol Channel, and CA on the Welsh coast of the Bristol Channel. Road bridges crossing the channel are shown as dotted lines; a train tunnel is not shown.

### *S. ratti* is partially genetically clustered at sample sites

We investigated the population genetics of *S*. *ratti* by analysing whole genome sequence of 90 individual infective larvae collected from the three sites. We identified 170,666 SNPs, giving an average density of 4.1 SNPs per kb. A total of 614 SNPs were tri-allelic, the remainder bi-allelic, with a ratio of 1.77 of transitions to transversions. Considering all SNPs together, the 90 parasites were in HWE (χ^2^ = 13.65, *df =* 19.35, P = 0.48) but this varied among SNPs. Overall, there was more heterozygosity than expected; specifically, allele frequencies predicted that 28% of SNP loci would be heterozygous in individual worms, whereas 36% (range 28-58%) were.

We analysed the pattern of parasite population genetic variation (i) within and among rat hosts, and (ii) within and among sampling sites, finding evidence of some differentiation of the parasites among the sample sites. The pairwise relatedness (Φ) among the 90 parasites was non-normal (Shapiro-Wilkes test for normalcy W = 0.902, P < 0.0001), suggesting genetic clustering among the worms at some level (**Supplementary Figure 2**). *S. ratti* parasitic females reproduce by mitotic parthenogenesis (Viney, 1994) and so siblings will be genetically identical save for individual-specific mutations. We did not detect putative sibling parasites within individual rats, despite sampling up to 4 parasites from each rat. (**Supplementary Figure 3**). This suggests that faecal pellets commonly contained larvae of more than 4 genotypes. In faecal pellets containing ≥ 10 larvae, if there indeed had only been 4 genotypes present then our chance of detecting the 4 would have been ≤ 0.19. It is more likely that faecal pellets contained worms of more than 4 genotypes. Specifically, we had a ≥ 0.50 chance of detecting 4 unique genotypes when: ≥ 6 genotypes were actually present in pellets containing 12-18 larvae; ≥ 7 genotypes were present in pellets containing 19-31 larvae. The average relatedness among pairs of parasites from the same rat and among parasites from different rats (Φ = 0.22 and 0.214, respectively), was not significantly different (t = -0.32, *df* = 108.81, P = 0.75). Thus, we conclude that individual rats contain genetically diverse parasites (consistent with our *S. ratti* RFLP genotyping, see Materials and Methods section), meaning that genetic clustering of parasites is not at the level of the rat.

However, parasites from the same sample site were more closely related (mean same site Φ = 0.225; single, same site Φ, CA = 0.128, LA = 0.258, AM = 0.227) than parasites from different sample sites (Φ = 0.206) (t = -3.68, *df* = 3975.9, P ≤ 0.001) (**Figure 2**). Overall, F_ST_ was very low (0.02), and indeed zero between parasites at the two English sample sites (**Figure 2**). Together, these results show that there is some genetic clustering of *S. ratti* at the level of the sampling site, but not at the level of individual rats.

**Figure 2.**
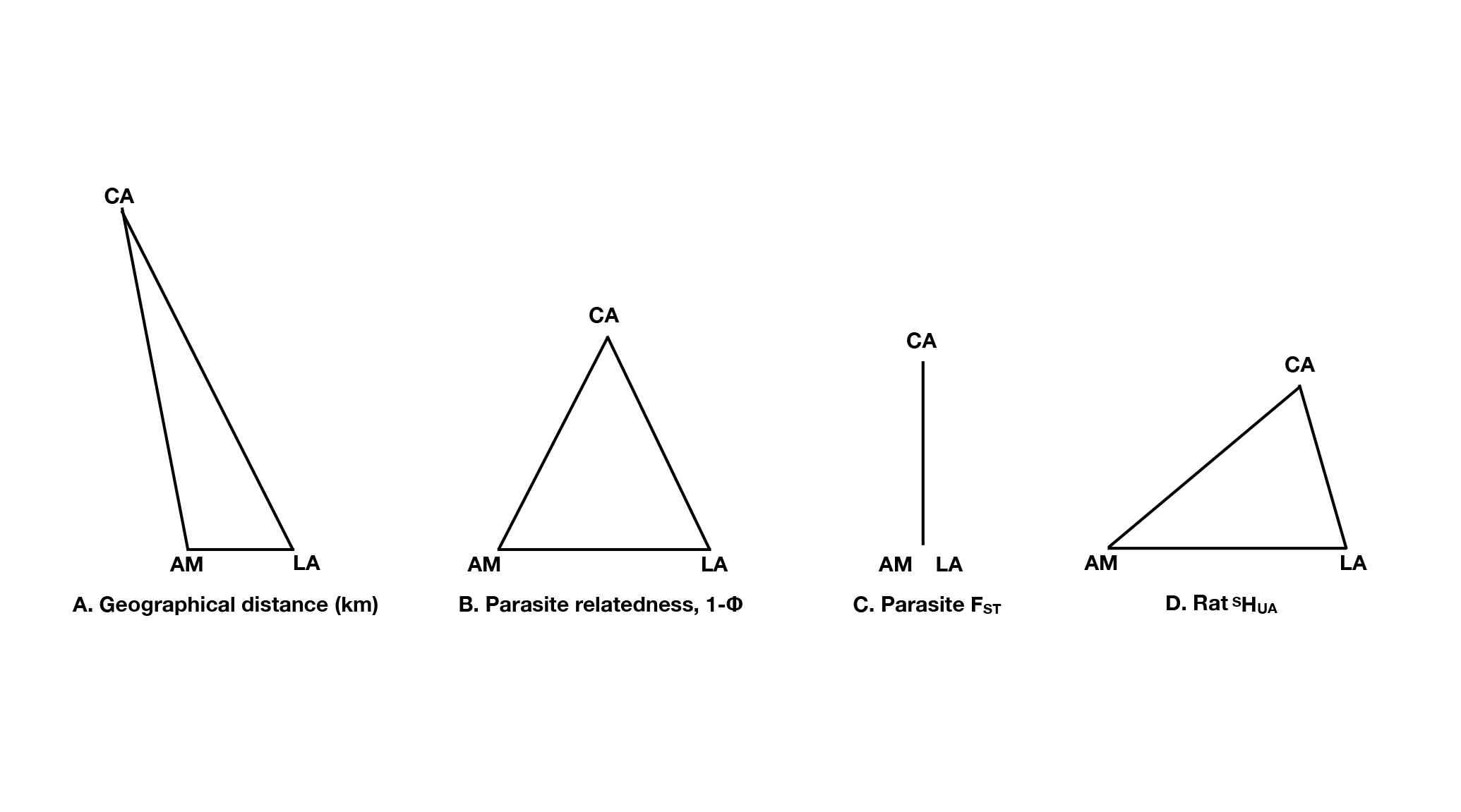
Physical and genetic differences among parasite and host populations. A schematic view of (A) Geographical distance among the sample sites, where direct distances between the sites are AM-LA = 9.2 km, CA-AM = 31.5 km, CA-LA = 33.5 km. (B) average pairwise relatedness (shown as 1-Φ) where Φ AM-LA = 0.244, CA-AM = 0.157, CA-LA = 0.162, (C) FST among parasites, from the three sample sites where, AM-LA = 0, CA-AM = 0.03, CA-LA = 0.03 and (D) ^S^H_UA_ among rats where, AM-LA = 0.21, CA-AM = 0.22, CA-LA = 0.15. Note, the scales of the metrics varies among the panels.

### *S. ratti* consists of divergent genetic clades that are widely distributed

We examined the parasite diversity more closely by constructing neighbour-joining dendrograms. This revealed five parasite clades, with most worms (78 of the 90) in one of three clades (clades 1 – 3) (**Figure 3A**). Maximum likelihood trees also confirmed the existence of these clades (**Figure 3B**; **Supplementary Figure 4**). We examined the admixture among the 90 parasites, which most reliably grouped the parasites into five genetic groups, corresponding to the five clades defined by the neighbour-joining tree (**Figure 3C; Supplementary Figure 5**). Moreover, even under other subdivision scenarios there is no evidence for gene flow (**Supplementary Figure 5**).

**Figure 3.**
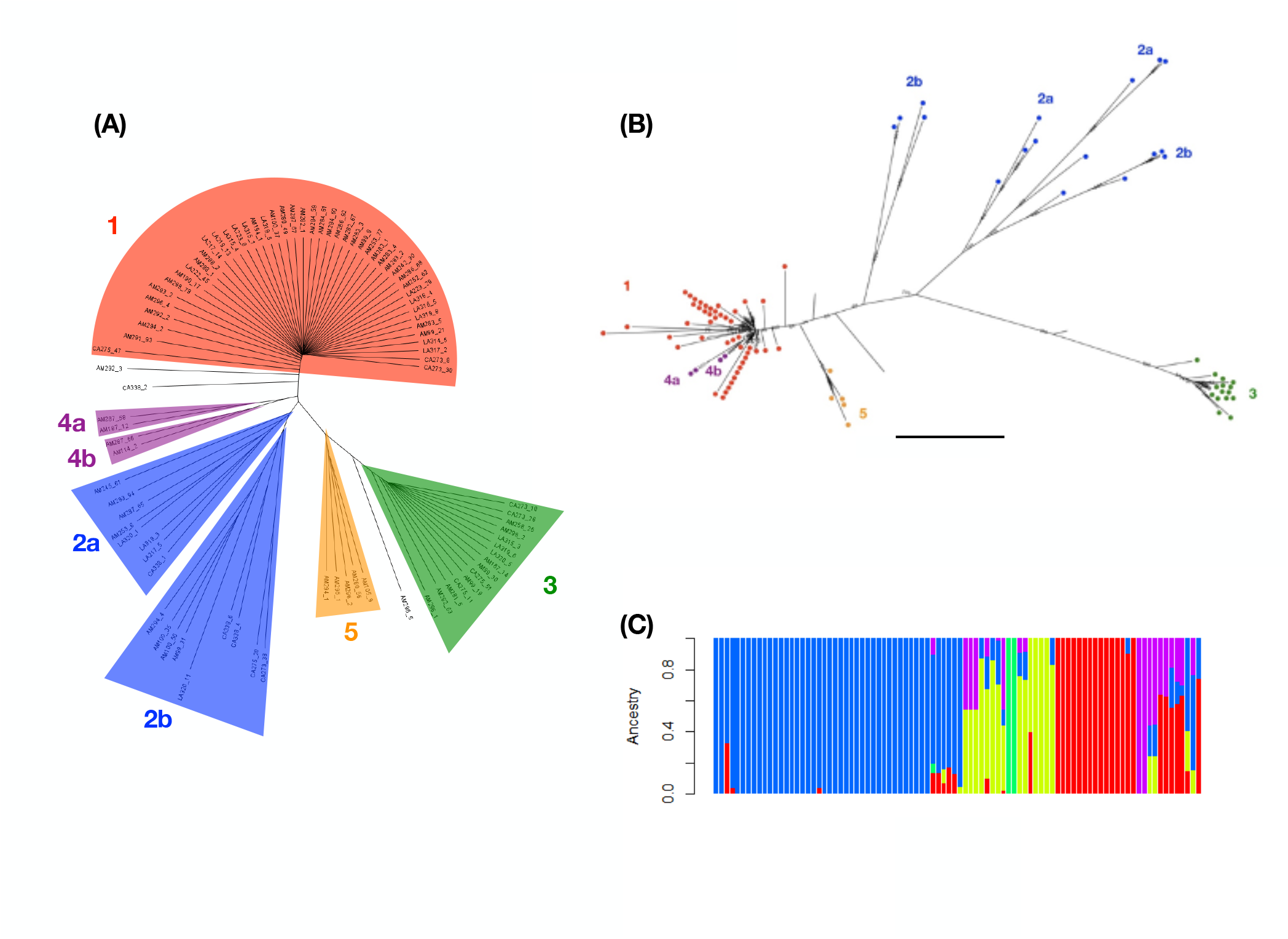
*S. ratti* consists of distinct genetic clades that are widely distributed. (A) A neighbour-joining dendrogram showing the five clades; (B) a maximum likelihood tree based on chromosome 1 where individuals are colour coded according to their clade membership in the neighbour-joining tree; the scale bar is 2 x 10^-4^ substitutions per site; chromosome specific trees are shown in **Supplementary** Figure 4; (C) the admixture of the 90 larvae for K = 5, which is the most strongly supported value of K; the order of individual worms and their neighbour-joining tree clade membership is shown in **Supplementary** Figure 5. Note, the colour coding in (C) does not correspond to (A) or (B).

Parasites belonging to the three main clades were present in all three sampling sites, in ratios expected based on the number of sequenced parasites from each sampling site (Fisher’s exact test, P = 0.14). Individual rats contained parasites from multiple clades; specifically, 11 rats contained parasites from two clades, and 3 rats contained parasites from three clades. Principal Component Analysis produced similar results, showing clustering of parasites within clades, and that these clades were dispersed across the three sampling sites (**Supplementary Figure 6**).

Analysis of Φ and F_ST_ within and among clades 1 – 3 is consistent with *S. ratti* being structured into sympatric, genetically distinct clades, with the three clades dispersed across the three sampling sites. Specifically, Φ was high within clades, especially clades 1 and 3 (0.43 and 0.45, respectively), but lower between clades; F_ST_ among the clades was 0.3 and similar (0.22 – 0.35) between clades (**Table 1**). Given that we did not observe any free-living sexual stages of *S. ratti* this further suggests that these clades are asexually derived and maintained.

**Table 1.** F_ST_ and Φ among *S. ratti* clades 1, 2 and 3. Pairwise F_ST_ is shown in italics above the diagonal, and Φ on and below the diagonal in non-italic text.

|  | Clade 1 | Clade 2 | Clade 3 |
| --- | --- | --- | --- |
| Clade 1 | 0.43 | *0.22* | *0.35* |
| Clade 2 | 0.18 | 0.23 | *0.23* |
| Clade 3 | 0.06 | 0.05 | 0.45 |

The population genetic structure of *S. ratti’s* mitochondrial genome consists of three divergent genetic clades that are not strongly associated with sampling sites, which is consistent with the nuclear genome results. Specifically, in the mitochondrial genome we identified 156 SNPs (average density 9.3 SNPs per kb) identifying 58 haplotypes among the 90 parasites. There was a strong, positive correlation between the pairwise similarities of mitochondrial and nuclear genomes (Mantel test, r = 0.76, P < 0.01), and between minimum spanning maps of mitochondrial haplotypes and neighbour-joining trees of nuclear genomes (**Supplementary Figure 7**). The number of mitochondrial SNPs in same sampling site and different sampling site comparisons (23.6 and 24.2, respectively) were not significantly different (t = -0.98, *df* = 3771, P = 0.33).

We wanted to investigate if the rat host population genetic structure enforced a population genetic structure on the parasites because of the partitioning of parasites among individual hosts. To do this we investigated the population genetics of the rats by genotyping faecal pellets at nine microsatellite locus (**Supplementary Table 2**). These loci were generally not in Hardy-Weinberg equilibrium (HWE); specifically 8, 3 and 5 of the 9 loci at sites CA, AM and LA, respectively, were not (**Supplementary Table 3**). Rat allele frequencies differed markedly among sampling sites, consistent with restricted rat gene flow among the sites. The pairwise relatedness among rats was higher within sites than among sites (average Ritland and Lynch relatedness values 0.06 and -0.06, respectively), and the distribution of these relatedness values at each site had a right-hand skew (**Supplementary Figure 8**), showing more closely related pairs of individual rats within sites than expected by chance. We could assign rats to each sample site based on allele frequencies with 89% accuracy. Shannon’s mutual information index (^S^H_UA_), which does not assume HWE (Hedrick, 2005; Jost *et al.,* 2018), shows that there is moderate genetic differentiation among rats from the three sites (**Figure 2**). Together these results – differences in allele frequencies among sample sites, higher within-site than among-site relatedness, high accuracy in assignment to sample site, moderate values of ^S^H_UA_ – show that there is genetic differentiation among rats at the three sampling sites. The evidence of moderate genetic differentiation among rats at sample sites is broadly consistent with the geographical separation of the three sites, and with limited migration of rats among the sites (**Figure 1**), as has been observed with urban rats (Combs *et al*. 2017). Notably, the genetic structure of the *S. ratti* populations does not mirror that of its rat hosts.

We sought to understand the age of the *S. ratti* clades. We did this using the ML trees of chromosomes 1 and 2 to calculate the number of substitutions per site that have occurred since the last common ancestor (LCA) of (i) clade 3 and (ii) clades 1 and 3. Assuming neutrality and the *C. elegans* mutation rate of 2.7 x 10^-9^ per site per generation (Denver *et al.,* 2009), we calculate that there were an average of 18,401 (range 17,037 – 20,195) and 322,011 (range 213,185 – 393,589) generations since the LCA of clade 3 and of clades 1 and 3, respectively. To convert these generations to years we assumed, conservatively, that there are two *S. ratti* generations per year in the wild, noting that *S. ratti* parasitic females have a maximum lifespan of a year when infecting immunodeficient laboratory rats (Gardner *et al*. 2006). On this basis the LCA of clade 3 existed approximately 9,000 years ago (*i.e.* 1200 CE) and the LCA of clades 1 and 3 approximately 161,000 years ago, which is the Middle Pleistocene (now renamed the Chibanian). Given the dates, it is a reasonable scenario that the LCA of these *S. ratti* clades evolved outside of the UK and arrived in the UK with their hosts. *R. norvegicus* is thought to have arrived in mainland Europe from Asia about 1,800 years ago (*c.* 200 CE) and into the British Isles from *c.*1750 CE (Lever 1977; Zeng *et al*. 2018), where the black rat (*R. rattus*) had likely been since *c.* 1100 CE. Clades 1 and 3 may well have emerged in Asia where *R. norvegicus* originated, and the LCA of clade 3 perhaps in mainland Europe.

We also investigated the spatial and temporal extent of the *S. ratti* clades by examining whole genome sequences of 10 *S. ratti* isofemale lines derived from rats sampled from the UK and Japan between 1989 and 2012 (**Supplementary Table 4**). A neighbour-joining dendrogram of these isofemale lines and the 90 wild parasites (sampled in 2017/18) showed that these isofemale lines are not genetically distinct from the 90 parasites, occurring in clades 1, 2 and 4 (**Supplementary Figure 9**). This observation – particularly of the two Japanese isofemale lines – is consistent with an Asian ancestry of the 90 contemporary UK genotypes, and more generally with the idea that *S. ratti* genotypes exist as long lived, asexually maintained lineages.

In summary, these results show that *S. ratti* populations consist of mixtures of different genetic clades consisting of asexually maintained lineages that are likely very long lived – over thousands and tens of thousands of years – and widely distributed across the global host population. The *S. ratti* life cycle is obligately mitotically parthenogenetic (Viney 1994), with facultative sexual reproduction (Viney *et al.,* 1993); the population genetic structure that we have observed suggests that sexual reproduction occurs vary rarely, if at all, in these populations. We observe no obvious geographical localisation of the different parasite clades, which might be expected with geographically-based host local adaptation. Indeed, given the likely asexual reproduction of these populations it would therefore appear that *S. ratti* cannot genetically adapt to its local host populations except through mutational processes.

### *S. ratti* genes involved in parasitism are highly diverse

We next investigated the diversity of genes that *S. ratti* uses in the parasitic phase of its life cycle. We focussed on two sets of genes: (i) “parasitism genes”, which are genes whose expression is at least one log_2_-fold greater in the parasitic female morph compared with the free-living female morph, and (ii) “expansion clusters”, which are genomic regions containing four or more genes encoding members of either astacin-like metallopeptidase, CAP domain-containing protein, or acetylcholinesterase families, which previous analyses have identified as having expanded with the evolution of parasitism in *Strongyloides* (Hunt *et al.,* 2016).

Genetic diversity in *S. ratti* is not evenly distributed across the *S. ratti* genome (**Supplementary Figure 14**), and we identified high SNP diversity regions, consisting of ≥ 200 SNPs per 10 kb. We first asked how often parasitism or free-living (which are *vice versa* compared with parasitism genes) genes occurred in these high SNP diversity regions. We found that parasitism genes were significantly overrepresented within these high diversity regions, compared with their representation in highly conserved genomic regions (≤ 4 SNPs per 10 kb region), or across the genome as a whole (**Figure 4A; Supplementary Table 5**). In contrast, free-living genes were represented at the same rate in high diversity regions, highly conserved regions, and across the genome as a whole (**Figure 4A**).

**Figure 4.**
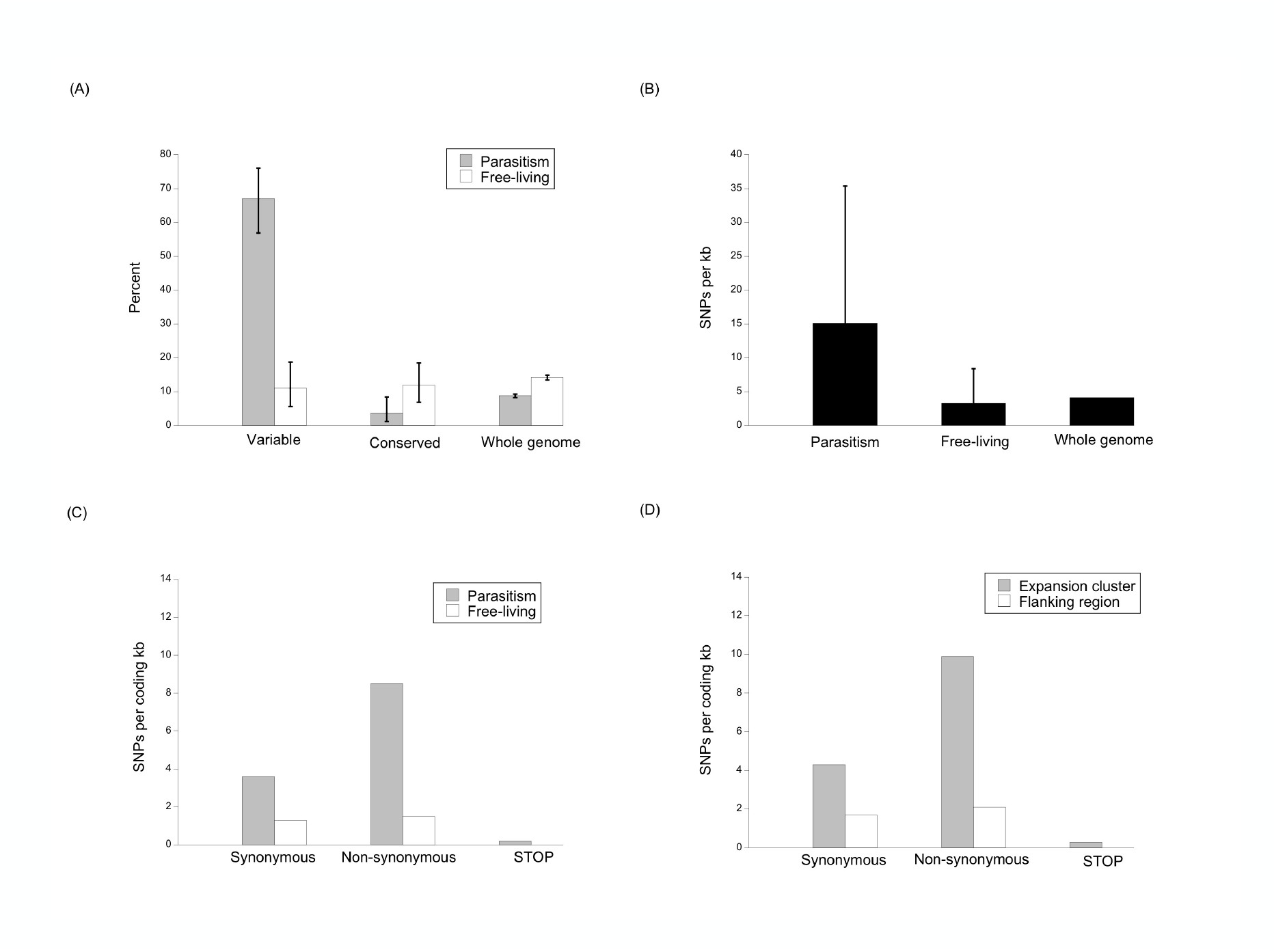
*S. ratti* genes involved in parasitism are highly diverse. (A) The percentage of genes in high diversity regions (≥ 200 SNPs per 10 kb), conserved regions (≤ 4 SNPs per 10 kb), or across the whole genome as a whole, that are parasitism or free-living genes. There were 100 genes in the variable regions, 137 in the conserved regions, and 12,464 across the whole genome. Parasitism and free- living genes are as defined by Hunt *et al.,* 2016. Errors bars are 95 % confidence intervals. (B) The number of SNPs per kb (+1 SD) in parasitism genes (range 0-85.9), free-living genes (0-41.5) or across the genome as a whole. The SNP density for parasitism and free- living genes is calculated from coding sequence only; for the whole genome, all sequence data are used and so no SD is given. (C) The density of SNPs of different effect in parasitism and free-living genes. (D) The density of SNPs of different effect in expansion clusters and flanking regions.

Furthermore, two classes of genes – those encoding astacin-like metallopeptidases and CAP domain-containing proteins, both of which are associated with parasitism in *S. ratti* – were more common in these high diversity regions, compared with the genome as a whole. Specifically, 5.6 and 11.8 % (95% confidence intervals 2.6-10.3 and 6.3-16.4 %, respectively) of genes in these regions encode astacin-like metallopeptidase and CAP domain-containing proteins, which is higher than their representation across the whole genome (1.5 and 0.7 %, respectively).

We compared the SNP density among the hundred most parasitic and most free-living genes, though we excluded six of the parasitism genes due to concerns about their underlying sequence assembly, leaving 94 parasitic genes (**Supplementary Tables 6 and 7**). Parasitism gene SNP density was approximately four times that of free-living genes, or that of the genome as a whole (**Figure 4B**). The SNPs in these parasitism genes mainly code for non-synonymous substitutions, rather than synonymous substitutions, which contrasts with free-living genes, where the rate of both types of SNPs occurred with similar frequency (**Figure 4C**).

We also found evidence of high genetic diversity in *S. ratti*’s expansion clusters, compared with the flanking regions (**Supplementary Table 8**). SNPs were three times more dense in the expansion clusters than in the flanking regions (SNP density (SD) per kb 15.9 (18.3) and 4.6 (7.2), in expansion clusters and flanking regions, respectively). Strikingly, SNPs within expansion clusters were approximately twice as likely to code non-synonymously, rather than synonymously, unlike the flanking regions where these rates were similar (**Figure 5D**); a pattern also seen for each individual expansion cluster, except clusters 1 and 8 (**Supplementary Table 5**).

We thought about possible reasons for the elevated genetic diversity in genes involved in *S. ratti*’s parasitism. Parasites can locally adapt to their hosts to maximise parasite fitness, and we hypothesised that evolution of the expansion clusters, independent of the rest of the genome, could be a means by which host local adaptation occurs, where the expansion clusters evolve differently from the rest of the genome. To investigate this, we created neighbour-joining trees based on individual expansion clusters to see if they strongly diverged from the whole genome-based trees. However, in these expansion cluster-specific trees, the whole genome-defined clades 1 and 3 were still strongly evident (**Supplementary Figure 15**). We next measured the selection in the expansion clusters and their flanking regions, but could also find no consistent evidence for diversifying selection in the expansion clusters compared with their flanking regions (**Supplementary Table 9**). These results therefore do not support the idea that the expansion clusters are locally adapting to host genotypes.

Together, these observations – that highly variable genomic regions have an over- representation of parasitism genes, and of astacin-like metallopeptidase and CAP domain- containing coding genes; that parasitism genes have SNP densities that are higher than those of free-living genes; that parasitism genes have a comparative excess of non- synonymous-coding SNPs; that expansion clusters have higher SNP densities and an excess of non-synonymous-coding SNPs – shows that in *S. ratti* there is a concentration of genetic diversity within genes and genomic regions that very likely play a key role in the parasitic phase of its life cycle.

## Discussion

Parasitic nematodes are ubiquitous parasites of animals and are partitioned among individual hosts between which they must transmit, and so parasite and host biology can affect their population genetic structure ((Blouin *et al*. 1985; Cole and Viney, 2018). Understanding the population genetics of parasitic nematodes can give an insight into their population biology, which is poorly known outside of species infecting humans and livestock (Cole and Viney, 2018).

For the facultatively sexual parasite of rats *S. ratti* we have discovered that in the UK its population consists of a mixture of genetically diverse clades, where those clades are widely dispersed across host populations (and possibly on a global scale), with very little evidence of structuring across three host populations. By estimating the age of the last common ancestor of these clades we find that these parasites lineages are very long lived, existing over some thousands to tens of thousands of years. This result does raise the question of whether these different lineages should be consider as different species or sub-species. Considering the age of these lineages with the history of the spread of the brown rat from Asia into mainland Europe and then to the British Isles, a plausible scenario is that the *S. ratti* lineages originated in Asia and spread to the UK, and likely the world. It is notable that the neighbour-joining dendrograms shows Japanese-derived *S. ratti* isofemale lines among the UK-derived genotypes. More extensive global sampling of *S. ratti* genetic diversity is now warranted.

This is a pattern of population genetic variation that has not, as far as we are aware, been observed in a parasitic nematode before. The life cycle of *S. ratti* contains an obligatory asexual parthenogenetic stage (Viney, 1994), as well as a facultative sexual stage (Viney *et al*., 1993). We observed no sexual stages during our work, suggesting that sexual reproduction is very rare, or even absent, in these UK populations, consistent with previous observations (Viney *et al*. 1992; Fisher and Viney, 1998). The population genetics of the populations that we studied is also consistent with the absence of sexual reproduction. Other lines of *S. ratti* (from Japan and a long-term lab adapted line originally isolated from the United States) show some development of free-living males and females, suggesting that some sexual reproduction may occur in wild *S. ratti* populations beyond the UK (Viney *et al.,* 1992). Even low levels of sexual reproduction can be effective in the spread od alleles in a population, especially with strong selection (Hartfield, 2016).

In contrast to the parasites’ population genetic patterns, the host rat populations did show some evidence of population genetic differentiation among the different sample sites, consistent with restricted movement of rats between the sites. Studies of the population genetics of rats in cities have also shown evidence of limited dispersal of rats and so population genetic differentiation among rats in different city regions (Gardner-Santana *et al.,* 2009).

Understanding a parasite’s population genetic structure can be important in understanding the host range of a parasite, which is of applied interest for parasites of humans. For example, population genetic analysis has been used to understand the possibly changing host range of Guinea worm (*Dracunculus medinensis*) in human and dog hosts during sustained control efforts in human populations (Durrant *et al*., 2020). There is considerable interest in understanding the host range of *S. stercoralis* that infects people; specifically, *Strongyloides* in dogs has been considered to be a source of human infection (Jaleta *et al.,* 2017). If the population genetic structure we have observed with *S. ratti* – a mixture of asexually maintained clones widely distributed across host populations – also pertains in *S. stercoralis*, then there is likely to be considerable complexity in understanding the population genetics and host range of *S. stercoralis* genotypes. Where a species exits as a collection of asexually maintained lineages, then each lineage could diverge genetically, which raises the possibility that *S. stercoralis* could exist as a mixture of lineages each with different host ranges, for example some able only to infect people, some only able to infect dogs and some intermediate. Current approaches to studying the host range of *S. stercoralis* have commonly only used single or a few loci, which cannot resolve more complex patterns of population genetic variation. Whole genome analysis (following whole genome amplification) of *S. stercoralis* from people in Japan and Myanmar found differences between parasites from these two sources (Kikuchi *et al.,* 2016). If population genetic structure similar to *S. ratti’s* also occurs in *Strongyloides* spp. infecting people and livestock, then there will be consequences for the evolution of anthelminthic resistance. In the absence of sexual reproduction, anthelmintic resistance could evolve separately in different *Strongyloides* lineages, but would not be passed among them. The effect of this might be to slow the development of anthelminthic resistance in *Strongyloides* populations, compared with obligatory sexually reproducing nematodes, though with *Strongyloides* sustained anthelminthic use would, of course, select against anthelminthic sensitive genotypes.

Genomic analyses of *Strongyloides* have discovered genes and gene families that play a critical role in the parasitic phase of its life cycle. We have discovered that in wild *S. ratti* both parasitism genes and genes in expansion clusters are highly diverse compared with other genes. Of particular note, many of these SNPs in the parasitism genes and genes in expansion clusters we observed are predicted to code for non-synonymous substitutions, meaning that this genetic diversity may cause functional differences in the gene products. The parasitism genes and genes in expansion clusters are dominated by large gene families. Large families can allow genetic diversity to accumulate among gene family members because any potential negative fitness consequence of a mutation in one gene of a family, could be effectively ameliorated by others in that family. In this way, large gene families can allow the exploration of genetic space.

Our observation of comparatively high genetic diversity in these genes is particularly interesting in light of recent observations of genetic diversity in the free-living nematode *Caenorhabditis* spp. Specifically, analysis of worldwide populations of *C. elegans* find that genetic variation is concentrated in a number of genomic regions (56 and 19 kb mean and median size, respectively), with evidence suggesting that diversity in these regions is maintained by balancing selection (Lee *et al*., 2021). In the *C. tropicalis* genome, genetic variation is also distributed heterogeneously across its genome, for example with some 140 high genetic diversity classified regions extending for more than 30 kb (Noble *et al*., 2021). While there is a superficial similarity between the patterns of genomically concentrated genetic diversity in two *Caenorhabditis* species and in *S. ratti*, the mechanisms generating these patterns might be different. Notably, in *S. ratti* we did not find any evidence of diversifying selection in the expansion cluster genes. What these two *Caenorhabditis* studies and the present study do demonstrate is that detailed, whole genome analysis of wild individuals is uncovering hitherto un-expected patterns of genomic diversity, which are likely to exit in other taxa too, and which need to be investigated.

Parasites have commonly been found to locally adapt to their host populations (Lively *et al*., 2004; Greischar and Koskella, 2007). If such a phenomenon was occurring in *S. ratti* then it may be manifest as geographical clustering of parasite genotypes *per se* or geographical clustering of parasitism gene genotypes and / or expansion cluster genotypes. However, the dispersion of *S. ratti* genotypes and of the genetic diversity in parasitism genes and expansion clusters that we have observed does not show any suggestive signatures of such local adaptation to hosts.

Alternatively, mindful that the products of the parasitism genes and genes in expansion clusters interface with the host, some variants of these genes may give a parasite a fitness advantage when infecting certain host genotypes, compared with parasites with other variants. Each individual parasite’s suite of these genes may therefore represent a combination of different variants that have been selected for as these parasite lineages have over their history infected a range of host genotypes. In this scenario, these parasites are not locally adapted to their host genotypes *pe se*, but rather have available a set of gene variants that are appropriate for a wide range of already-encountered host genotypes. This idea – the so-called grey pawn hypothesis – that the aggregate effect of a large number of diverse genes is selected so as to cover a large phenotypic space has previously been suggested in trying to understand the very large number of *Caenorhabditis* spp. chemoreceptor-coding gene families in (Thomas and Robertson, 2008); this idea has also been applied to hookworm parasites (Schwarz *et al*., 2015). It is clear that that further research is needed to understand the evolution of large gene families, as well as the full biological significance of high levels of genetic diversity in genes underlying *S. ratti’*s parasitism.

Our whole-genome, single worm analysis of wild *S. ratti* is a non-destructive method of sampling parasite genetic diversity, and the hope must be that these approaches are now expanded to other parasitic nematodes. Such analyses will likely uncover different patterns of population genetic variation in other species, and understanding this will more fully illuminate the interactions between parasites and hosts, and so underpin a better understanding of the rich ecology of parasites. If *Strongyloides* infecting people has a similar population genetic structure, then this may have consequences for understanding *Strongyloides* host range and so zoonotic potential, as well as for the evolution of anthelmintic resistance.

## Materials and Methods

### Parasite and rat sampling

We sampled at three sites in the southern UK – Avonmouth (AV), Cardiff (CA) and Long Ashton (LA) (**Supplementary Table 10, Figure 1**), collecting fresh rat faecal pellets, which were cultured at 19°C and visually inspected for *S. ratti* infective third stage larvae (Viney and Lok, 2007), which were washed twice in distilled water, once in 1 % w/v SDS, and then twice more in distilled water, before being stored at -80°C. We genotyped rat faecal pellets at 9 dinucleotide repeat microsatellite loci (**Supplementary Table 2**) that had previously been used with wild rats (Desvars-Larrive *et al.,* 2017; Giraudeau *et al*., 1999; Gardner-Santana *et al.,* 2009; Steen *et al*., 1999), preparing DNA using the QIAamp DNA Stool Mini Kit (Qiagen) (Cole, 2020).

### *S. ratti* genotyping

Of the more than 10,000 *S. ratti* larvae that we isolated from wild rats, we had to select a sub-sample for whole genome sequencing. We did this mindful that *S. ratti* parasitic females reproduce by mitotic parthenogenesis (Viney, 1994), such that in pellets containing more than one larva, that those larvae may be genetically identical siblings. Alternatively, a rat may be infected with multiple genotypes of *S. ratti* in which case there will also be a genetically different larvae in individual faecal pellets. We used RFLP genotyping to initially assess the genetic diversity among larvae within individual faecal pellets (Cole 2020). This showed that *S. ratti* infrapopulations are typically composed of multiple, genetically distinct parasitic adults and so we concluded that whole genome sequencing of multiple infective larvae from the same faecal pellet was unlikely to result in extensive resequencing of genetically identical siblings, and as such would be informative both for measuring the genetic diversity within sampling sites as a whole, and for assessing the extent of genetic variation within infrapopulations (Cole 2020)

For whole genome sequencing larvae were lysed and DNA prepared (Cole 2020). Samples were quantified with Biotium Accuclear Ultra high sensitivity dsDNA Quantitative kits using Mosquito LV liquid platform, Bravo WS and BMG FLUOstar Omega plate reader and cherrypicked to 200 ng / 120 μL using a Tecan liquid handling platform. Cherrypicked plates were sheared to 450 bp using a Covaris LE220 instrument and post-sheared samples purified using Agencourt AMPure XP SPRI beads on Agilent Bravo WSLibraries were constructed using the NEB Ultra II custom kit on an Agilent Bravo WS automation system. PCRs were set-up using KapaHiFi Hot start mix and IDT 96 iPCR tag barcodes on an Agilent Bravo WS automation system, and then purified using Agencourt AMPure XP SPRI beads on Beckman BioMek NX96 liquid handling platform. Libraries were quantified with Biotium Accuclear Ultra high sensitivity dsDNA Quantitative kit using Mosquito LV liquid handling platform, Bravo WS and BMG FLUOstar Omega plate reader. Libraries were pooled in equimolar amounts on a Beckman BioMek NX-8 liquid handling platform and libraries normalised to 2.8 nM ready for cluster generation on a c-BOT and loading on the Illumina X Ten platform. Sequencing reads from the libraries were aligned to the *S. ratti* reference assembly version 5_0_4 (Hunt *et al*. 2016, taken from WormBase ParaSite release WBPS7) using Bowtie 2 version 2.2.9 (Langmead and Salzberg, 2012) with default settings. We initially whole genome sequenced 225 *S. ratti* infective larvae at low depth of coverage and calculated the proportion of reads that aligned to the *S. ratti* genome, and used this metric to choose 90 libraries for further deep sequencing.

### Sequence analysis

We used BCFtools (Li 2011) to identify SNPs using the criteria that they (i) fell on a nucleotide covered by at least 1,000 reads (cumulative across all samples), (ii) had a mean mapping quality of at least 20, and (iii) had a QUAL score of at least 50. Among the 90 *S. ratti* genome sequences, nucleotides that were identical among all samples (but different from the ED321 reference genome) were removed. We sequenced to an average coverage of 96 % of nucleotides (range 75.8-99.3 %), and an average read depth of 68 (range 20- 246; just 5 larvae had mean read depths of less than 30).

We noticed that the mean read depth on the X chromosome was 67.9 % of the mean read depth on the two autosomes. We concluded that this was due to the GC content because (i) there was a significant correlation between read depth and GC content (**Supplementary Figure 16**) and (ii) that the X chromosome has a slightly lower GC content (19.7 %) compared with the autosomes (22 %) (Hunt *et al*., 2016).

Basic genetic diversity and population genetic statistics were calculated using VCFtools version 0.1.12 (Danecek *et al*., 2011). Hardy-Weinberg equilibrium (HWE) was calculated considering only biallelic SNPs. Φ relatedness values (Manichaikul *et al.,* 2010) of each pair of larvae were calculated using VCFtools and we used t-tests to compare Φ values. We also measured the differentiation among sites using the fixation index, F_ST_.

We generated neighbour-joining dendrograms using TASSEL 5.0 (Bradbury *et al.,* 2007), and visualised these in FigTree Version 1.4.3. Clades within the neighbour-joining trees were identified by eye. Fisher’s exact tests, performed in R, were used to determine whether there were significant differences in the frequencies of these clades among sampling sites or sampling seasons.

We constructed maximum likelihood trees of the 90 parasites, producing consensus fasta sequences for each individual, but where an individual was heterozygous the reference allele was applied, with sequences aligned with MAFFT version 7 (Katoh *et al*. 2009) using strategy FF-NST-1 for fast alignment, and maximum likelihood tree estimation performed using RaxML version 8.1.15 (Stamatakis 2006), using the general time reversible gamma model of substitution rate heterogeneity, and rapid bootstrapping with 100 replicates. We generated separate maximum likelihood trees for chromosome 1, the first 80 Mb of chromosome 2, the remainder of chromosome 2 and the two largest contigs of the X chromosome.

We conducted Principal Component Analysis using the R package pcadapt version 4.1.0 (Luu *et al*. 2017) using only loci with a minor allele frequency greater than 0.05. We investigated the admixture among the 90 parasite genotypes using ADMIXTURE version 1.3.0 (Alexander and Large 2011). Due to computational constraints, for the 90 parasites SNP data were first thinned so that no two SNPs were within 500 bp of each other, leaving a dataset of 35,559 SNPs. ADMIXTURE was run separately for k values 2-15.

We measured linkage disequilibrium (LD) among the 90 samples for the two autosomes and the two largest X chromosome scaffolds. We initially phased the genotype data into haplotypes using Beagle version 5.0 (Browning and Browning, 2007; Browning *et al.,* 2018), where we used 100 burn-in iterations to generate an initial estimate of haplotype frequency, and a further 100 iterations were used to estimate genotype phase for each SNP in each sample. Phasing is influenced by the effective population size (Ne), which isn’t known for *S. ratti*, but we estimated this as 50,000; otherwise, default Beagle parameters were used. We also undertook phasing using Shapeit version 2-r900 (O’Connell *et al*. 2014), where we used 100 burn-in iterations, 100 phasing iterations, and an estimated Ne of 50,000. For both we used a window size of 0.5 Mb to estimate haplotypes. Only biallelic loci were used in Shapeit, but triallelic loci were also included in Beagle. We report the results from phasing using Beagle; Shapeit gave similar results.

To reduce computational time during linkage decay analysis, phased VCF files were thinned so that no two remaining loci were within 100 bp of one another. To perform linkage decay analysis, VCFTools was used to compare each SNP to each other SNP within a 50 kb window of it, with Pearson’s coefficient of correlation, r^2^, calculated for each pair. To measure LD across the whole genome we further thinned the phased data so that no two SNPs were within 500 bp of each other, when the analysis was repeated as above, except that this time each SNP was compared to every other SNP in the entire genome. We also repeated these analyses for sub-sets of parasites within the clades that we identified.

We also analysed the mitochondrial genomes (excluding one individual from site AM due to unexpectedly low mitochondrial read depth), and used Analysis of Molecular Variance (AMOVA) which was conducted in GenAlEx version 6.5 (Peakall and Smouse 2006, 2012). Haplotype maps were generated in PopART version 1.7 (Leigh and Bryant 2015) using the minimum spanning network method (Bandelt *et al*. 1999), and maximum likelihood trees based on unique haplotypes were generated with RaxML version 8.1.15 (Stamatakis 2006), using the general time reversible gamma model of substitution rate heterogeneity, and rapid bootstrapping with 100 replicates was applied. We calculated the proportion of SNPs shared among all pairs or worms and compared this to the nuclear Φ relatedness using a Mantel test.

We also used whole genome sequence data of 10 isofemale lines derived from wild *S. ratti* (**Supplementary Table 4**), which we obtained from the European Variant Archive, study code PRJEB41 https://www.ebi.ac.uk/ena/data/view/PRJEB4163. Among the 90 wild *S. ratti* and 10 isofemale lines there were 235,393 SNPs of which 928 were tri-allelic, the remainder bi-allelic, with a ratio of 1.8 of transitions to transversions.

### Rat population genetic analysis

We only used data for faecal pellets that were successfully genotyped at 6 or more loci, resulting in 132 genotyped faecal pellets. Locus D12Rat42 was excluded from further population genetic analyses due to the low number of rats successfully genotyped at this locus. We used GenAlEx’s (version 6.5; Peakall and Smouse 2006, 2012) pairwise relatedness function) to detect pellets with identical multi-locus genotypes, which we took to have come from the same individual rat. We calculated Ritland and Lynch pairwise relatedness (Lynch and Ritland 1999), where each individual was compared with each other individual, and doubled these values to give a possible range of -1 to 1 from which we calculated the mean within-sample-site and mean among-sample-site relatedness. We determined the log-likelihood of the rat originating from each sampling site using GenAlEx to assign pellet genotypes to each sample site by comparing the multilocus genotype of each rat with the allele frequencies of each of the sampling sites (excluding the rat currently being investigated).

We used Shannon’s mutual information index (^S^H_UA_) to quantify the differences in allele frequencies among sampling sites and to estimate the number of effective migrants per generation. ^S^H_UA_ measures and is valid despite deviations from HWE within subpopulations (Hedrick, 2005; Sherwin *et al.,* 2006). ^S^H_UA_ ranges from 0 (indicating unhindered gene flow) to 1 (indicating a complete lack of gene flow).

We ensured (beyond visual identification) that none of the faecal pellets that we had genotyped were from species other than *R. norvegicus* by seeking to amplify the nine rat microsatellite loci from DNA isolated from other species that may potentially produce contaminating faecal material, specifically that from black rats (*R. rattus*), moles (*Talpa europaea*), and squirrels (*Sciurus carolinensis*). With one exception, none of these microsatellite loci successfully amplified from these species (the exception was locus D12Rat42 that did amplify *R. rattus* DNA) confirming that all successful genotypes were from *R. norvegicus*. As positive controls we used primer pairs (i) Scv1 previously used to amplify *Sciurus* sp. DNA (Hale *et al*., 2001), which successful amplified our *Sciurus* sp. DNA, and weakly amplified *R. norvegicus* DNA, and (ii) RodActin previously used to amplify *R. rattus* DNA (Apte *et al*., 2007), which successfully amplified our *R. rattus* and *R. norvegicus* DNA.

### Parasitism and free-living genes and expansion clusters

We followed previous work that identified “parasitism genes” (Hunt *et al*., 2016). We excluded parasitism genes if they were part of an expansion cluster (below) and the underlying genome assembly was poor. We calculated 95% confidence intervals for percentages from www.sample-size.net.

We define an “expansion cluster” as a genomic region containing four or more genes coding for members of one of three protein families (astacin-like metallopeptidases, CAP domain- containing proteins, or acetylcholinesterases), where there is not more than one other gene between any two genes of those families. This definition differs somewhat from, and is more conservative than that used by (Hunt *et al*., 2016). As controls we used “flanking regions”, which we define as the genomic region directly adjacent to the expansion cluster that is the same size as the cluster itself. Each expansion cluster has two flanking regions.

Lists of genes belonging to these three gene families were collated from Hunt *et al*., 2016 and from this we initially identified 15 expansion clusters. Because clusters 10 and 11 were very close to each other, cluster 10’s right flanking region was shortened to end where expansion cluster 11 began, and expansion cluster 11 was considered to not have a left flanking region. Similarly, to avoid overlap of cluster 11’s right flanking region and cluster 12’s left flanking region we shortened cluster 12’s left flanking region to the start of cluster 11’s right flanking region. Across all expansion clusters there were 135 genes in total, of which 126 belonged to one of the three gene families: 46 encoding CAP domain-containing proteins, 70 encoding astacin-like metallopeptidases, and 10 encoding acetylcholinesterases, representing 51.7%, 38% and 33.3% of CAP domain-containing proteins, astacin-like metallopeptidase and acetylcholinesterase encoding genes, respectively, in the genome as a whole. Flanking regions collectively contained 216 protein- coding genes the products of which had varying predicted functional descriptions.

Mindful that these expansion cluster regions were repetitive in nature we checked their original reference genome assembly by realigning the sequencing reads originally used to build the reference assembly back to the reference, available at NCBI, BioProject code PRJEB2398, and then assessed the quality of these alignments using Gap5 (Bonfield and Whitman 2010). Repetitiveness of the sequence was examined *via* Dotplots with the software package Dotter (Sonnhammer and Durbin, 1995). Gene annotation schematics were retrieved from Ensembl’s ‘Region in Detail’ tool (Hubbard *et al.,* 2002), accessed via WormBase Parasite (Howe *et al.,* 2017) version 12 (https://parasite.wormbase.org/index.html) and added to the graphics produced by Gap5. Regions with poor mapping quality, unusually large distances between mate pairs and the occurrence of mate pairs facing opposite directions are suggestive of high rates of sequence misalignment. Peaks in read depth and fragment depth above background levels were evidence that multiple copies of a repetitive sequence were collapsed in the reference assemblies. Where expansion cluster genes or genes in flanking regions fell in poorly resolved reference assembly areas, these genes were excluded from further analysis.

Using this approach, we excluded expansion clusters 4, 11 and 13, and their flanking regions entirely and other genes within various clusters, resulting in 196 genes remaining, of which 61 were in expansion clusters and 135 were in flanking regions (**Supplementary Table 8**). Expansion clusters 6, 7, 8, 12 and 14 had no genes excluded. Three expansion clusters had genes that did not belong to one of the three target gene families, and were excluded from analyses. Of the remaining 58 in the expansion cluster genes, 29 encoded CAP domain- containing proteins, 27 encoded astacin-like metallopeptidases, and 2 encoded acetylcholinesterases.

## Data Deposition

The genome data of the 90 larvae are deposited in the European Variation Archive, study PRJEB32744, https://www.omicsdi.org/dataset/eva/PRJEB32744.

## Acknowledgements

We would like to thank the land owners for access to the sample sites; Benito Wainwright, Amy Williams Schwartz, and Bristol Zoo Gardens for the provision of mammalian tissue; Andrea Betancourt for substantial help with mutation rate analyses; and Vicky Hunt for helpful discussions. RC was funded by a NERC studentship. The sequence data for this project were produced by the Wellcome Sanger Institute with funding from the Wellcome Trust grant 206194.

## SUPPLEMENTARY INFORMATION

### SUPPLEMENTARY TABLES

1. The occurrence of *S. ratti* in rat faecal pellets.
2. Rat microsatellite loci.
3. Population genetics of rat microsatellite loci.
4. Isofemale lines of *S. ratti* that were whole genome sequenced.
5. Genes in highly variable 10 kb regions.
6. The hundred most parasitic genes.
7. The hundred most free-living genes.
8. *S. ratti* expansion clusters, revised after further inspection of genome assembly in the cluster region.
9. dN/dS ratios of expansion clusters and their flanking regions.
10. Sampling sites and times.

### SUPPLEMENTARY FIGURES

1. Frequency distribution of the number of *S. ratti* infective larvae isolated from infected rat faecal pellets.
2. Histogram of Φ relatedness values among 90 *S. ratti* larvae.
3. The frequency distribution of the pairwise number of SNP differences among the 90 parasites.
4. Maximum likelihood trees of the 90 parasites.
5. ADMIXTURE analysis of the 90 parasites.
6. PCA analysis of *S. ratti* parasites.
7. Minimum spanning *S. ratti* mitochondrial haplotype maps.
8. Ritland and Lynch pairwise relatedness values of rats within sampling sites.
9. *S. ratti* neighbour-joining dendrograms of 10 isofemale lines and 90 larvae collected from the three sample sites.
10. Linkage disequilibrium in the *S. ratti* genome.
11. Linkage disequilibrium in the *S. ratti* genome for clade 1 and 3 parasites.
12. Heatmaps of linkage disequilibrium in the *S. ratti* genome for clade 1 and 3 parasites.
13. Heatmaps of linkage disequilibrium in the *S. ratti* genome.
14. The distribution of SNPs across the *S. ratti* genome.
15. Neighbour-joining dendrograms based on five expansion clusters.
16. Correlation of read depth and GC content for 90 *S. ratti* larvae.

## SUPPLEMENTARY TABLES

**Supplementary Table 1.** The occurrence of *S. ratti* in rat faecal pellets.. The proportion of infected pellets did not differ significantly among the seasons when the faecal pellets were collected (χ^2^ = 6, *df* = 3, P = 0.11).

| Site and season | Number of pellets collected | Number (and %) of infected pellets | Number of larvae collected | Mean (SD) number of larvae per infected pellet |
| --- | --- | --- | --- | --- |
| CA, Spring | 35 | 7 (20%) | 244 | 34.9 (24.3) |
| CA, Summer | 32 | 5 (15.6%) | 89 | 17.8 (22.1) |
| CA, Autumn | 19 | 2 (10.5%) | 256 | 128 (7.1) |
| CA, Winter | 26 | 1 (3.8%) | 6 | 6 (0) |
| **CA, All seasons** | **112** | **15 (13.4%)** | **595** | **39.7 (42.1)** |
| AM, Spring | 11 | 8 (72.7%) | 137 | 17.1 (23.5) |
| AM, Summer | 75 | 23 (30.7%) | 428 | 18.6 (29.3) |
| AM, Autumn | 27 | 15 (55.6%) | 3,044 | 202.9 (185) |
| AM, Winter | 21 | 17 (81%) | 5,067 | 298.1 (500.3) |
| **AM, All seasons** | **134** | **63 (47%)** | **8,676** | **137.7 (296.5)** |
| LA, Spring | 27 | 14 (51.9%) | 211 | 15.1 (23.1) |
| LA, Summer | 11 | 7 (63.6%) | 40 | 5.7 (3.7) |
| LA, Autumn | 7 | 6 (85.7%) | 360 | 60 (46.8) |
| LA, Winter | 13 | 9 (69.2%) | 589 | 65.4 (83.8) |
| **LA, All seasons** | **58** | **36 (62.1%)** | **1,200** | **33.3 (52.8)** |
| All sites, Spring | 73 | 29 (39.7%) | 592 | 20.4 (24.3) |
| All sites, Summer | 118 | 35 (29.7%) | 557 | 15.9 (25) |
| All sites, Autumn | 53 | 23 (43.4%) | 3,660 | 159.1 (162.4) |
| All sites, Winter | 60 | 27 (45%) | 5,662 | 209.7 (412.4) |
| **Total** | **304** | **114 (37.5%)** | **10,471** | **91.9 (226.9)** |

**Supplementary Table 2.**
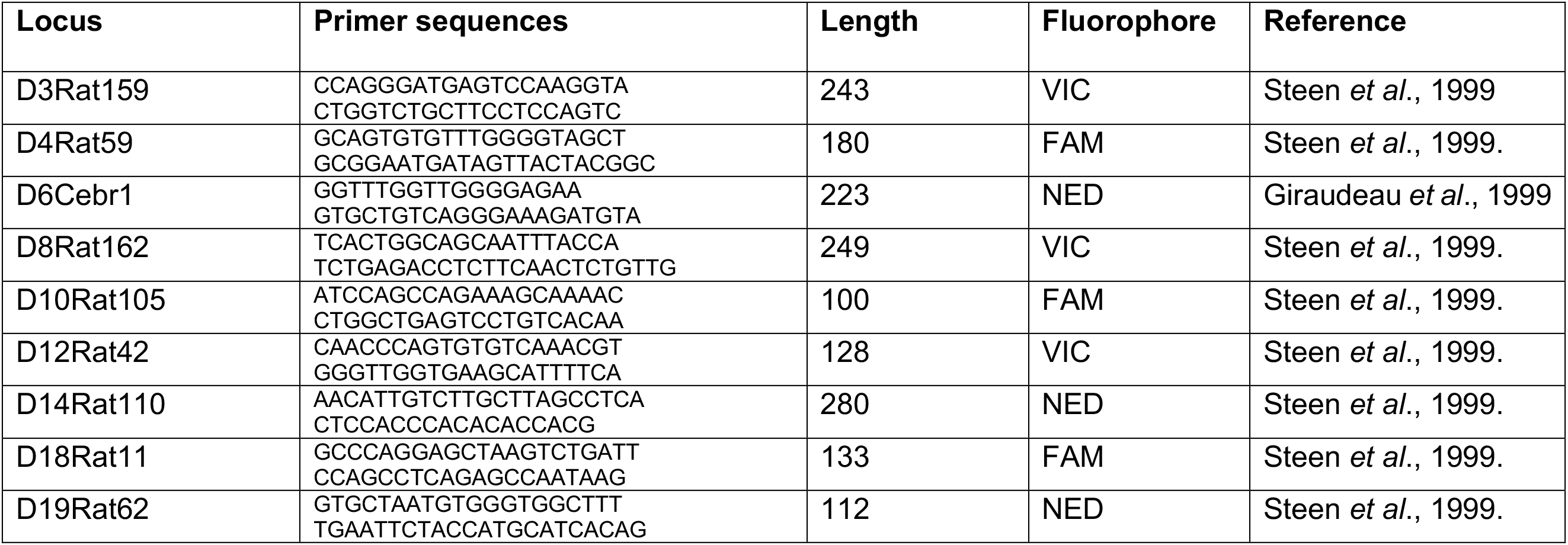
Rat microsatellite loci.. The forward primers are listed first and all primers are 5’ to 3’; Length refers to the length in bp of the region amplified by the given primer sequences as determined from Rnor 6.0 (1.7.2014) release of the *R. norvegicus* genome; Fluorophore indicates the fluorophore used to label forward primers and thus PCR products.

**Supplementary Table 3.** Population genetics of rat microsatellite loci. A. Genetic diversity of loci.. Rats genotyped is the number of individual rats in which genotyping was successful; Allele number is the number of alleles identified at the locus; He and Ho are expected and observed heterozygosity, respectively; Site diversity is the proportion of allelic diversity that partitions within sites, as opposed to among sites, according to ^S^H_UA_. locus D12Rat42 was not used in analyses due to the low genotyping success rate. **B. HWE of loci.** Locus shows the loci; Site is the sampling site; Rats genotyped is the number of rats that were genotyped; Alleles is the number of alleles detected; X^2^ is the statistic testing whether the observed genotype frequencies match HWE expectations with the degrees of freedom (df) in parentheses. Those still significant after Bonferroni correction are shown in bold.

| **A.** |  |  |  |  |  |
| --- | --- | --- | --- | --- | --- |
| **Locus** | **Rats genotyped** | **Allele number** | **He** | **Ho** | **Site diversity (%)** |
| D3Rat159 | 103 | 14 | 0.779 | 0.756 | 18 |
| D4Rat59 | 105 | 12 | 0.822 | 0.652 | 8 |
| D6Cebr1 | 109 | 14 | 0.752 | 0.356 | 13 |
| D8Rat162 | 73 | 15 | 0.835 | 0.600 | 16 |
| D10Rat105 | 112 | 8 | 0.696 | 0.617 | 13 |
| D12Rat42 | 34 | 9 | 0.762 | 0.267 | NA |
| D14Rat110 | 83 | 16 | 0.824 | 0.529 | 18 |
| D18Rat11 | 112 | 10 | 0.686 | 0.319 | 19 |
| D19Rat62 | 96 | 12 | 0.678 | 0.462 | 13 |

| **B.** |  |  |  |  |  |
| --- | --- | --- | --- | --- | --- |
| **Locus** | **Site** | **Rats genotyped** | **Alleles** | **$X^2$ (df)** | **Probability** |
| **D3Rat159** | CA | 45 | 7 | 41.8 (21) | 0.01 |
|  | LA | 17 | 11 | 89.9 (55) | 0.01 |
|  | **AM** | **41** | **10** | **98.4 (45)** | **0.001** |
| **D4Rat59** | **CA** | **46** | **9** | **74.6 (36)** | **0.001** |
|  | LA | 15 | 9 | 39.1 (36) | 0.03 |
|  | AM | 44 | 10 | 67.2 (45) | 0.05 |
| **D6Cebr1** | **CA** | **45** | **11** | **100.7 (55)** | **0.001** |
|  | **LA** | **18** | **10** | **87.6 (45)** | **0.001** |
|  | **AM** | **46** | **6** | **105.1 (15)** | **0.001** |
| **D8Rat162** | **CA** | **25** | **10** | **120.1 (45)** | **0.001** |
|  | LA | 13 | 8 | 28.3 (28) | 0.45 |
|  | AM | 35 | 10 | 67.7 (45) | 0.05 |
| **D10Rat105** | **CA** | **47** | **5** | **89.6 (10)** | **0.001** |
|  | **LA** | **19** | **6** | **64 (15)** | **0.001** |
|  | **AM** | **46** | **7** | **94.2 (21)** | **0.001** |
| **D12Rat42** | **CA** | **15** | **9** | **71.8 (36)** | **0.001** |
|  | LA | 5 | 4 | 12 (6) | 0.06 |
|  | AM | 14 | 6 | 33.1 (15) | 0.01 |
| **D14Rat110** | **CA** | **34** | **11** | **112.4 (55)** | **0.001** |
|  | LA | 16 | 9 | 45.2 (36) | 0.14 |
|  | **AM** | **33** | **11** | **113.6 (55)** | **0.001** |
| **D18Rat11** | **CA** | **47** | **8** | **66.6 (28)** | **0.001** |
|  | **LA** | **19** | **5** | **34.5 (10)** | **0.001** |
|  | **AM** | **46** | **9** | **141 (36)** | **0.001** |
| **D19Rat62** | **CA** | **39** | **9** | **115.2 (36)** | **0.001** |
|  | LA | 17 | 7 | 40.8 (21) | 0.01 |
|  | AM | 40 | 5 | 17.9 (10) | 0.06 |

**Supplementary Table 4.**
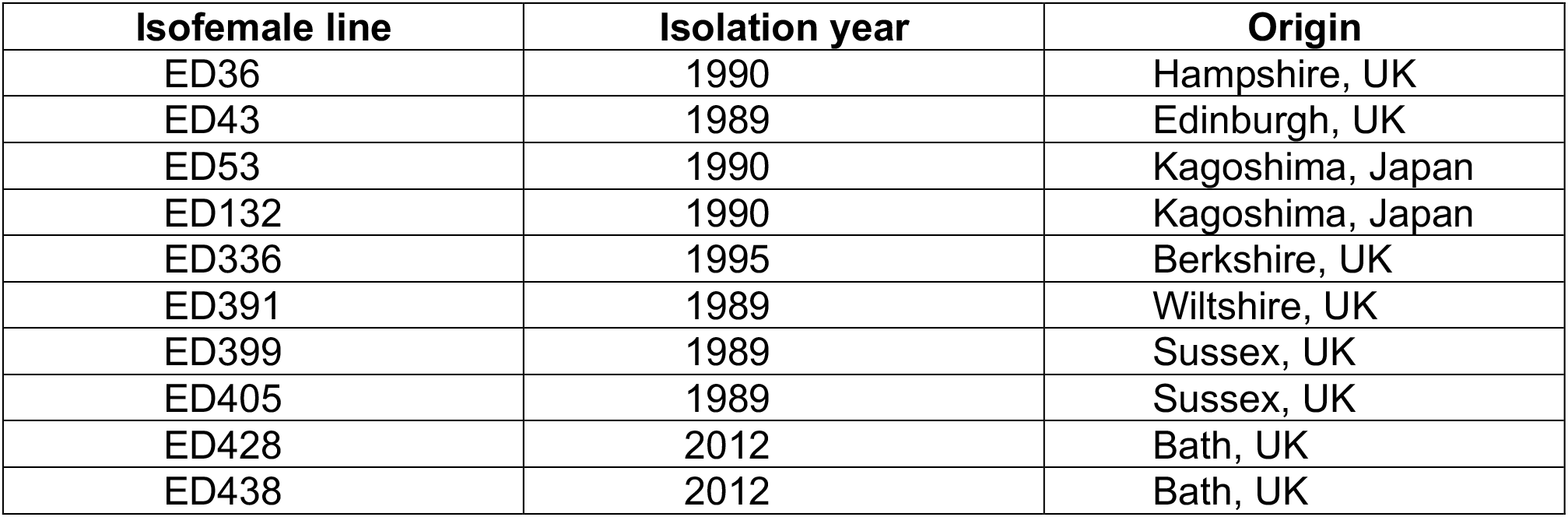
Isofemale lines of *S. ratti* that were whole genome sequenced.

**Supplementary Table 5.** Genes in highly variable 10 kb regions.. Region lists the 61 regions (region 1 does not contain any genes); Gene gives the gene’s designation; Predicted function is the WormBase ParaSite description of each gene, where Astacin-like metalloendopeptidase is abbreviated to Astacin; Coding SNPs per kb is the number of SNPs per kb within the coding sequence of that gene; SNP type is the absolute number of synonymous (S), nonsynonymous (NS) and STOP codon-causing SNPs; Expression is taken from Hunt *et al*., 2016, which compared the expression of genes between the parasitic female and free-living female morph, and where the expression is one log_2_ fold more in the parasitic female morph (Parasitic), free-living female morph (Free-living), not different (Same), or not listed (Unlisted); Expansion cluster is expansion cluster or associated flanking region a gene belongs to, if any. Some genes are marked as “discarded” because they had poor underlying assembly according to Gap5 analysis of expansion clusters and flanking regions and so were discounted from further analysis.

| Region | Gene | Predicted function | Coding SNPs per kb | SNP type (S/NS/STOP) | Expression | Expansion cluster |
| --- | --- | --- | --- | --- | --- | --- |
| 2 | SRAE_0000058700 | Kinesin, motor domain and P-loop-containing | 26.7 | 3 / 3 / 0 | Unlisted | None |
| 3 | SRAE_2000499400 | Hypothetical protein | 48.2 | 5 / 28 / 0 | Parasitic | None |
|  | SRAE_2000499500 | Hypothetical protein | 0 | 0 / 0 / 0 | Parasitic | None |
|  | SRAE_2000499600 | Hypothetical protein | 72.2 | 8 / 20 / 1 | Parasitic | None |
|  | SRAE_2000499700 | Hypothetical protein | 45.7 | 5 / 22 / 0 | Parasitic | None |
| 4 | SRAE_0000058500 | Hypothetical protein | 3.3 | 0 / 1 / 0 | Unlisted | None |
| 5 | SRAE_2000124300 | CAP domain-containing | 69 | 17 / 49 / 4 | Parasitic | EC3 |
|  | SRAE_2000124400 | CAP domain-containing | 43.3 | 8 / 30 / 1 | Parasitic | EC3 |
|  | SRAE_2000124500 | CAP domain-containing | 50.9 | 11 / 33 / 2 | Parasitic | EC3 |
| 6 | SRAE_X000232600 | Reverse transcriptase domain and Aspartic peptidase domain-containing | 45.8 | 62 / 51 / 1 | Unlisted | None |
|  | SRAE_X000232700 | Integrase, catalytic core domain and Ribonuclease H-like domain-containing | 22.2 | 5 / 1 / 1 | Unlisted | None |
|  | SRAE_X000232800 | Hypothetical protein | 32 | 6 / 6 / 0 | Same | None |
|  | SRAE_X000232900 | Hypothetical protein | 31.7 | 4 / 4 / 0 | Unlisted | None |
|  | SRAE_X000233000 | Zinc finger, CCHC-type domain-containing | 11.8 | 14 / 36 / 0 | Unlisted | None |
|  | SRAE_X000233100 | Hypothetical protein | 11.3 | 12 / 22 / 1 | Unlisted | None |
| 7 | SRAE_0000057800 | Hypothetical protein | 28.5 | 1 / 9 / 0 | Parasitic | None |
| 8 | SRAE_X000062200<br>(discarded) | Poly-glutamine tract binding protein 1 | 0 | 0 / 0 / 0 | Unlisted | EC13 |
|  | SRAE_X000062300<br>(discarded) | Acetylcholinesterase | 66.2 | 29 / 86 / 0 | Same | EC13 |
| 9 | SRAE_2000478200 | Hypothetical protein | 6.2 | 4 / 0 / 0 | Unlisted | None |
|  | SRAE_2000478300 | Hypothetical protein | 21.1 | 5 / 6 / 0 | Same | None |
|  | SRAE_2000478400 | Hypothetical protein | 90.7 | 10 / 29 / 1 | Unlisted | None |
|  | SRAE_2000478500 | CAP domain-containing | 89.3 | 17 / 47 / 0 | Parasitic | None |
| 10 | SRAE_X000050900 | Hypothetical protein | 66.7 | 13 / 34 / 0 | Parasitic | None |
|  | SRAE_X000051000 | Hypothetical protein | 44.5 | 8 / 21 / 0 | Parasitic | None |
|  | SRAE_X000051100 | Hypothetical protein | 53 | 6 / 27 / 1 | Parasitic | None |
| 12 | SRAE_X000026900 | Hypothetical protein | 51 | 5 / 19 / 0 | Parasitic | None |
|  | SRAE_X000027000 | Hypothetical protein | 15.8 | 3 / 4 / 0 | Unlisted | None |
| 13 | SRAE_1000139800 | Hypothetical protein | 8 | 12 / 2 / 0 | Unlisted | None |
|  | SRAE_1000139900 | Protein-tyrosine phosphatase-containing | 54.6 | 42 / 145 / 1 | Same | None |
|  | SRAE_1000140000 | Translation initiation factor SUI1 domain-containing | 0 | 0 / 0 / 0 | Same | None |
|  | SRAE_1000140100 | Hypothetical protein | 0 | 0 / 0 / 0 | Same | None |
|  | SRAE_1000140200 | Hypothetical protein | 18.1 | 2 / 8 / 0 | Same | None |
| 14 | SRAE_2000076600 | CAP domain-containing | 21.2 | 3 / 14 / 0 | Parasitic | EC2 |
|  | SRAE_2000076700 | CAP domain-containing | 60.4 | 11 / 39 / 0 | Parasitic | EC2 |
|  | SRAE_2000076800 | CAP domain-containing | 39.2 | 11 / 18 / 1 | Parasitic | EC2 |
|  | SRAE_2000076900 | CAP domain-containing | 36.8 | 6 / 25 / 0 | Parasitic | EC2 |
|  | SRAE_2000077000 | CAP domain-containing | 19 | 6 / 9 / 1 | Parasitic | EC2 |
| 15 | SRAE_2000499000 | Hypothetical protein | 55.9 | 2 / 25 / 1 | Parasitic | None |
|  | SRAE_2000499100 | Hypothetical protein | 35.8 | 7 / 12 / 0 | Parasitic | None |
| 16 | SRAE_X000037500 | Hypothetical protein | 21.7 | 8 / 18 / 1 | Unlisted | None |
| 17 | SRAE_X000066000 | Trypsin Inhibitor-like | 36.3 | 38 / 77 / 0 | Parasitic | None |
|  | SRAE_X000066100 | Astacin | 28.2 | 18 / 23 / 0 | Parasitic | None |
| 18 | SRAE_X000032850 | Hypothetical protein | 69.2 | 4 / 18 / 0 | Unlisted | None |
|  | SRAE_X000032900 | Hypothetical protein | 54.6 | 8 / 11 / 0 | Unlisted | None |
|  | SRAE_X000033000 | Hypothetical protein | 78.6 | 9 / 20 / 0 | Unlisted | None |
|  | SRAE_X000033100 | Hypothetical protein | 46 | 4 / 12 / 0 | Unlisted | None |
| 19 | SRAE_X000050700 | Hypothetical protein | 34.9 | 9 / 15 / 0 | Parasitic | None |
|  | SRAE_X000050800 | Hypothetical protein | 37 | 7 / 16 / 0 | Parasitic | None |
| 20 | SRAE_0000074500 | Aspartic peptidase domain-containing | 46.4 | 93 / 116 / 6 | Free-living | None |
|  | SRAE_0000074600 | Aspartic peptidase domain-containing | 9.5 | 15 / 19 / 1 | Unlisted | None |
| 21 | SRAE_X000145900 | Astacin | 82.7 | 27 / 76 / 1 | Same | None |
|  | SRAE_X000146000 | Astacin | 21.4 | 7 / 19 / 1 | Para | None |
| 22 | SRAE_X000186500 | Reverse transcriptase domain-containing | 25.3 | 37 / 65 / 2 | Free-living | None |
|  | SRAE_X000186600 | Cytochrome C oxidase subunit II | 43.6 | 14 / 51 / 0 | Same | None |
| 23 | SRAE_X000146300<br>(discarded) | Hypothetical protein | 31.3 | 5 / 5 / 1 | Unlisted | FR15L |
|  | SRAE_X000146400<br>(discarded) | Hypothetical protein | 6.4 | 2 / 1 / 0 | Unlisted | FR15L |
| 24 | SRAE_0000074000 | Hypothetical protein | 16.7 | 3 / 6 / 1 | Unlisted | None |
|  | SRAE_0000074100 | Hypothetical protein | 10.4 | 2 / 6 / 0 | Unlisted | None |
|  | SRAE_0000074200 | Aspartic peptidase domain-containing | 42 | 86 / 74 / 0 | Same | None |
| 25 | SRAE_2000460700 | Hypothetical protein | 52.4 | 20 / 112 / 1 | Parasitic | None |
|  | SRAE_2000460800 | Hypothetical protein | 4.6 | 1 / 1 / 0 | Parasitic | None |
|  | SRAE_2000460900 | Hypothetical protein | 6.7 | 1 / 3 / 1 | Same | None |
| 26 | SRAE_X000063000<br>(discarded) | Acetylcholinesterase | 15.2 | 9 / 18 / 0 | Parasitic | EC13 |
|  | SRAE_X000063100<br>(discarded) | Poly-glutamine tract binding protein 1 | 0.4 | 0 / 1 / 0 | Same | FR13R |
| 27 | SRAE_1000045700 | E3 ubiquitin-protein ligase MYCBP2 | 4.9 | 51 / 20 / 0 | Free-living | None |
|  | SRAE_1000045800 | Lipase, class 3 family-containing | 15.2 | 7 / 8 / 0 | Parasitic | None |
|  | SRAE_1000045900 | CAP domain-containing | 76 | 19 / 67 / 0 | Parasitic | None |
|  | SRAE_1000046000 | Speckle-type POZ | 4.2 | 4 / 1 / 0 | Same | None |
| 28 | SRAE_X000051500 | CTP synthase 2 | 26.6 | 5 / 23 / 1 | Unlisted | None |
|  | SRAE_X000051600 | CTP synthase 2 | 62.4 | 6 / 26 / 0 | Unlisted | None |
|  | SRAE_X000051700 | Hypothetical protein | 9.1 | 1 / 3 / 0 | Parasitic | None |
|  | SRAE_X000051800 | CTP synthase 2 | 6,4 | 2 / 2 / 1 | Unlisted | None |
| 29 | SRAE_0000058000 | Reverse transcriptase domain-containing | 40.7 | 19 / 46 / 0 | Unlisted | None |
|  | SRAE_0000058100 | Integrase | 38.9 | 18 / 21 / 3 | Unlisted | None |
|  | SRAE_0000058200 | Hypothetical protein | 26.5 | 3 / 3 / 1 | Unlisted | None |
|  | SRAE_0000058300 | Reverse transcriptase domain-containing | 3.3 | 2 / 8 / 0 | Unlisted | None |
| 30 | SRAE_X000146200<br>(discarded) | Hypothetical protein | 23.2 | 4 / 6 / 1 | Unlisted | FR15L |
| 31 | SRAE_0000049800 | Transposase, ISXO2-like domain-containing | 24.9 | 2 / 6 / 2 | Unlisted | None |
|  | SRAE_0000049900 | Integrase | 9.6 | 4 / 20 / 2 | Unlisted | None |
| 32 | SRAE_X000088600 | Ras-like protein 3 | 0 | 0 / 0 / 0 | Free-living | None |
| 33 | SRAE_X000195800 | Hypothetical protein | 0 | 0 / 0 / 0 | Parasitic | None |
|  | SRAE_X000195900 | Hypothetical protein | 30.7 | 2 / 6 / 0 | Parasitic | None |
|  | SRAE_X000196000 | Hypothetical protein | 12.4 | 1 / 6 / 0 | Parasitic | None |
|  | SRAE_X000196100 | Hypothetical protein | 31.1 | 2 / 4 / 1 | Unlisted | None |
| 34 | SRAE_0000025500 | Plasma membrane calcium-transporting ATPase 3 | 2.2 | 4 / 5 / 0 | Free-living | None |
|  | SRAE_0000025600 | Transthyretin-like family-containing | 103.9 | 11 / 37 / 0 | Parasitic | None |
|  | SRAE_0000025650 | Hypothetical protein | 26.3 | 2 / 9 / 1 | Unlisted | None |
|  | SRAE_0000025700 | Transthyretin-like family-containing | 0 | 0 / 0 / 0 | Parasitic | None |
|  | SRAE_0000025800 | Protein argonaute-4 | 4.9 | 5 / 9 / 0 | Parasitic | None |
| 35 | SRAE_0000007600 | Amino acid transporter | 2.5 | 4 / 0 / 0 | Free-living | None |
|  | SRAE_0000007700 | Cathepsin L.1 | 13.4 | 9 / 4 / 0 | Unlisted | None |
|  | SRAE_0000007800 | Cathepsin L.1 | 8.2 | 3 / 5 / 0 | Free-living | None |
|  | SRAE_0000007900 | Hypothetical protein | 29.5 | 14 / 20 / 0 | Parasitic | None |
|  | SRAE_0000008000 | Hypothetical protein | 37.5 | 10 / 32 / 1 | Parasitic | None |
| 36 | SRAE_X000138600 | Hypothetical protein | 2.2 | 0 / 1 / 0 | Same | None |
|  | SRAE_X000138700 | MAM domain and Concanavalin A-like lectin | 2.3 | 4 / 0 / 0 | Same | None |
| 37 | SRAE_X000236650 | Hypothetical protein | 24.4 | 3 / 18 / 1 | Unlisted | None |
|  | SRAE_X000236700 | Ribonuclease H-like domain and AT hook-like family-containing | 10 | 2 / 7 / 0 | Unlisted | None |
|  | SRAE_X000236800 | Hypothetical protein | 18.3 | 1 / 8 / 1 | Unlisted | None |
|  | SRAE_X000236900 | Aspartic peptidase domain-containing | 13.9 | 10 / 25 / 0 | Unlisted | None |
|  | SRAE_X000237000 | Hypothetical protein | 26.5 | 8 / 1 / 1 | Unlisted | None |
|  | SRAE_X000237100 | Hypothetical protein | 29.2 | 3 / 7 / 0 | Unlisted | None |
| 38 | SRAE_0000057900 | Hypothetical protein | 42.9 | 3 / 10 / 0 | Parasitic | None |
| 39 | SRAE_0000008000 | Hypothetical protein | 37.5 | 10 / 32 / 1 | Parasitic | None |
|  | SRAE_0000008100 | Sulfotransferase family-containing | 19.8 | 5 / 20 / 0 | Parasitic | None |
|  | SRAE_0000008200 | Astacin-like metalloendopeptidase | 4.3 | 4 / 1 / 0 | Parasitic | None |
| 40 | SRAE_2000478600 | CAP domain-containing | 90 | 12 / 51 / 1 | Parasitic | None |
|  | SRAE_2000478610 | Hypothetical protein | 7.5 | 1 / 3 / 0 | Unlisted | None |
|  | SRAE_2000478700 | Zinc finger, BED-type predicted domain-containing | 8 | 8 / 8 / 0 | Free-living | None |
| 41 | SRAE_2000499200 | Hypothetical protein | 75.7 | 8 / 27 / 2 | Parasitic | None |
|  | SRAE_2000499210 | Hypothetical protein | 40.7 | 5 / 17 / 0 | Parasitic | None |
|  | SRAE_2000499300 | Hypothetical protein | 15.5 | 2 / 6 / 0 | Parasitic | None |
| 42 | SRAE_X000104500 | Hypothetical protein | 22.2 | 30 / 81 / 0 | Parasitic | None |
| 43 | SRAE_X000038200 | Hypothetical protein | 25.2 | 8 / 5 / 0 | Unlisted | None |
|  | SRAE_X000038300 | Synaptogyrin | 3.1 | 2 / 1 / 0 | Free-living | None |
|  | SRAE_X000038400 | EF-hand domain | 2.9 | 2 / 0 / 0 | Unlisted | None |
|  | SRAE_X000038500 | RUN domain-containing | 4.9 | 4 / 5 / 0 | Parasitic | None |
| 44 | SRAE_X000246300 | Hypothetical protein | 0 | 0 / 0 / 0 | Same | None |
| 45 | SRAE_X000222600 | Hypothetical protein | 30.8 | 1 / 10 / 0 | Parasitic | None |
| 46 | SRAE_X000158400 | Astacin-like metalloendopeptidase | 1.4 | 0 / 2 / 0 | Free-living | None |
|  | SRAE_X000158500 | Acetylcholinesterase | 8.7 | 1 / 14 / 0 | Parasitic | None |
| 47 | SRAE_2000420700 | CAP domain-containing | 34.1 | 5 / 28 / 0 | Parasitic | None |
|  | SRAE_2000420800 | Hypothetical protein | 11.1 | 0 / 4 / 0 | Unlisted | None |
| 48 | SRAE_X000221400 | U1 small nuclear ribonucleoprotein<br>70 kDa | 0 | 0 / 0 / 0 | Same | None |
|  | SRAE_X000221500 | Hypothetical protein | 5.8 | 0 / 2 / 0 | Unlisted | None |
|  | SRAE_X000221600 | Hypothetical protein | 46.4 | 7 / 9 / 0 | Unlisted | None |
| 49 | SRAE_X000062400<br>(discarded) | Acetylcholinesterase | 24.9 | 14 / 28 / 1 | same | EC13 |
| 50 | SRAE_X000201100 | Astacin | 29.8 | 17 / 27 / 1 | Parasitic | None |
|  | SRAE_X000201200 | Hypothetical protein | 12 | 1 / 5 / 0 | Unlisted | None |
|  | SRAE_X000201300 | Hypothetical protein | 0 | 0 / 0 / 0 | Unlisted | None |
| 51 | SRAE_X000247200 | Carboxylesterase | 6.1 | 1 / 13 / 0 | Free-living | None |
|  | SRAE_X000247300 | ShKT domain-containing | 41.8 | 28 / 100 / 0 | Parasitic | None |
| 52 | SRAE_1000127500 | Amino acid transporter | 48.2 | 41 / 34 / 0 | Same | None |
|  | SRAE_1000127600 | Farnesyltransferase, CAAX box,<br>beta | 14 | 34 / 16 / 0 | Same | None |
|  | SRAE_1000127700 | Hypothetical protein | 2.6 | 1 / 1 / 0 | Unlisted | None |
| 53 | SRAE_X000027100 | Hypothetical protein | 16.4 | 2 / 6 / 0 | Parasitic | None |
|  | SRAE_X000027200 | Hypothetical protein | 38.3 | 13 / 13 / 1 | Parasitic | None |
|  | SRAE_X000027300 | Hypothetical protein | 8.2 | 2 / 2 / 0 | Parasitic | None |
| 54 | SRAE_X000144200 | Astacin | 39.5 | 24 / 24 / 1 | Same | EC14 |
|  | SRAE_X000144210 | Astacin | 21.5 | 6 / 21 / 0 | Parasitic | EC14 |
|  | SRAE_X000144220 | Hypothetical protein | 9.4 | 4 / 4 / 0 | Same | FR14R |
| 55 | SRAE_X000199600 | Hypothetical protein | 11.8 | 2 / 9 / 0 | Unlisted | None |
|  | SRAE_X000199700 | Integrase | 8.7 | 5 / 11 / 0 | Unlisted | None |
|  | SRAE_X000199800 | Hypothetical protein | 19.5 | 2 / 5 / 1 | Unlisted | None |
|  | SRAE_X000199900 | Zinc finger, CCHC-type domain-containing | 24.5 | 9 / 14 / 0 | Unlisted | None |
|  | SRAE_X000200000 | Reverse transcriptase domain-containing | 38.4 | 25 / 20 / 1 | Unlisted | None |
|  | SRAE_X000200100 | Hypothetical protein | 34.2 | 22 / 16 / 1 | Unlisted | None |
| 56 | SRAE_1000059600 | Ground-like domain-containing | 22.8 | 22 / 50 / 0 | Same | None |
|  | SRAE_1000059700 | Phosphate-regulating neutral endopeptidase | 14.6 | 10 / 13 / 0 | Parasitic | None |
|  | SRAE_1000059800 | Phosphate-regulating neutral endopeptidase | 11.3 | 6 / 10 / 0 | Parasitic | None |
|  | SRAE_1000059900 | Phosphate-regulating neutral endopeptidase | 12.5 | 9 / 9 / 0 | Same | None |
| 57 | SRAE_X000168300 | Hypothetical protein | 46.3 | 8 / 22 / 0 | Parasitic | None |
|  | SRAE_X000168400 | Hypothetical protein | 23.8 | 2 / 7 / 0 | Parasitic | None |
| 58 | SRAE_X000200200 | Hypothetical protein | 3.2 | 0 / 1 / 0 | Unlisted | None |
|  | SRAE_X000200300 | Hypothetical protein | 2 | 1 / 0 / 0 | Parasitic | None |
| 59 | SRAE_0000065600 | Hypothetical protein | 69.8 | 28 / 30 / 3 | Unlisted | None |
|  | SRAE_0000065700 | Hypothetical protein | 24.8 | 3 / 6 / 0 | Unlisted | None |
|  | SRAE_0000065800 | Hypothetical protein | 12.6 | 1 / 3 / 0 | Unlisted | None |
|  | SRAE_0000065900 | Ribonuclease H-like domain-containing protein | 6.4 | 2 / 2 / 0 | Unlisted | None |
| 60 | SRAE_2000325900<br>(discarded) | Astacin | 25.3 | 13 / 20 / 0 | Parasitic | EC5 |
|  | SRAE_2000326000 | Astacin | 7.6 | 3 / 7 / 0 | Parasitic | EC5 |
| 61 | SRAE_X000233200 | Hypothetical protein | 11.8 | 6 / 14 / 1 | Unlisted | None |
|  | SRAE_X000233300 | Hypothetical protein | 11.3 | 2 / 3 / 0 | Unlisted | None |
|  | SRAE_X000233400 | Hypothetical protein | 23.8 | 3 / 14 / 0 | Unlisted | None |
|  | SRAE_X000233500 | Reverse transcriptase domain-containing | 23.7 | 10 / 31 / 4 | Unlisted | None |
|  | SRAE_X000233600 | Hypothetical protein | 30.5 | 9 / 37 / 1 | Unlisted | None |
| 62 | SRAE_2000124600 | CAP domain-containing | 8.7 | 3 / 5 / 0 | Parasitic | EC3 |
|  | SRAE_2000124700 | CAP domain-containing | 5.2 | 0 / 4 / 1 | Parasitic | EC3 |
|  | SRAE_2000124800 | CAP domain-containing | 6.5 | 4 / 2 / 0 | Parasitic | EC3 |
|  | SRAE_2000124900 | CAP domain-containing | 6.6 | 2 / 4 / 0 | Parasitic | EC3 |
|  | SRAE_2000125000<br>(discarded) | CAP domain-containing | 2.2 | 1 / 1 / 0 | Parasitic | EC3 |
|  | SRAE_2000125100<br>(discarded) | CAP domain-containing | 38.6 | 9 / 25 / 2 | Parasitic | EC3 |

**Supplementary Table 6.** The hundred most parasitic genes.. Gene gives the gene’s designation; Predicted function is the WormBase ParaSite description of each gene, where Astacin-like metalloendopeptidase is abbreviated to Astacin; Coding SNPs per kb is the number of SNPs per kb within the coding sequence of that gene; SNP type is the absolute number of synonymous (S), nonsynonymous (NS) and STOP codon-causing SNPs; Fold change is the log_2_ difference in expression of the gene between the parasitic female and free-living female morphs taken from Hunt *et al.,* 2016, here shown as positive values indicating greater expression in the parasitic female morph; Expansion cluster is expansion cluster or associated flanking region a gene belongs to, if any; Variable region is as defined in **Supplementary** Table 5. Some genes are marked as “discarded”, because they had poor underlying assembly according to Gap5 analysis of expansion clusters and flanking regions and so were discounted from analyses.

| Gene | Predicted function | Coding SNPs per kb | SNP type (S/NS/STOP) | Fold Change | Expansion cluster | Variable region |
| --- | --- | --- | --- | --- | --- | --- |
| SRAE_2000436100 | Hypothetical protein | 4.7 | 1 / 2 / 0 | 14.5 | None | None |
| SRAE_X000014200 | Hypothetical protein | 15.2 | 0 / 6 / 1 | 13.4 | None | None |
| SRAE_X000201000 | Hypothetical protein | 1.9 | 0 / 1 / 0 | 13 | None | 50 |
| SRAE_1000182300 | CAP domain-containing | 0 | 0 / 0 / 0 | 12.9 | EC1 | None |
| SRAE_2000498800 | Phloem filament PP1 domain-containing | 0 | 0 / 0 / 0 | 12.7 | None | None |
| SRAE_2000067500 | Transthyretin-like family-containing | 15.9 | 1 / 6 / 0 | 12.6 | None | None |
| SRAE_X000201200 | Hypothetical protein | 12 | 1 / 5 / 0 | 12.1 | None | None |
| SRAE_1000182700 (discarded) | CAP domain-containing | 0 | 0 / 0 / 0 | 12.1 | EC1 | None |
| SRAE_2000420000 | Astacin | 1.7 | 0 / 1 / 1 | 12 | None | None |
| SRAE_X000200300 | Hypothetical protein | 2 | 1 / 0 / 0 | 11.8 | None | None |
| SRAE_X000124900 | Hypothetical protein | 0 | 0 / 0 / 0 | 11.8 | None | None |
| SRAE_2000485600 | Tissue inhibitor of metalloproteinase family | 15 | 3 / 3 / 1 | 11.8 | None | None |
| SRAE_X000222400 | Hypothetical protein | 73.8 | 7 / 20 / 0 | 11.7 | None | 45 |
| SRAE_X000055800 | Hypothetical protein | 54.3 | 3 / 12 / 0 | 11.7 | None | None |
| SRAE_2000498700 | Hypothetical protein | 27.8 | 4 / 9 / 0 | 11.7 | None | None |
| SRAE_X000124800 | Hypothetical protein | 0 | 0 / 0 / 0 | 11.6 | None | None |
| SRAE_X000037700 | Hypothetical protein | 0 | 0 / 0 / 0 | 11.6 | None | None |
| SRAE_0000057900 | Hypothetical protein | 42.9 | 3 / 10 / 0 | 11.5 | None | 38 |
| SRAE_2000457510 | CAP domain-containing | 13.8 | 4 / 6 / 0 | 11.4 | None | None |
| SRAE_1000045800 | Lipase, class 3 family-containing | 15.2 | 7 / 8 / 0 | 11.4 | None | 27 |
| SRAE_2000506800 | Hypothetical protein | 14.7 | 8 / 27 / 0 | 11.3 | None | None |
| SRAE_0000045600 | CAP domain-containing protein | 9.5 | 3 / 4 / 0 | 11.3 | None | None |
| SRAE_2000453600 | Astacin | 0 | 0 / 0 / 0 | 11.3 | EC7 | None |
| SRAE_2000522700 | Trypsin Inhibitor-like | 9 | 3 / 3 / 0 | 11.2 | None | None |
| SRAE_0000071120 | Metalloendopeptidase | 1 | 0 / 1 / 0 | 11.2 | None | None |
| SRAE_2000522300 | Purple acid phosphatase | 4.8 | 2 / 5 / 0 | 11.2 | None | None |
| SRAE_2000465200 | Hypothetical protein | 0 | 0 / 0 / 0 | 11.1 | None | None |
| SRAE_X000246200 | Hypothetical protein | 0 | 0 / 0 / 0 | 11.1 | None | None |
| SRAE_2000475600 | Hypothetical protein | 34.9 | 3 / 10 / 0 | 11.1 | None | None |
| SRAE_2000077400 | CAP domain-containing | 13 | 4 / 7 / 0 | 11.1 | EC2 | None |
| SRAE_2000465000 | Hypothetical protein | 0 | 0 / 0 / 0 | 11.1 | None | None |
| SRAE_X000168300 | Hypothetical protein | 46.3 | 8 / 22 / 0 | 11 | None | 57 |
| SRAE_2000499910 | Hypothetical protein | 7.5 | 2 / 2 / 0 | 11 | None | None |
| SRAE_2000499810 | Hypothetical protein | 41.4 | 4 / 17 / 0 | 11 | None | None |
| SRAE_0000057800 | Hypothetical protein | 28.5 | 1 / 9 / 0 | 11 | None | 7 |
| SRAE_0000077300 | Trypsin Inhibitor-like | 6.3 | 1 / 1 / 1 | 11 | None | None |
| SRAE_1000296600 | Hypothetical protein | 0 | 0 / 0 / 0 | 10.9 | None | None |
| SRAE_2000499820 | Hypothetical protein | 1.6 | 1 / 0 / 0 | 10.9 | None | None |
| SRAE_2000522600 | Trypsin Inhibitor-like | 25.6 | 10 / 7 / 0 | 10.9 | None | None |
| SRAE_X000144210 | Astacin | 21.5 | 6 / 21 / 0 | 10.9 | EC14 | 54 |
| SRAE_2000486100 | Tissue inhibitor of metalloproteinase family | 51.9 | 7 / 17 / 0 | 10.9 | None | None |
| SRAE_X000168800 | Prolyl endopeptidase | 0 | 0 / 0 / 0 | 10.9 | None | None |
| SRAE_2000453500 | Astacin | 2.5 | 0 / 3 / 0 | 10.8 | EC7 | None |
| SRAE_2000485800 | Tissue inhibitor of metalloproteinase family | 0 | 0 / 0 / 0 | 10.8 | None | None |
| SRAE_2000499500 | Hypothetical protein | 0 | 0 / 0 / 0 | 10.7 | None | 3 |
| SRAE_X000065700 | Prolyl endopeptidase | 3.1 | 4 / 3 / 0 | 10.7 | None | None |
| SRAE_2000461200 | Hypothetical protein | 25.2 | 3 / 13 / 0 | 10.7 | None | None |
| SRAE_X000051700 | Hypothetical protein | 9.1 | 1 / 3 / 0 | 10.6 | None | 28 |
| SRAE_2000456500 | Metalloendopeptidase | 2.3 | 2 / 1 / 0 | 10.6 | None | None |
| SRAE_1000182400 | CAP domain-containing | 1.2 | 1 / 0 / 0 | 10.6 | EC1 | None |
| SRAE_2000460300 | Zinc metalloproteinase | 17.1 | 5 / 19 / 0 | 10.6 | None | None |
| SRAE_2000525700<br>(discarded) | Astacin | 29.4 | 3 / 22 / 0 | 10.6 | EC11 | None |
| SRAE_X000201100 | Astacin | 29.8 | 17 / 27 / 1 | 10.5 | None | 50 |
| SRAE_0000008000 | Hypothetical protein | 37.5 | 10 / 32 / 0 | 10.5 | None | 35 |
| SRAE_X000038800 | Hypothetical protein | 0 | 0 / 0 / 0 | 10.4 | None | None |
| SRAE_2000076800 | CAP domain-containing | 39.2 | 11 / 18 / 1 | 10.4 | EC2 | 14 |
| SRAE_2000515500 | Hypothetical protein | 2.9 | 9 / 15 / 0 | 10.4 | None | None |
| SRAE_X000055700 | Hypothetical protein | 0 | 0 / 0 / 0 | 10.4 | None | None |
| SRAE_1000182600<br>(discarded) | CAP domain-containing | 17.8 | 2 / 14 / 0 | 10.4 | EC1 | None |
| SRAE_X000168400 | Astacin | 23.8 | 2 / 7 / 0 | 10.3 | None | None |
| SRAE_X000169100 | Prolyl endopeptidase | 0 | 0 / 0 / 0 | 10.3 | None | None |
| SRAE_0000081000 | Metalloendopeptidase | 0 | 0 / 0 / 0 | 10.3 | None | None |
| SRAE_0000000600 | Hypothetical protein | 0 | 0 / 0 / 0 | 10.3 | None | None |
| SRAE_X000195900 | Hypothetical protein | 30.7 | 2 / 6 / 0 | 10.3 | None | 58 |
| SRAE_2000523800 | Astacin | 2.8 | 2 / 2 / 0 | 10.3 | EC10 | None |
| SRAE_1000183100<br>(discarded) | CAP domain-containing | 20.4 | 3 / 14 / 1 | 10.3 | EC1 | None |
| SRAE_2000077000 | CAP domain-containing | 19 | 6 / 9 / 1 | 10.2 | EC2 | 14 |
| SRAE_X000226400 | CAP domain-containing | 2 | 1 / 1 / 0 | 10.2 | None | None |
| SRAE_X000066100 | Astacin | 28.2 | 18 / 23 / 0 | 10.2 | None | 17 |
| SRAE_X000055500 | Hypothetical protein | 0 | 0 / 0 / 0 | 10.2 | None | None |
| SRAE_2000499920 | Hypothetical protein | 85.9 | 14 / 52 / 0 | 10.2 | None | None |
| SRAE_X000191900 | Hypothetical protein | 0 | 0 / 0 / 0 | 10.1 | None | 33 |
| SRAE_X000222600 | Hypothetical protein | 30.8 | 1 / 10 / 0 | 10.1 | None | None |
| SRAE_2000451600 | Transthyretin-like family-containing | 20.6 | 2 / 8 / 0 | 10.1 | None | None |
| SRAE_X000191800 | Hypothetical protein | 0 | 0 / 0 / 0 | 10.1 | None | None |
| SRAE_2000455000 | Astacin | 2.6 | 1 / 2 / 0 | 10.1 | EC8 | None |
| SRAE_0000078700 | Hypothetical protein | 0 | 0 / 0 / 0 | 10 | None | None |
| SRAE_2000455300 | Acetylcholinesterase | 0 | 0 / 0 / 0 | 10 | EC8 | None |
| SRAE_X000200700 | Hypothetical protein | 57.9 | 3 / 26 / 0 | 10 | None | None |
| SRAE_2000499200 | Hypothetical protein | 75.7 | 8 / 27 / 2 | 10 | None | 41 |
| SRAE_2000071300 | Hypothetical protein | 30.3 | 8 / 17 / 0 | 10 | None | None |
| SRAE_X000158300 | Acetylcholinesterase | 3 | 3 / 2 / 0 | 10 | None | 57 |
| SRAE_2000124300 | CAP domain-containing protein | 69 | 17 / 49 / 4 | 10 | EC3 | 5 |
| SRAE_2000457710 | Metallopeptidase, catalytic domain-containing | 1.8 | 2 / 0 / 0 | 10 | None | None |
| SRAE_2000527500 | CAP domain-containing | 3.3 | 3 / 0 / 0 | 10 | EC12 | None |
| SRAE_2000453700 | Astacin | 5.5 | 0 / 7 / 0 | 9.9 | EC7 | None |
| SRAE_2000325600 | Astacin | 0.9 | 1 / 0 / 0 | 9.9 | EC5 | None |
| SRAE_2000525800<br>(discarded) | Astacin | 1.4 | 0 / 1 / 0 | 9.9 | EC11 | None |
| SRAE_X000055600 | Hypothetical protein | 0 | 0 / 0 / 0 | 9.9 | None | None |
| SRAE_2000482710 | Zinc metalloproteinase | 17.8 | 6 / 19 / 0 | 9.8 | None | None |
| SRAE_0000077800 | Hypothetical protein | 4.6 | 0 / 1 / 0 | 9.8 | None | None |
| SRAE_2000489900 | Aspartic peptidase family | 0.9 | 1 / 0 / 0 | 9.8 | None | None |
| SRAE_2000289800<br>(discarded) | Astacin | 0 | 0 / 0 / 0 | 9.8 | EC11 | None |
| SRAE_2000508100 | Hypothetical protein | 66 | 4 / 14 / 2 | 9.8 | None | None |
| SRAE_2000456300 | Acetylcholinesterase | 5.2 | 1 / 8 / 0 | 9.7 | None | None |
| SRAE_0000071100 | Metalloendopeptidase | 0 | 0 / 0 / 0 | 9.7 | None | None |
| SRAE_0000082200 | Trypsin Inhibitor-like | 18.1 | 6 / 19 / 0 | 9.7 | None | None |
| SRAE_2000126100 | Hypothetical protein | 8.6 | 6 / 2 / 0 | 9.7 | FR3R | None |
| SRAE_X000147300 | Astacin | 3 | 3 / 2 / 0 | 9.7 | None | None |
| SRAE_0000081500 | CAP domain-containing protein | 5.6 | 2 / 3 / 0 | 9.7 | None | None |

**Supplementary Table 7.** The hundred most free-living genes.. Gene gives the gene’s designation; Predicted function is the WormBase ParaSite description of each gene; Coding SNPs per kb is the number of SNPs per kb within the coding sequence of that gene; SNP type is the absolute number of synonymous (S), nonsynonymous (NS) and STOP codon-causing SNPs; Fold change is the log_2_ difference in expression of the gene between the parasitic female and free-living female morphs taken from Hunt *et al.,* 2016, here shown as positive values indicating greater expression in the free-living female morph; Expansion cluster is expansion cluster or associated flanking region gene belongs to, if any; Variable region is as defined in **Supplementary** Table 5.

| Gene | Predicted function | Coding SNPs per kb | SNP type (S/NS/STOP) | Fold Change | Expansion cluster | Variable region |
| --- | --- | --- | --- | --- | --- | --- |
| SRAE_2000529300 | ShKT domain-containing protein | 13.5 | 0 / 4 / 0 | 11 | None | None |
| SRAE_2000226500 | ShKT domain-containing protein | 41.5 | 5 / 17 / 0 | 10.8 | None | None |
| SRAE_X000147400 | Phosphate-regulating neutral endopeptidase | 4.8 | 2 / 5 / 0 | 10.4 | None | None |
| SRAE_1000161500 | DUF148 domain-containing | 1.6 | 0 / 1 / 0 | 10.3 | None | None |
| SRAE_X000018600 | Mucin 18B | 2.4 | 2 / 2 / 0 | 10.3 | None | None |
| SRAE_X000158400 | Astacin-like metalloendopeptidase | 1.4 | 0 / 2 / 0 | 10.1 | None | 46 |
| SRAE_2000474600 | ShKT domain-containing protein | 2.2 | 0 / 2 / 0 | 10.1 | None | None |
| SRAE_X000135300 | Collagen alpha-5(IV) chain | 1 | 1 / 0 / 0 | 10 | None | None |
| SRAE_2000463500 | Hypothetical protein | 5.5 | 1 / 1 / 1 | 9.8 | None | None |
| SRAE_2000079600 | Transcription factor Sp6 | 5.3 | 3 / 1 / 0 | 8.8 | None | None |
| SRAE_1000062000 | Nematode cuticle collagen | 5.3 | 4 / 1 / 0 | 8.8 | None | None |
| SRAE_2000491300 | ShKT domain-containing protein | 0 | 0 / 0 / 0 | 8.7 | None | None |
| SRAE_1000170700 | Hypothetical protein | 6.9 | 8 / 2 / 0 | 8.6 | None | None |
| SRAE_1000213600 | Aspartic peptidase family | 2.2 | 2 / 1 / 0 | 8.5 | None | None |
| SRAE_2000400600 | Saposin-like type B, 1 domain | 0 | 0 / 0 / 0 | 8.4 | None | None |
| SRAE_2000477400 | ShKT domain-containing protein | 8.4 | 1 / 5 / 0 | 8.4 | None | None |
| SRAE_1000227500 | Hypothetical protein | 3.5 | 0 / 2 / 0 | 8.2 | None | None |
| SRAE_2000292900 | GH07323p | 1.2 | 1 / 1 / 0 | 8 | None | None |
| SRAE_X000034300 | Hypothetical protein | 8 | 1 / 5 / 0 | 8 | None | None |
| SRAE_2000126900 | Nematode cuticle collagen | 2.1 | 1 / 1 / 0 | 7.9 | FR4R | None |
| SRAE_X000032100 | Lipase, class 3 family-containing | 3.1 | 2 / 1 / 0 | 7.9 | None | None |
| SRAE_1000151800 | Protein COL-120 | 4.1 | 4 / 1 / 0 | 7.7 | None | None |
| SRAE_X000215700 | Hypothetical protein | 9.6 | 2 / 1 / 0 | 7.7 | None | None |
| SRAE_2000363400 | Hypothetical protein | 2.8 | 1 / 0 / 0 | 7.5 | None | None |
| SRAE_2000434700 | Collagen alpha-5(IV) chain | 1 | 0 / 1 / 0 | 7.5 | None | None |
| SRAE_1000352300 | Serine/threonine- /dual specificity protein kinase | 0.8 | 1 / 0 / 0 | 7.4 | None | None |
| SRAE_1000099100 | Hypothetical protein | 0.7 | 1 / 0 / 0 | 7.4 | None | None |
| SRAE_X000100400 | Hypothetical protein | 0.9 | 0 / 1 / 0 | 7.4 | None | None |
| SRAE_1000220400 | Hypothetical protein | 0 | 0 / 0 / 0 | 7.4 | None | None |
| SRAE_2000033400 | Heat shock protein Hsp-12.2 | 0 | 0 / 0 / 0 | 7.2 | None | None |
| SRAE_1000228700 | Cell death specification protein 2 | 1.2 | 1 / 0 / 0 | 7.2 | None | None |
| SRAE_X000095500 | Hypothetical protein | 0 | 0 / 0 / 0 | 7.2 | None | None |
| SRAE_1000073100 | Protein lin-32 | 0 | 0 / 0 / 0 | 7.2 | None | None |
| SRAE_1000271200 | MSP domain; PapD-like domain-containing | 0 | 0 / 0 / 0 | 7.2 | None | None |
| SRAE_2000115300 | Metallothionein, family 4, echinoidea-containing | 3 | 0 / 2 / 0 | 7.1 | None | None |
| SRAE_1000098100 | Hypothetical protein | 0 | 0 / 0 / 0 | 7.1 | None | None |
| SRAE_2000006500 | Hypothetical protein | 0 | 0 / 0 / 0 | 7 | None | None |
| SRAE_2000425600 | Hypothetical protein | 0 | 0 / 0 / 0 | 7 | None | None |
| SRAE_0000045200 | von Willebrand factor, type A domain-containing | 3.2 | 10 / 8 / 0 | 7 | None | None |
| SRAE_1000198200 | Glycoside hydrolase | 1.2 | 1 / 1 / 0 | 7 | None | None |
| SRAE_1000159100 | Hypothetical protein | 0.7 | 0 / 1 / 0 | 6.9 | None | None |
| SRAE_1000268100 | Nematode cuticle collagen | 0 | 0 / 0 / 0 | 6.9 | None | None |
| SRAE_2000473500 | Sulfotransferase family | 11 | 5 / 3 / 3 | 6.9 | None | None |
| SRAE_X000144300 | Protein GLF-1 | 0.6 | 0 / 1 / 0 | 6.8 | FR14R | None |
| SRAE_X000086900 | Hypothetical protein | 7.2 | 1 / 4 / 0 | 6.8 | None | None |
| SRAE_1000058300 | Hypothetical protein | 7.6 | 3 / 6 / 0 | 6.8 | None | None |
| SRAE_X000224800 | Nematode cuticle collagen | 1.9 | 1 / 1 / 0 | 6.8 | None | None |
| SRAE_1000033500 | Collagen alpha-5(IV) chain | 0.9 | 0 / 1 / 0 | 6.7 | None | None |
| SRAE_1000004900 | Glycoside hydrolase, family 25 | 1.3 | 1 / 0 / 0 | 6.7 | None | None |
| SRAE_1000015400 | Lipase EstA/Esterase EstB family-containing | 2 | 1 / 1 / 0 | 6.7 | None | None |
| SRAE_2000439500 | Collagen alpha-5(IV) chain | 0 | 0 / 0 / 0 | 6.7 | None | None |
| SRAE_X000039100 | Brain-specific homeobox protein homolog | 0 | 0 / 0 / 0 | 6.7 | None | None |
| SRAE_1000049700 | Hypothetical protein | 0 | 0 / 0 / 0 | 6.7 | None | None |
| SRAE_1000036400 | Histidine decarboxylase | 12 | 3 / 4 / 0 | 6.6 | None | None |
| SRAE_2000476800 | Hypothetical protein | 0 | 0 / 0 / 0 | 6.5 | None | None |
| SRAE_2000431300 | Nematode fatty acid retinoid binding family-containing | 0 | 0 / 0 / 0 | 6.5 | None | None |
| SRAE_X000210100 | Domain of unknown function DB domain-containing | 1.3 | 1 / 0 / 0 | 6.4 | None | None |
| SRAE_2000437000 | CAP domain-containing protein | 12.6 | 3 / 5 / 0 | 6.4 | None | None |
| SRAE_X000195000 | Si:ch211-105d4.5 | 0 | 0 / 0 / 0 | 6.4 | None | None |
| SRAE_2000145200 | Nematode cuticle collagen | 0.9 | 1 / 0 / 0 | 6.4 | None | None |
| SRAE_2000482000 | N-acylethanolamine-hydrolyzing acid amidase | 1.7 | 1 / 1 / 0 | 6.4 | None | None |
| SRAE_1000115300 | Hypothetical protein | 1.3 | 1 / 1 / 0 | 6.3 | None | None |
| SRAE_2000459400 | Acyl-CoA N-acyltransferase domain-containing | 0 | 0 / 0 / 0 | 6.3 | None | None |
| SRAE_2000468700 | Protein dyf-8 | 0 | 0 / 0 / 0 | 6.3 | None | None |
| SRAE_2000214100 | Nematode cuticle collagen | 1 | 1 / 0 / 0 | 6.3 | None | None |
| SRAE_2000481600 | MD-2-related lipid-recognition domain | 1.9 | 1 / 0 / 0 | 6.3 | None | None |
| SRAE_1000158100 | Hypothetical protein | 2.5 | 2 / 2 / 0 | 6.2 | None | None |
| SRAE_0000015000 | MD-2-related lipid-recognition domain-containing | 3.7 | 0 / 2 / 0 | 6.2 | None | None |
| SRAE_2000365700 | Glycoside hydrolase, catalytic domain | 2.5 | 0 / 1 / 1 | 6.2 | None | None |
| SRAE_2000466700 | Properdin | 5 | 7 / 4 / 0 | 6 | None | None |
| SRAE_2000324800 | Hypothetical protein | 16.2 | 15 / 5 / 0 | 6 | None | None |
| SRAE_2000014200 | Saposin B domain | 2.4 | 0 / 1 / 0 | 5.9 | None | None |
| SRAE_1000271800 | MSP domain; PapD-like domain-containing | 0 | 0 / 0 / 0 | 5.9 | None | None |
| SRAE_1000068700 | Thrombospondin, type 1 repeat-containing | 6.7 | 4 / 0 / 0 | 5.9 | None | None |
| SRAE_X000188000 | Sphingosine-1-phosphate lyase 1 | 2.2 | 2 / 4 / 0 | 5.8 | None | None |
| SRAE_X000258800 | Thioredoxin-like fold domain-containing | 0 | 0 / 0 / 0 | 5.7 | None | None |
| SRAE_2000009100 | Protein-tyrosine phosphatase | 1.1 | 0 / 1 / 0 | 5.7 | None | None |
| SRAE_0000012300 | Hypothetical protein | 4.7 | 0 / 2 / 0 | 5.7 | None | None |
| SRAE_2000448200 | Hypothetical protein | 5.6 | 0 / 2 / 0 | 5.6 | None | None |
| SRAE_X000101600 | Hypothetical protein | 5.6 | 5 / 3 / 0 | 5.6 | None | None |
| SRAE_2000041300 | LDLR class B repeat | 0 | 0 / 0 / 0 | 5.5 | None | None |
| SRAE_X000083300 | Protein CUTL-16 | 2.1 | 2 / 1 / 0 | 5.5 | None | None |
| SRAE_X000079800 | Hypothetical protein | 0 | 0 / 0 / 0 | 5.5 | None | None |
| SRAE_X000138300 | Protein mesh | 0 | 0 / 0 / 0 | 5.4 | None | None |
| SRAE_1000020600 | Hypothetical protein | 4.3 | 0 / 2 / 0 | 5.4 | None | None |
| SRAE_2000476700 | Hypothetical protein | 0 | 0 / 0 / 0 | 5.4 | None | None |
| SRAE_2000347100 | Lipase EstA/Esterase EstB family-containing | 1 | 0 / 1 / 0 | 5.4 | None | None |
| SRAE_2000380600 | GPCR, rhodopsin-like | 0.9 | 0 / 1 / 0 | 5.4 | None | None |
| SRAE_2000052400 | Alpha crystallin/Hsp20 domain | 1.2 | 3 / 1 / 0 | 5.4 | None | None |
| SRAE_X000138500 | Lipase, class 3 family-containing | 8.8 | 4 / 4 / 0 | 5.3 | None | None |
| SRAE_X000166100 | Protein CDH-10 | 3.2 | 5 / 12 / 0 | 5.3 | None | None |
| SRAE_1000022400 | Epidermal growth factor-like domain | 2.7 | 11 / 7 / 0 | 5.3 | None | None |
| SRAE_X000157300 | Homeobox protein HMX1 | 2.5 | 1 / 2 / 0 | 5.3 | None | None |
| SRAE_X000043300 | Proteasomal ubiquitin receptor ADRM1 homolog | 6.3 | 0 / 7 / 0 | 5.3 | None | None |
| SRAE_2000424300 | Fatty-acid amide hydrolase 2 | 1.8 | 3 / 0 / 0 | 5.3 | None | None |
| SRAE_X000098100 | Astacin-like metalloendopeptidase | 1.2 | 0 / 2 / 0 | 5.3 | None | None |
| SRAE_X000095600 | Hypothetical protein | 4.1 | 0 / 2 / 0 | 5.2 | None | None |
| SRAE_2000482100 | Hypothetical protein | 11.4 | 6 / 11 / 0 | 5.2 | None | None |
| SRAE_2000377500 | Nematode cuticle collagen | 2 | 2 / 0 / 0 | 5.2 | None | None |
| SRAE_2000360100 | Alpha amylase family; | 5 | 3 / 3 / 0 | 5.2 | None | None |

**Supplementary Table 8.** *S. ratti* expansion clusters, revised after further inspection of genome assembly in the cluster region.. Region shows the numbered Flanking Regions (FR) and Expansion Clusters (EC); Gene the genes present within these regions; Predicted function is the WormBase ParaSite’s description of the gene; Coding SNPs per kb is the number of SNPs per kb within the coding sequence of that gene; SNP type is synonymous (S), non-synonymous (NS) or STOP-causing (STOP); Expression shows whether the gene is upregulated (with a difference in expression of log_2_-fold of at least 1 being considered upregulation) in the parasitic adult female morph (Parasitic), the free-living adult female morph (Free-living), or not differentially expressed (Same), all taken from Hunt *et al.,* 2016, or unlisted there.

| Region | Gene | Predicted function | Coding SNPs per kb | SNP type (S/NS/STOP) | Expression |
| --- | --- | --- | --- | --- | --- |
| FR1L | SRAE_1000180400 | Transcriptional regulator ATRX | 3.7 | 6 / 5 / 0 | Parasitic |
|  | SRAE_1000180500 | Formin | 2.8 | 4 / 5 / 0 | Same |
|  | SRAE_1000180600 | Hypothetical protein | 1.4 | 0 / 2 / 0 | Same |
|  | SRAE_1000180700 | Glycosylphosphatidylinositol-mannosyltransferase I | 1.7 | 0 / 1 / 0 | Same |
|  | SRAE_1000180800 | Hypothetical protein | 1.9 | 1 / 0 / 0 | Same |
|  | SRAE_1000180900 | Rhodopsin-like | 0.7 | 1 / 0 / 0 | Same |
|  | SRAE_1000181000 | TPM domain-containing protein | 1.4 | 0 / 1 / 0 | Same |
|  | SRAE_1000181100 | MIF4-like | 1.2 | 1 / 1 / 0 | Same |
| | SRAE_1000181200 | Sodium/potassium-transporting ATPase $\alpha$ subunit | 0 | 0 / 0 / 0 | Free-living |
|  | SRAE_1000181300 | Hypothetical protein | 0 | 0 / 0 / 0 | Free-living |
|  | SRAE_1000181400 | Mediator of RNA polymerase II transcription subunit 9 | 5.2 | 2 / 0 / 0 | Same |
|  | SRAE_1000181500 | Serine/threonine-protein kinase Chk1 | 0.7 | 1 / 0 / 0 | Same |
|  | SRAE_1000181600 | Hypothetical protein | 1.1 | 1 / 0 / 0 | Same |
|  | SRAE_1000181700 | Hypothetical protein | 2.7 | 3 / 1 / 0 | Same |
|  | SRAE_1000181800 | Hypothetical protein | 1.2 | 0 / 1 / 0 | Free-living |
|  | SRAE_1000181900 | Eukaryotic translation initiation factor 2A | 2.5 | 0 / 4 / 0 | Same |
|  | SRAE_1000182000 | Hypothetical protein | 0 | 0 / 0 / 0 | Same |
|  | SRAE_1000182100 | Heme transporter HRG | 0 | 0 / 0 / 0 | Same |
| EC1 | SRAE_1000182200 | CAP domain-containing | 1.1 | 0 / 1 / 0 | Parasitic |
|  | SRAE_1000182300 | CAP domain-containing | 0 | 0 / 0 / 0 | Parasitic |
|  | SRAE_1000182400 | CAP domain-containing | 1.2 | 1 / 0 / 0 | Parasitic |
|  | SRAE_1000183300 | CAP domain-containing | 3.4 | 2 / 1 / 0 | Parasitic |
| FR1R | SRAE_1000183400 | Nanchung | 1.4 | 1 / 2 / 0 | Same |
|  | SRAE_1000183500 | Hypothetical protein | 0 | 0 / 0 / 0 | Free-living |
|  | SRAE_1000183600 | SMc04008-like domain-containing | 1.4 | 0 / 3 / 0 | Same |
|  | SRAE_1000183700 | Hypothetical protein | 0 | 0 / 0 / 0 | Unlisted |
|  | SRAE_1000183800 | 1,2-dihydroxy-3-keto-5-methylthiopentene dioxxygenase | 0 | 0 / 0 / 0 | Same |
|  | SRAE_1000183900 | Transmembrane receptor | 7.1 | 9 / 4 / 0 | Parasitic |
|  | SRAE_1000184000 | Predicted transmembrane / coiled-coil 2-containing | 0.7 | 0 / 1 / 0 | Unlisted |
|  | SRAE_1000184100 | Band 7 protein family and Stomatin family-containing | 3.2 | 3 / 0 / 0 | Unlisted |
|  | SRAE_1000184200 | Basic-leucine zipper domain-containing | 2.5 | 3 / 2 / 0 | Unlisted |
|  | SRAE_1000184300 | TBC1 domain family member 13 | 1.6 | 2 / 0 / 0 | Unlisted |
|  | SRAE_1000184400 | BTB/POZ domain-containing | 0.7 | 1 / 0 / 0 | Unlisted |
| FR2L | SRAE_2000075900 | NHR/GATA-type domain-containing | 2.5 | 4 / 2 / 0 | Same |
|  | SRAE_2000076000 | MIP20649p | 3.2 | 6 / 3 / 0 | Same |
|  | SRAE_2000076100 | MIP20649p | 0.8 | 1 / 0 / 0 | Unlisted |
|  | SRAE_2000076200 | Hypothetical protein | 5.7 | 10 / 5 / 0 | Free-living |
|  | SRAE_2000076300 | Hypothetical protein | 2.4 | 2 / 3 / 0 | Parasitic |
| EC2 | SRAE_2000076400 | CAP domain-containing | 29.8 | 7 / 13 / 0 | Parasitic |
|  | SRAE_2000076600 | CAP domain-containing | 21.2 | 3 / 14 / 0 | Parasitic |
|  | SRAE_2000076700 | CAP domain-containing | 60.4 | 11 / 39 / 0 | Parasitic |
|  | SRAE_2000076800 | CAP domain-containing | 39.2 | 11 / 18 / 1 | Parasitic |
|  | SRAE_2000076900 | CAP domain-containing | 36.8 | 6 / 25 / 0 | Parasitic |
|  | SRAE_2000077000 | CAP domain-containing | 19 | 6 / 9 / 1 | Parasitic |
|  | SRAE_2000077100 | CAP domain-containing | 14 | 6 / 6 / 0 | Parasitic |
|  | SRAE_2000077200 | UDP-glucosyltransferase | 12.7 | 12 / 8 / 0 | Parasitic |
|  | SRAE_2000077300 | CAP domain-containing | 17.4 | 5 / 10 / 0 | Parasitic |
|  | SRAE_2000077400 | CAP domain-containing | 13 | 4 / 7 / 0 | Parasitic |
|  | SRAE_2000077500 | CAP domain-containing | 4.4 | 2 / 4 / 0 | Same |
| FR2R | SRAE_2000077600 | Hypothetical protein | 10 | 8 / 14 / 0 | Same |
|  | SRAE_2000077700 | BTB/POZ domain-containing | 7.9 | 1 / 4 / 1 | Unlisted |
|  | SRAE_2000077800 | Hypothetical protein | 2.9 | 1 / 0 / 0 | Unlisted |
|  | SRAE_2000077900 | Protein kinase-like domain-containing | 2.1 | 1 / 2 / 0 | Unlisted |
|  | SRAE_2000078000 | Ubiquitin-conjugating enzyme, E2 domain | 4.5 | 2 / 0 / 0 | Same |
|  | SRAE_2000078100 | WH2 domain-containing | 5.9 | 3 / 0 / 0 | Same |
|  | SRAE_2000078200 | Casein kinase II subunit beta | 8.3 | 2 / 3 / 0 | Same |
|  | SRAE_2000078300 | Hypothetical protein | 0 | 0 / 0 / 0 | Unlisted |
|  | SRAE_2000078400 | Hypothetical protein | 0 | 0 / 0 / 0 | Same |
|  | SRAE_2000078500 | Serine/threonine-protein kinase haspin | 11.1 | 7 / 9 / 0 | Same |
|  | SRAE_2000078600 | PDZ domain-containing | 1.4 | 0 / 1 / 0 | Unlisted |
|  | SRAE_2000078700 | CAP domain-containing | 15.2 | 5 / 10 / 0 | Parasitic |
|  | SRAE_2000078800 | Hypothetical protein | 1.4 | 4 / 0 / 0 | Same |
| FR3L | SRAE_2000123100 | Transcription elongation factor SPT5 | 0.4 | 0 / 1 / 0 | Same |
|  | SRAE_2000123200 | Armadillo-like helical domain -containing | 1.3 | 3 / 3 / 0 | Same |
|  | SRAE_2000123300 | Hypothetical protein | 0 | 0 / 0 / 0 | Unlisted |
|  | SRAE_2000123400 | Gamma-aminobutyric acid receptor subunit beta | 2 | 3 / 0 / 0 | Same |
|  | SRAE_2000123500 | Hypothetical protein | 6.1 | 2 / 1 / 0 | Unlisted |
|  | SRAE_2000123600 | Phosphodiesterase | 0.9 | 2 / 0 / 0 | Same |
|  | SRAE_2000123700 | Glycosyl transferase | 3.4 | 1 / 4 / 0 | Same |
|  | SRAE_2000123800 | Tyrosine-protein kinase | 0 | 0 / 0 / 0 | Unlisted |
|  | SRAE_2000123900 | Bax inhibitor 1-related family-containing | 0 | 0 / 0 / 0 | Unlisted |
|  | SRAE_2000124000 | Bloom syndrome protein | 2 | 3 / 6 / 0 | Free-living |
|  | SRAE_2000124100 | Protein lethal(2)essential for life | 7.8 | 4 / 1 / 0 | Parasitic |
|  | SRAE_2000124200 | Hypothetical protein | 14.5 | 5 / 21 / 0 | Parasitic |
| EC3 | SRAE_2000124300 | CAP domain-containing | 69 | 17 / 49 / 4 | Parasitic |
|  | SRAE_2000124400 | CAP domain-containing | 43.3 | 8 / 30 / 1 | Parasitic |
|  | SRAE_2000124500 | CAP domain-containing | 50.9 | 11 / 33 / 2 | Parasitic |
|  | SRAE_2000124600 | CAP domain-containing | 8.7 | 3 / 5 / 0 | Parasitic |
|  | SRAE_2000124700 | CAP domain-containing | 5.2 | 0 / 4 / 1 | Parasitic |
|  | SRAE_2000124800 | CAP domain-containing | 6.5 | 4 / 2 / 0 | Parasitic |
|  | SRAE_2000124900 | CAP domain-containing | 6.6 | 2 / 4 / 0 | Parasitic |
| FR3R | SRAE_2000126100 | Hypothetical protein | 8.6 | 6 / 2 / 0 | Parasitic |
|  | SRAE_2000126200 | Hypothetical protein | 50 | 3 / 17 / 1 | Unlisted |
|  | SRAE_2000126290 | Hypothetical protein | 10.7 | 3 / 7 / 0 | Unlisted |
|  | SRAE_2000126300 | Hypothetical protein | 14.1 | 6 / 7 / 0 | Parasitic |
|  | SRAE_2000126400 | Hypothetical protein | 11.8 | 3 / 7 / 1 | Parasitic |
|  | SRAE_2000126500 | Hypothetical protein | 0 | 0 / 0 / 0 | Parasitic |
|  | SRAE_2000126600 | CAP domain-containing | 18.5 | 6 / 11 / 0 | Parasitic |
|  | SRAE_2000126700 | Hypothetical protein | 3.6 | 1 / 2 / 0 | Same |
|  | SRAE_2000126800 | Serine/threonine-protein phosphatase | 0.5 | 0 / 1 / 0 | Unlisted |
|  | SRAE_2000126900 | Nematode cuticle collagen | 2.1 | 1 / 1 / 0 | Free-living |
|  | SRAE_2000127000 | Hypothetical protein | 8.5 | 3 / 7 / 0 | Unlisted |
|  | SRAE_2000127100 | Hypothetical protein | 2.5 | 1 / 1 / 0 | Unlisted |
|  | SRAE_2000127200 | Flavin-containing monooxygenase | 2.9 | 2 / 2 / 0 | Same |
|  | SRAE_2000127300 | Serine/threonine-protein kinase RIO1 | 2.6 | 3 / 1 / 0 | Same |
|  | SRAE_2000127400 | Hypothetical protein | 1.9 | 2 / 2 / 0 | Same |
| FR5L | SRAE_2000325100 | IA-2 ortholog | 2 | 2 / 5 / 0 | Same |
|  | SRAE_2000325200 | Hypothetical protein | 4.7 | 0 / 1 / 0 | Unlisted |
|  | SRAE_2000325300 | Integrator complex subunit 9 | 0.5 | 0 / 1 / 0 | Same |
|  | SRAE_2000325400 | Hypothetical protein | 0 | 0 / 0 / 0 | Unlisted |
|  | SRAE_2000325500 | UDP-glucuronosyltransferase | 3.8 | 2 / 4 / 0 | Free-living |
| EC5 | SRAE_2000325600 | Astacin-like metalloendopeptidase | 0.9 | 1 / 0 / 0 | Parasitic |
|  | SRAE_2000326000 | Astacin-like metalloendopeptidase | 7.6 | 3 / 7 / 0 | Parasitic |
| FR5R | SRAE_2000326100 | Epidermal growth factor-like domain-containing | 0.6 | 1 / 1 / 0 | Free-living |
| FR6L | SRAE_2000450300 | UDP-glucosyltransferase | 5.7 | 2 / 7 / 0 | Parasitic |
| EC6 | SRAE_2000450400 | Astacin-like metalloendopeptidase | 1.9 | 1 / 2 / 0 | Parasitic |
|  | SRAE_2000450500 | Astacin-like metalloendopeptidase | 4 | 1 / 4 / 0 | Parasitic |
|  | SRAE_2000450600 | Astacin-like metalloendopeptidase | 6.5 | 2 / 6 / 0 | Parasitic |
|  | SRAE_2000450700 | Astacin-like metalloendopeptidase | 22.3 | 7 / 25 / 0 | Parasitic |
| FR6R | SRAE_2000450800 | Nematode fatty acid retinoid binding | 5.5 | 3 / 0 / 0 | Unlisted |
|  | SRAE_2000450900 | Hypothetical protein | 5.3 | 1 / 5 / 0 | Unlisted |
|  | SRAE_2000451000 | Hypothetical protein | 22.2 | 6 / 3 / 0 | Parasitic |
| FR7L | SRAE_2000452400 | Hypothetical protein | 3.6 | 3 / 0 / 0 | Unlisted |
|  | SRAE_2000452500 | ATP-binding cassette sub-family D member 4 | 2.3 | 3 / 1 / 0 | Free-living |
|  | SRAE_2000452600 | ATP-binding cassette sub-family D member 4 | 1.1 | 1 / 1 / 0 | Parasitic |
|  | SRAE_2000452700 | Hypothetical protein | 1.7 | 1 / 2 / 0 | Unlisted |
|  | SRAE_2000452800 | Hypothetical protein | 1.8 | 3 / 0 / 0 | Unlisted |
|  | SRAE_2000452900 | Hypothetical protein | 0 | 0 / 0 / 0 | Unlisted |
|  | SRAE_2000453000 | Hypothetical protein | 0 | 0 / 0 / 0 | Same |
|  | SRAE_2000453100 | Hypothetical protein | 20.9 | 7 / 10 / 0 | Parasitic |
| EC7 | SRAE_2000453200 | Astacin-like metalloendopeptidase | 9.6 | 3 / 8 / 0 | Parasitic |
|  | SRAE_2000453300 | Astacin-like metalloendopeptidase | 2.6 | 0 / 3 / 0 | Parasitic |
|  | SRAE_2000453400 | Sulfotransferase family-containing | 0 | 0 / 0 / 0 | Unlisted |
|  | SRAE_2000453500 | Astacin-like metalloendopeptidase | 2.5 | 0 / 3 / 0 | Parasitic |
|  | SRAE_2000453600 | Astacin-like metalloendopeptidase | 0 | 0 / 0 / 0 | Parasitic |
|  | SRAE_2000453700 | Astacin-like metalloendopeptidase | 5.5 | 0 / 7 / 0 | Parasitic |
|  | SRAE_2000453800 | Astacin-like metalloendopeptidase | 34.8 | 15 / 26 / 0 | Parasitic |
|  | SRAE_2000453900 | Astacin-like metalloendopeptidase | 5.1 | 4 / 2 / 0 | Parasitic |
| FR7R | SRAE_2000454000 | Zinc metalloproteinase | 3.4 | 3 / 1 / 0 | Unlisted |
|  | SRAE_2000454100 | Hypothetical protein | 3.7 | 7 / 4 / 0 | Free-living |
|  | SRAE_2000454200 | Hypothetical protein | 0 | 0 / 0 / 0 | Same |
|  | SRAE_2000454300 | Sulfotransferase | 4.2 | 4 / 0 / 0 | Same |
|  | SRAE_2000454400 | Sulfotransferase | 1.8 | 2 / 0 / 0 | Same |
|  | SRAE_2000454500 | Estradiol 17-beta-dehydrogenase 12 | 0 | 0 / 0 / 0 | Parasitic |
|  | SRAE_2000454600 | Alpha-(1,3)-fucosyltransferase C | 3.1 | 0 / 3 / 0 | Unlisted |
|  | SRAE_2000454700 | Glycosyl transferase, family 14-containing | 2.3 | 1 / 2 / 0 | Same |
| FR8L | SRAE_2000454800 | Cytochrome P450 4V2 | 1.3 | 1 / 1 / 0 | Same |
| EC8 | SRAE_2000454900 | Acetylcholinesterase | 1.2 | 2 / 0 / 0 | Parasitic |
|  | SRAE_2000455000 | Astacin-like metalloendopeptidase | 2.6 | 1 / 2 / 0 | Parasitic |
|  | SRAE_2000455100 | Hypothetical protein | 1.6 | 1 / 0 / 0 | Parasitic |
|  | SRAE_2000455200 | CAP domain-containing | 0 | 0 / 0 / 0 | Parasitic |
|  | SRAE_2000455300 | Acetylcholinesterase | 0 | 0 / 0 / 0 | Parasitic |
| FR8R | SRAE_2000455400 | Hypothetical protein | 0 | 0 / 0 / 0 | Unlisted |
|  | SRAE_2000455500 | Hypothetical protein | 0 | 0 / 0 / 0 | Same |
|  | SRAE_2000455600 | Zona pellucida domain-containing | 0.9 | 0 / 1 / 0 | Unlisted |
|  | SRAE_2000455700 | Hypothetical protein | 0 | 0 / 0 / 0 | Free-living |
|  | SRAE_2000455800 | Hypothetical protein | 0 | 0 / 0 / 0 | Unlisted |
| FR9L | SRAE_2000497000 | Hypothetical protein | 1.4 | 5 / 4 / 0 | Free-living |
| EC9 | SRAE_2000497100 | Astacin-like metalloendopeptidase | 5.2 | 1 / 5 / 0 | Parasitic |
| FR9R | SRAE_2000497600 | Cad96Cb | 4.5 | 9 / 0 / 0 | Parasitic |
|  | SRAE_2000497700 | Zinc finger, RING/FYVE/PHD-type domain-containing | 1.5 | 1 / 0 / 0 | Parasitic |
|  | SRAE_2000497800 | Protein lethal(2)essential for life | 0 | 0 / 0 / 0 | Parasitic |
| FR10L | SRAE_2000522800 | Trypsin Inhibitor-like, cysteine rich domain-containing | 22.7 | 10 / 5 / 0 | Parasitic |
|  | SRAE_2000522900 | Glycosyl transferase, family 14-containing | 4.3 | 2 / 4 / 0 | Unlisted |
|  | SRAE_2000523000 | Transthyretin-like family-containing | 9.1 | 3 / 1 / 0 | Parasitic |
|  | SRAE_2000523100 | 7TM GPCR, (Sre) family-containing | 0 | 0 / 0 / 0 | Unlisted |
|  | SRAE_2000523200 | 7TM GPCR, (Sre) family-containing | 4.3 | 0 / 2 / 0 | Unlisted |
|  | SRAE_2000523300 | Hypothetical protein | 1.4 | 0 / 1 / 0 | Unlisted |
|  | SRAE_2000523400 | Nuclear hormone receptor | 3.9 | 4 / 1 / 0 | Same |
|  | SRAE_2000523500 | Proteinase inhibitor I25 | 0 | 0 / 0 / 0 | Free-living |
|  | SRAE_2000523600 | Carboxylic ester hydrolase | 3.4 | 2 / 4 / 0 | Unlisted |
| EC10 | SRAE_2000523700 | Astacin-like metalloendopeptidase | 5.2 | 1 / 5 / 0 | Parasitic |
|  | SRAE_2000523800 | Astacin-like metalloendopeptidase | 2.8 | 2 / 2 / 0 | Parasitic |
|  | SRAE_2000523900 | Astacin-like metalloendopeptidase | 26.1 | 8 / 29 / 1 | Parasitic |
|  | SRAE_2000524000 | Astacin-like metalloendopeptidase | 4 | 1 / 4 / 0 | Parasitic |
| FR12L | SRAE_2000526900 | Importin-beta | 2.2 | 4 / 2 / 0 | Same |
|  | SRAE_2000527000 | Hypothetical protein | 14.3 | 8 / 17 / 0 | Unlisted |
| EC12 | SRAE_2000527100 | CAP domain-containing | 71.1 | 9 / 54 / 1 | Parasitic |
|  | SRAE_2000527200 | CAP domain-containing | 18.6 | 1 / 15 / 1 | Parasitic |
|  | SRAE_2000527300 | CAP domain-containing | 4.6 | 2 / 2 / 0 | Parasitic |
|  | SRAE_2000527400 | CAP domain-containing | 5.5 | 2 / 3 / 0 | Unlisted |
|  | SRAE_2000527500 | CAP domain-containing | 3.3 | 3 / 0 / 0 | Parasitic |
|  | SRAE_2000527600 | CAP domain-containing | 2.2 | 0 / 2 / 0 | Parasitic |
|  | SRAE_2000527700 | CAP domain-containing | 1.1 | 0 / 1 / 0 | Parasitic |
| FR12R | SRAE_2000527800 | Hypothetical protein | 18.5 | 2 / 10 / 0 | Unlisted |
|  | SRAE_2000527900 | Hypothetical protein | 41.5 | 9 / 17 / 0 | Unlisted |
|  | SRAE_2000528000 | Hypothetical protein | 1.8 | 0 / 1 / 0 | Unlisted |
|  | SRAE_2000528100 | Hypothetical protein | 3 | 3 / 4 / 0 | Unlisted |
| FR14L | SRAE_X000143400 | Ferric-chelate reductase 1 | 10.1 | 5 / 3 / 0 | Unlisted |
|  | SRAE_X000143500 | Hypothetical protein | 7.1 | 7 / 9 / 0 | Same |
|  | SRAE_X000143600 | 1x GPCR, rhodopsin-like, 7TM domain-containing | 0.7 | 1 / 0 / 0 | Same |
|  | SRAE_X000143700 | Hypothetical protein | 4.1 | 3 / 1 / 0 | Same |
| EC14 | SRAE_X000143800 | Astacin-like metalloendopeptidase | 15.5 | 8 / 9 / 1 | Parasitic |
|  | SRAE_X000143900 | Astacin-like metalloendopeptidase | 18.3 | 8 / 13 / 2 | Parasitic |
|  | SRAE_X000144000 | Astacin-like metalloendopeptidase | 7.2 | 4 / 5 / 0 | Parasitic |
|  | SRAE_X000144100 | Astacin-like metalloendopeptidase | 2.4 | 0 / 3 / 0 | Parasitic |
|  | SRAE_X000144200 | Astacin-like metalloendopeptidase | 39.5 | 24 / 24 / 1 | Parasitic |
|  | SRAE_X000144210 | Astacin-like metalloendopeptidase | 21.5 | 6 / 21 / 0 | Parasitic |
| FR14R | SRAE_X000144220 | Hypothetical protein | 9.4 | 4 / 4 / 0 | Same |
|  | SRAE_X000144300 | Amine oxidase domain-containing | 0.6 | 0 / 1 / 0 | Free-living |
| EC15 | SRAE_X000146800 | Astacin-like metalloendopeptidase | 29.3 | 16 / 21 / 0 | Parasitic |
|  | SRAE_X000146900 | Astacin-like metalloendopeptidase | 47.5 | 25 / 34 / 0 | Free-living |
| FR15R | SRAE_X000146950 | Hypothetical protein | 3.2 | 1 / 0 / 0 | Unlisted |
|  | SRAE_X000147000 | Hypothetical protein | 5.1 | 0 / 1 / 0 | Parasitic |
|  | SRAE_X000147100 | Hypothetical protein | 6.3 | 2 / 1 / 0 | Unlisted |
|  | SRAE_X000147200 | Hypothetical protein | 26.9 | 15 / 22 / 0 | Parasitic |

**Supplementary Table 9.** dN/dS ratios of expansion clusters and their flanking regions.. The dN/dS ratio for genes in expansion clusters and in parentheses the number of genes, N. We calculated dN/dS ratios by analysing parasites from clades 1 and 3 (**Figure 3**), treating each as separate populations.

| Expansion cluster | Expansion cluster mean dN/dS ratio (N) | Flanking region mean dN/dS ratio (N) |
| --- | --- | --- |
| 2 | 0.257 (1) | 2.088 (2) |
| 3 | 0.454 (5) | 0.735 (6) |
| 6 | 1.373 (1) | 0.408 (1) |
| 7 | 3.419 (2) | 0.278 (3) |
| 14 | 0.322 (3) | 1.242 (1) |

**Supplementary Table 10.** Sampling sites and times.

| Site (code) | Coordinates | Type |
| --- | --- | --- |
| Cardiff (CA) | 51°29'54"N 3°07'25"W | Industrial |
| Avonmouth (AM) | 51°30'43"N 2°40'15"W | Industrial |
| Long Ashton (LA) | 51°26'08"N 2°38'41"W | Farm |
| Season | Sampling start date | Sampling End date |
| Spring | 2.2017 | 3.2017 |
| Summer | 6.2017 | 6.2017 |
| Autumn | 9.2017 | 11.2017 |
| Winter | 12.2017 | 2.2018 |

## SUPPLEMENTARY FIGURES

**Supplementary Figure 1.**
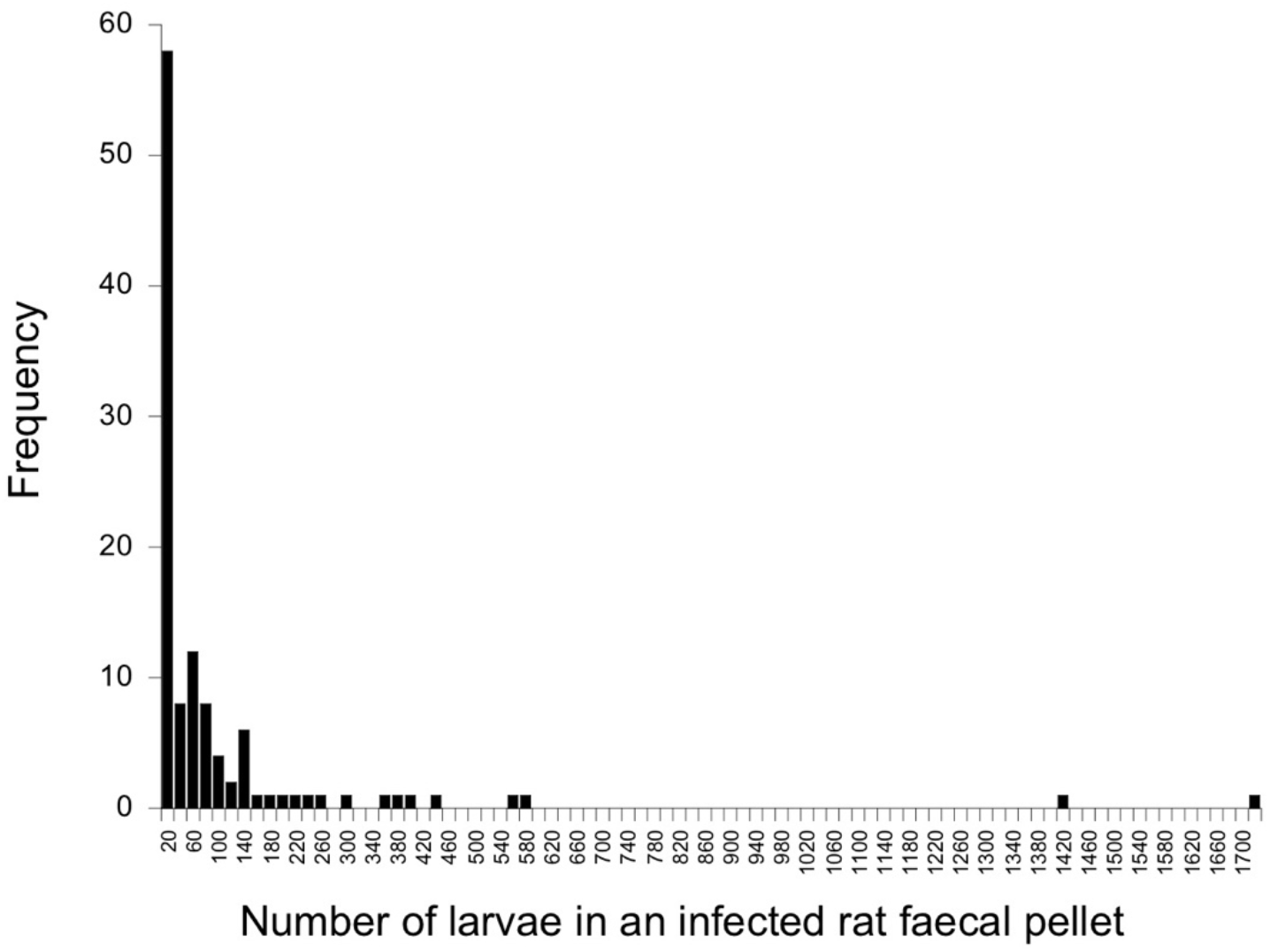
Frequency distribution of the number of *S. ratti* infective larvae isolated from infected rat faecal pellets. Uninfected pellets (N = 178) are not shown. The x-axis is in increments of 20 larvae, with only the upper limit shown, and only alternate increments labelled.

**Supplementary Figure 2.**
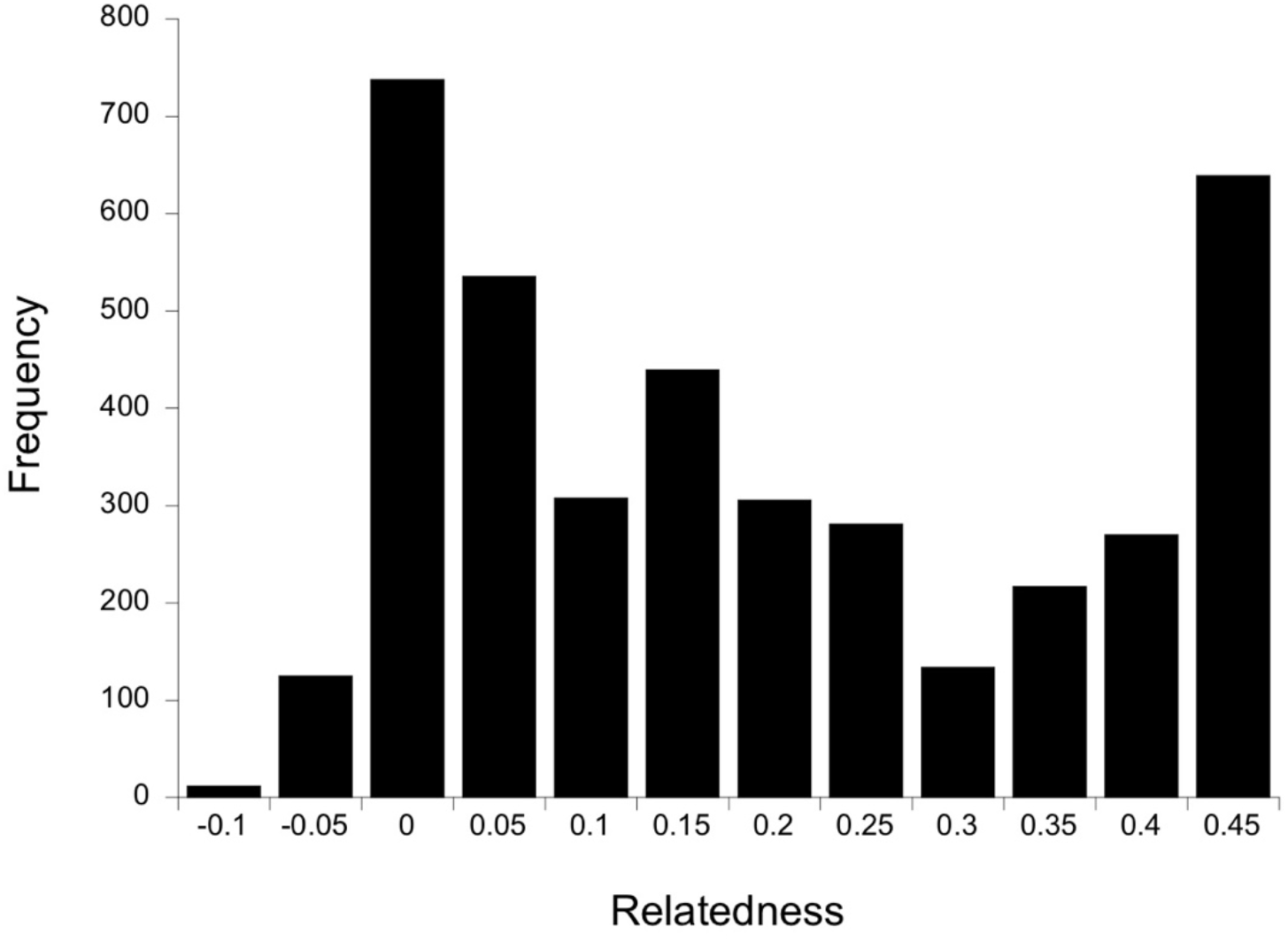
Histogram of Φ relatedness values among 90 *S. ratti* larvae. The x-axis is in increments of 0.05 with only the upper limit shown.

**Supplementary Figure 3.**
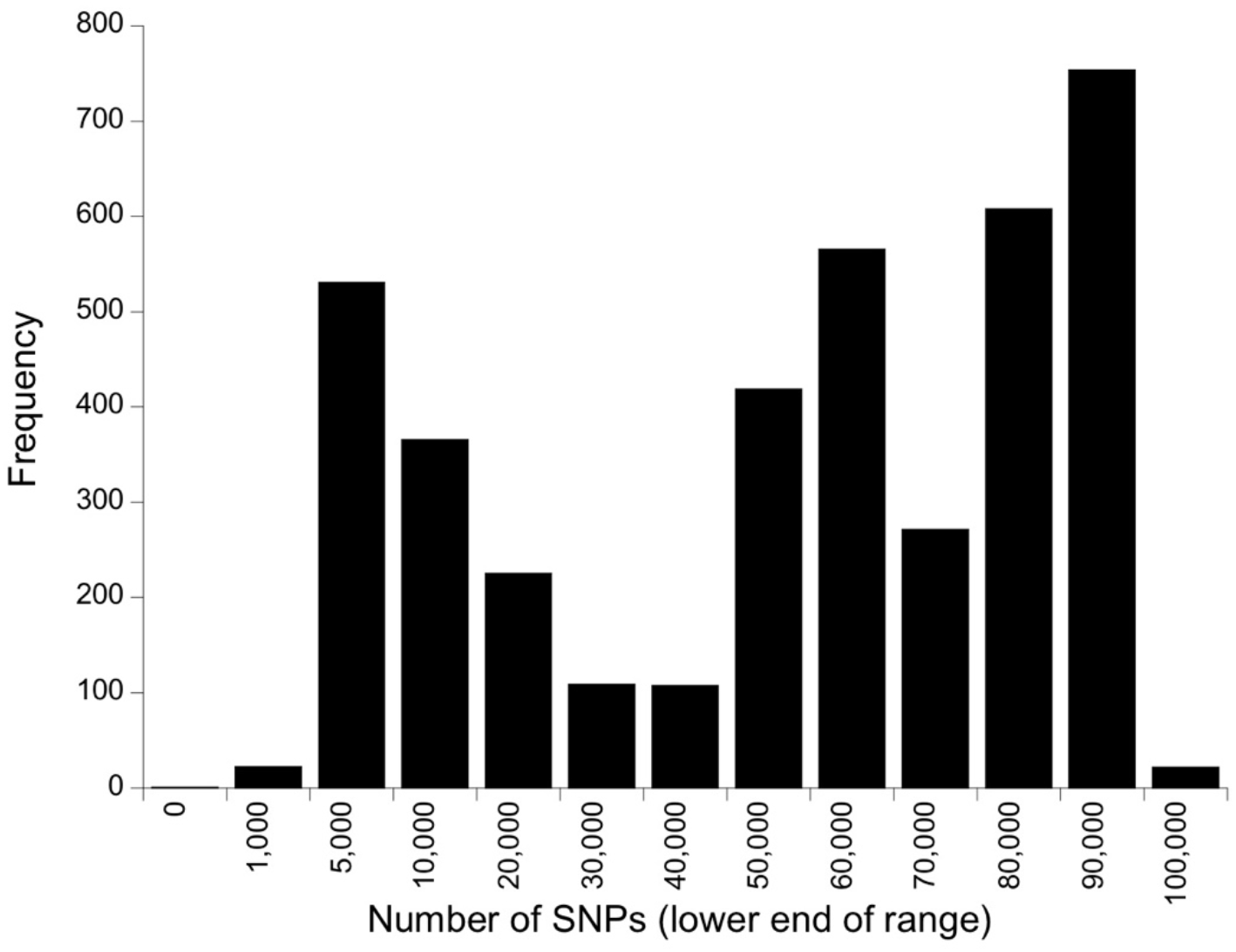
The frequency distribution of the pairwise number of SNP differences among the 90 parasites. The x-axis is in categories of 10,000, save for the first three which are 0-1,000, 1,000-5,000, 5,000-10,000; in all the lower end of the range is shown. The frequency is 1 in the first category.

**Supplementary Figure 4.**
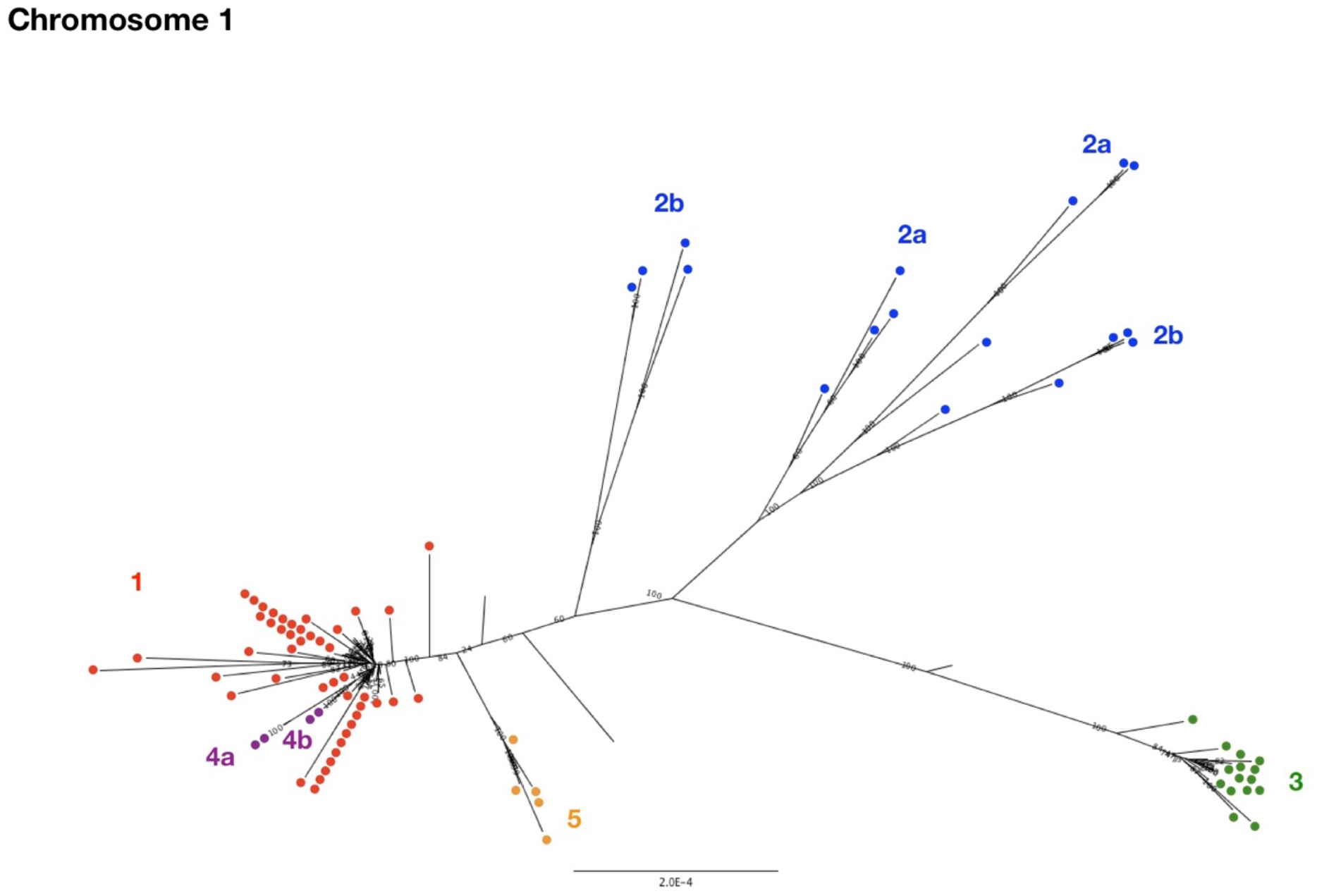
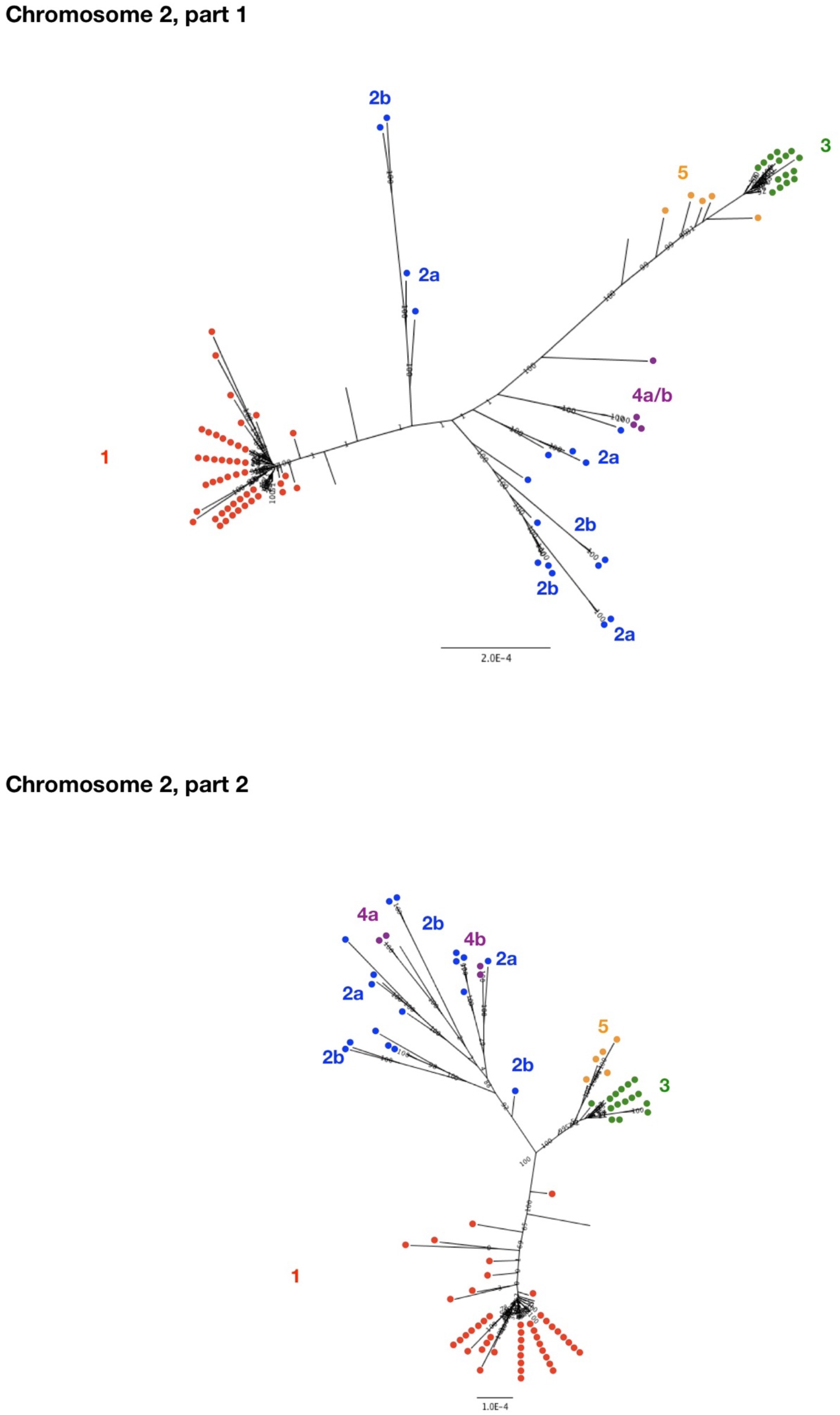
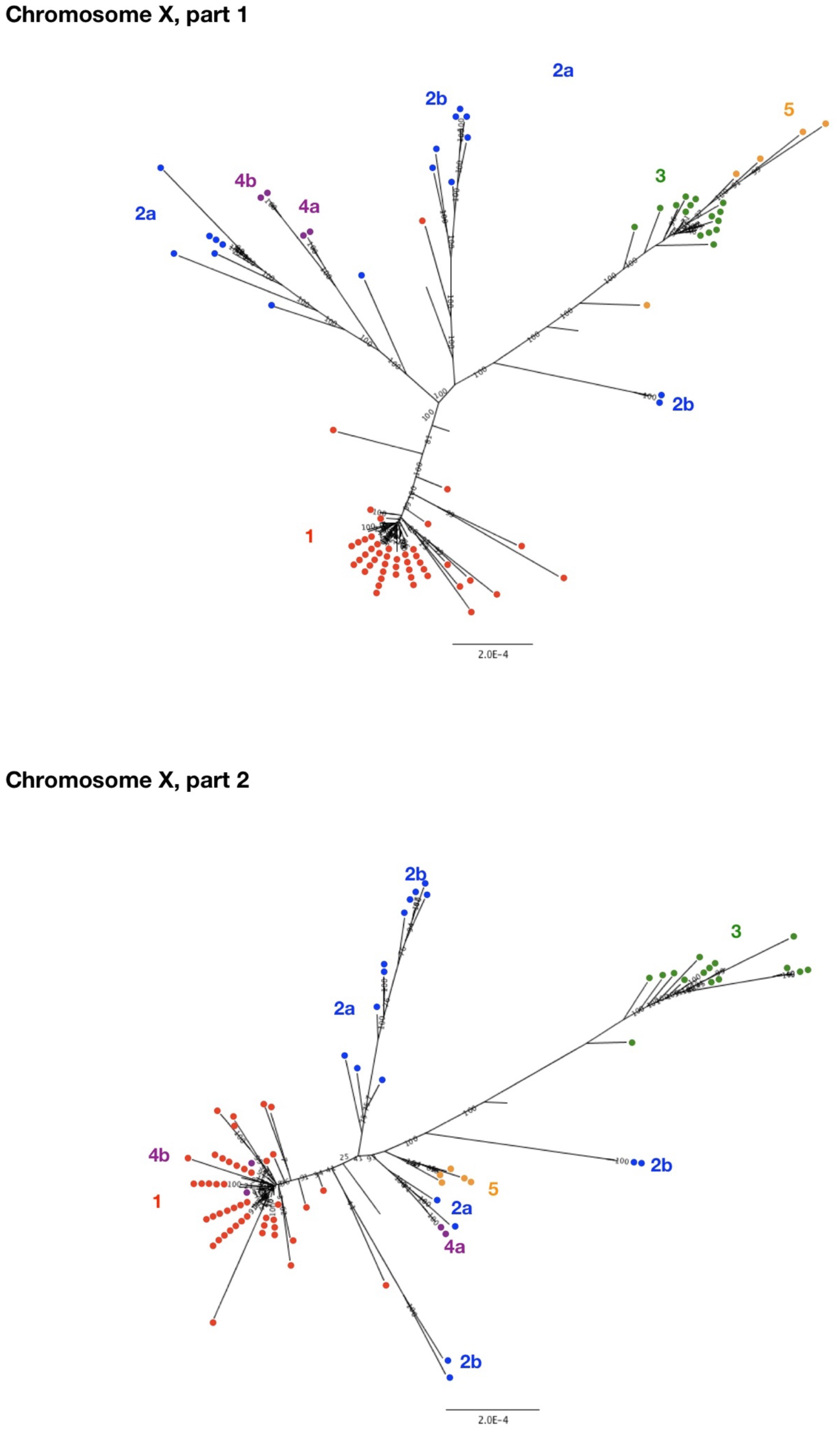
Maximum likelihood trees of the 90 parasites. In each tree, the support for each node is shown, and the clade membership of the worm is shown by the coloured dots, which correspond to the neighbour-joining tree (**Figure 3**). Trees are calculated based on analysis of chromosome 1 only, part 1 of chromosome 2, part 2 of chromosome 2, part 1 of the X chromosome and part 2 of the X chromosome. In all the scale bar is in units of substitutions per site.

**Supplementary Figure 5.**
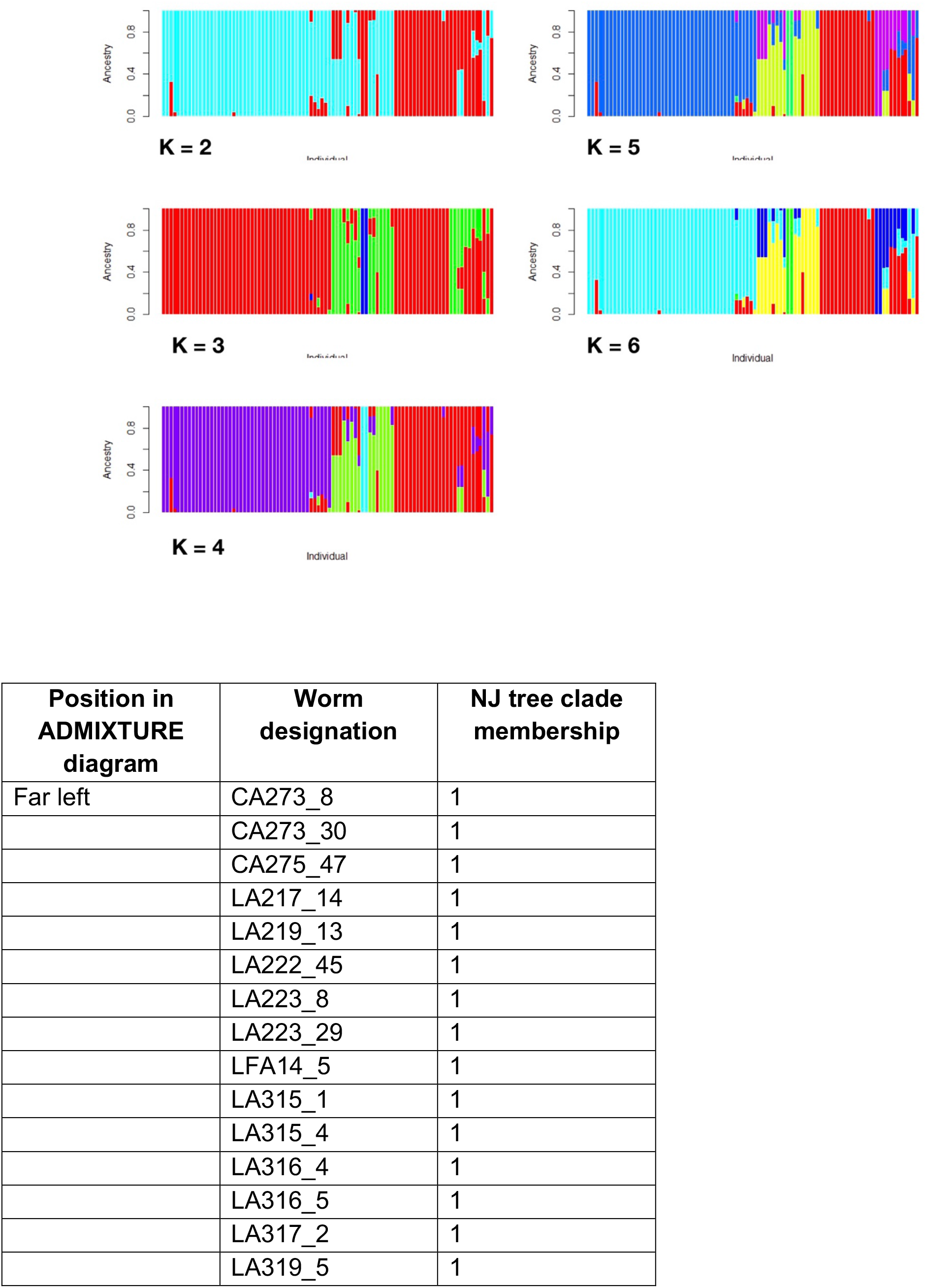
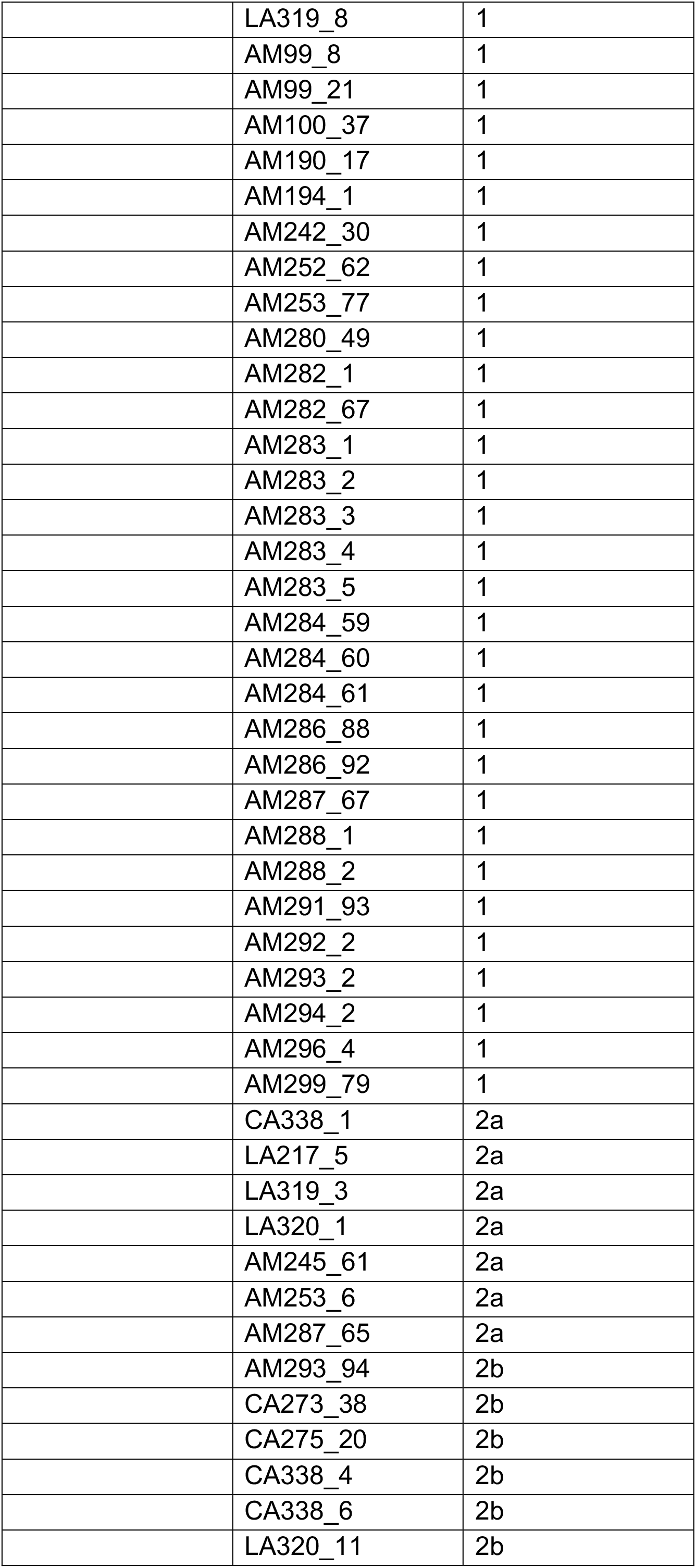
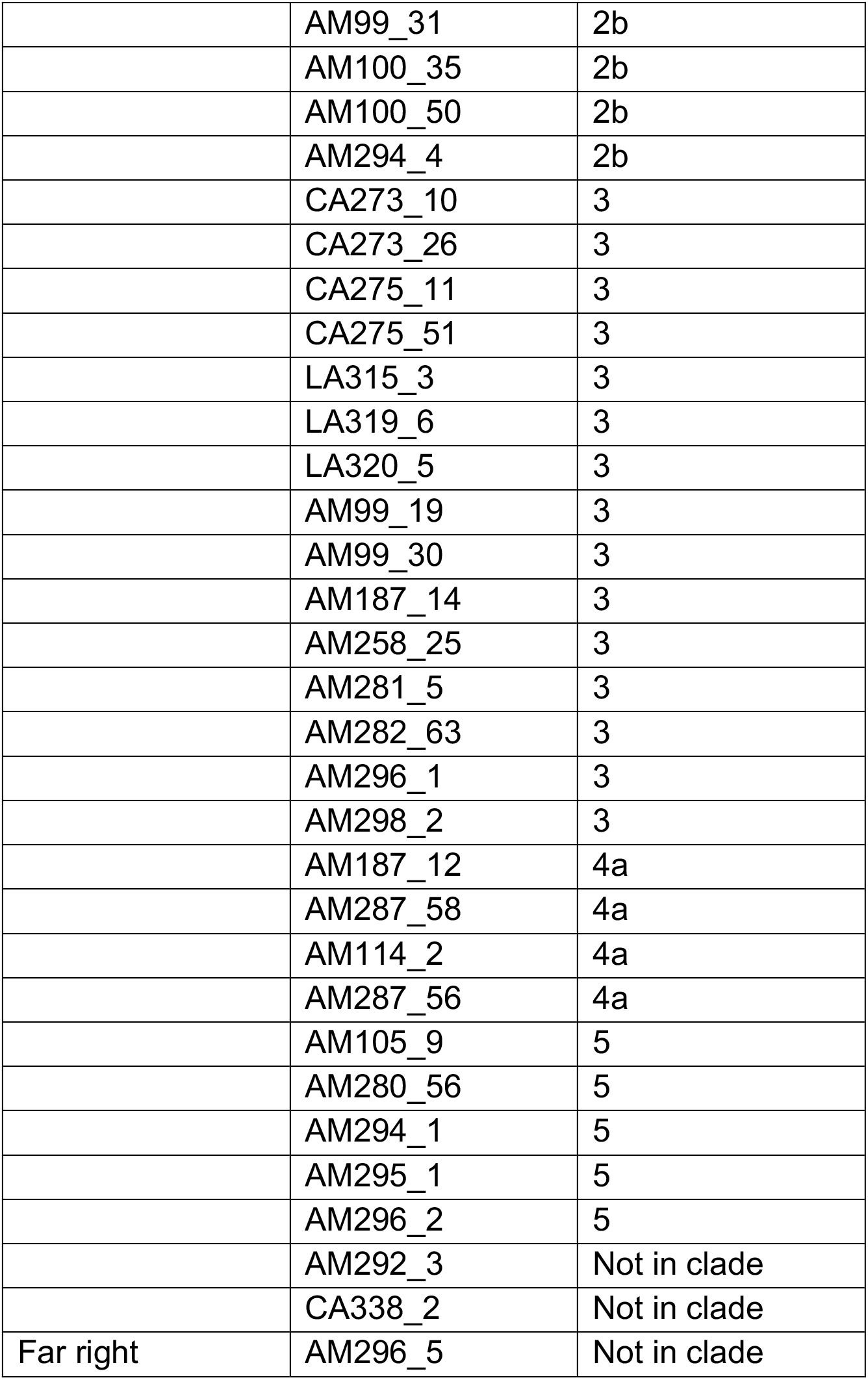
ADMIXTURE analysis of the 90 parasites. The ADMIXTURE diagrams are shown for K = 2, 3, 4, 5 (with cross validation errors of 0.22965, 0.16255, 0.16335, 0.16136, respectively). For K = 6, only five groups are detected. All diagrams have the same right to left order of worms, which is shown in a Table at the end of the figure.

**Supplementary Figure 6.**
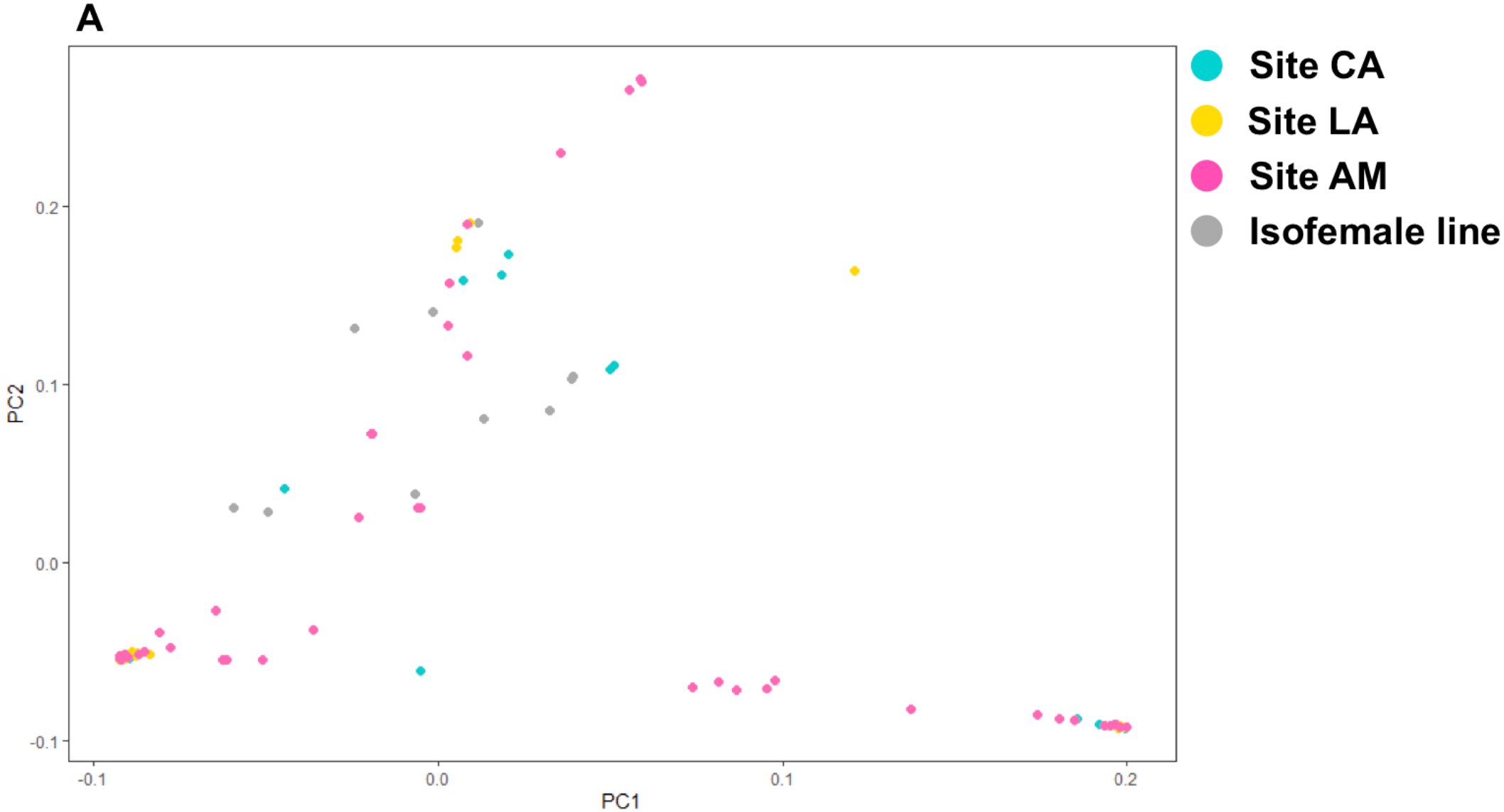
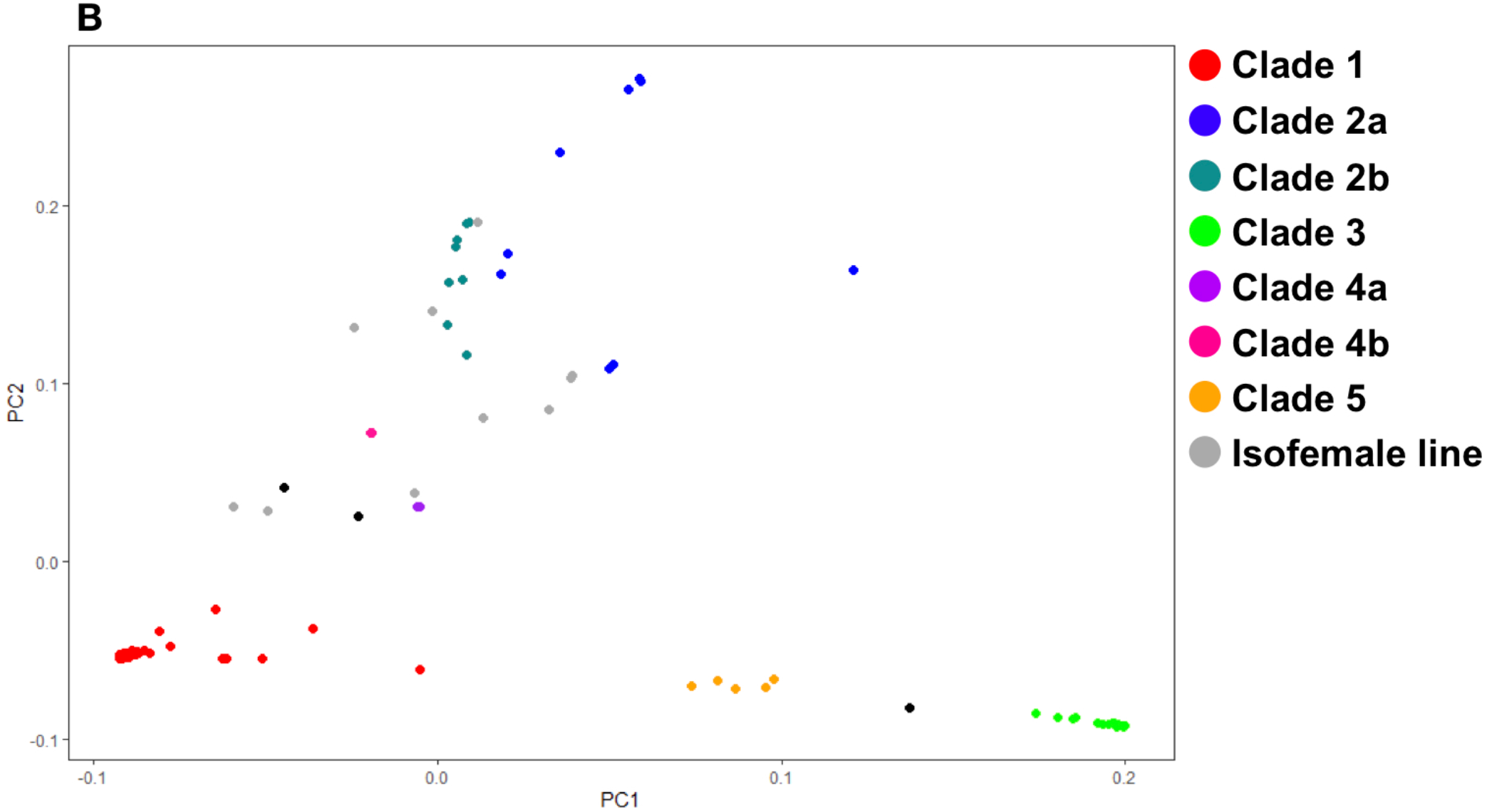
PCA analysis of *S. ratti* parasites. Projections of principal components (PC) 1 and 2 of the 90 parasites from the three samples sites and the 10 isofemale lines, where PCs 1 and 2 explained 91% of the variance. Individuals are coloured according to (A) sampling site and (B) neighbour-joining dendrogram clades (**Figure 3**) with the colour coding scheme shown in each panel.

**Supplementary Figure 7.**
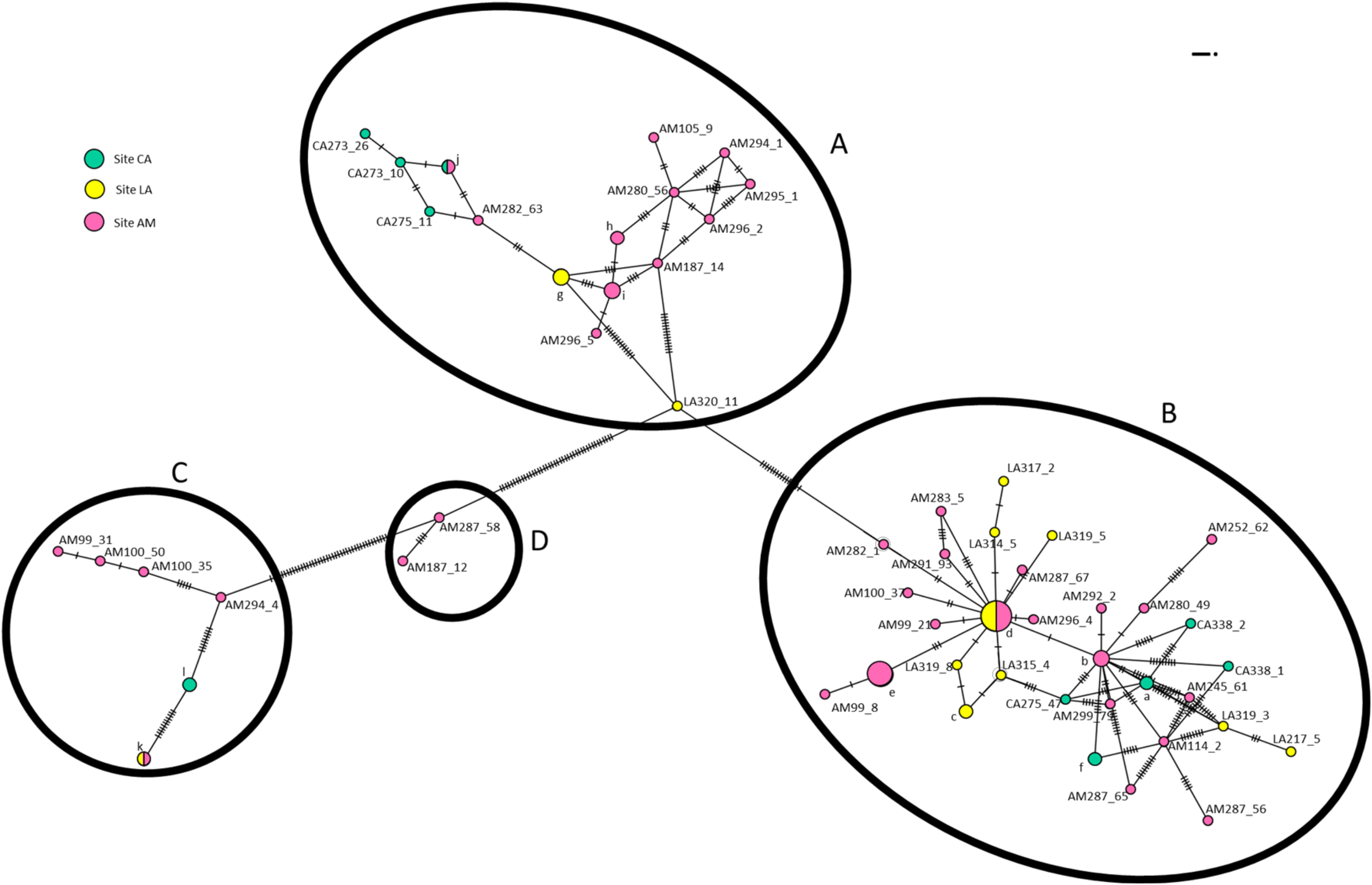
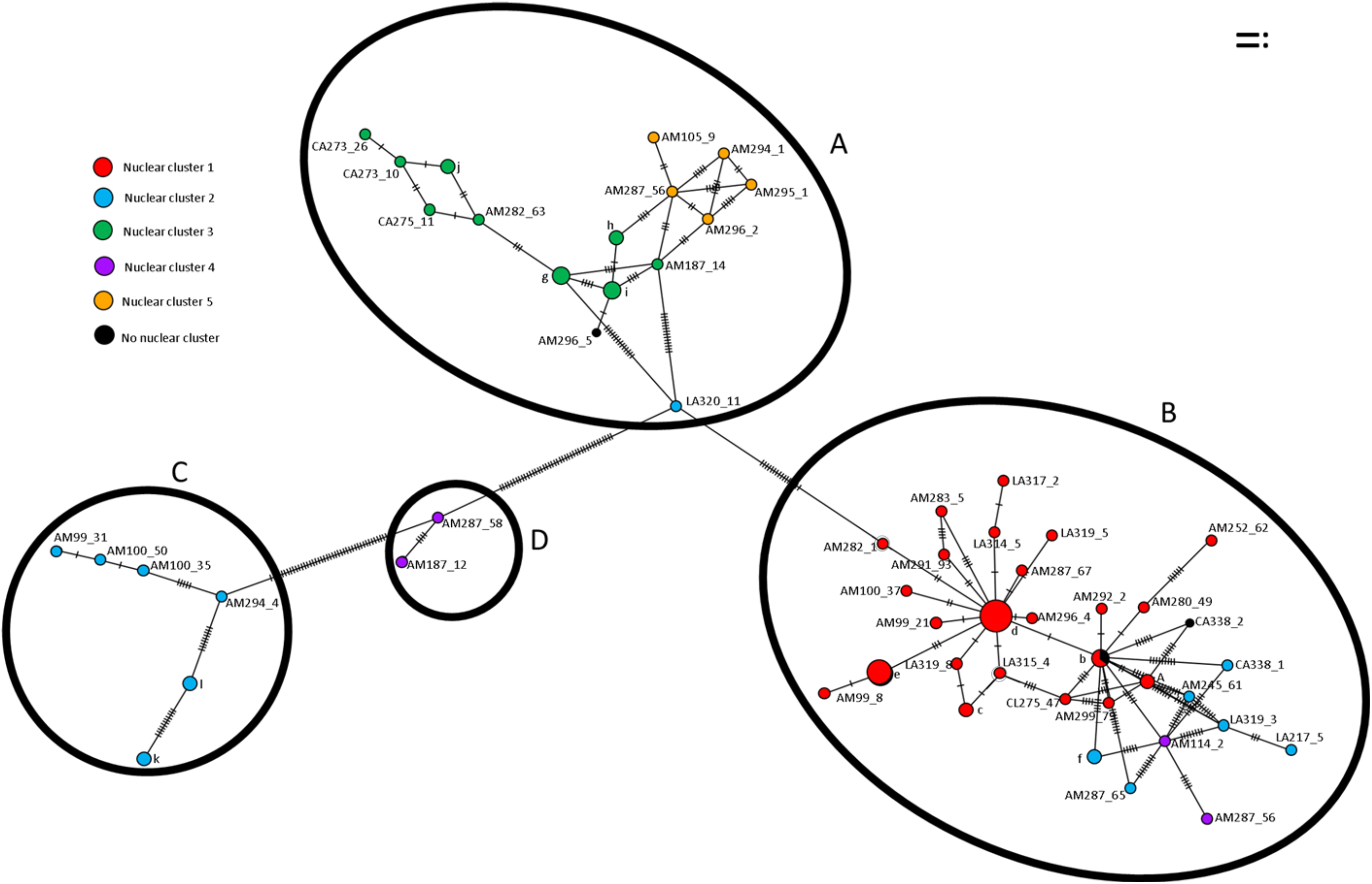
Minimum spanning *S. ratti* mitochondrial haplotype maps. Individual pellets are shown by their sampling site (AM, CA, LA) and the number preceding the underscore; the number after the underscore is the unique identifier of that worm. Haplotypes represented by multiple individuals are denoted by single letters. Four mitochondrial clades (A – D) are evident. Individual worms are coded either by the sampling site from which they were obtained or by the nuclear clade to which they belong (**Figure 3**). Mitochondrial clades A, B and C contained individuals from all 3 sampling sites, though at different rates. Mitochondrial clade A contains all individuals from nuclear clades 3 and 5 as well as one individual from nuclear clade 2b. Mitochondrial clade B contains individuals from nuclear clades 1, 2a and 4b. Mitochondrial clade C contains only individuals from nuclear clade 2b. The two individuals of nuclear clade 4a appear as intermediate between mitochondrial clades B and C in the haplotype map and so are designated as minor mitochondrial clade D.

**Supplementary Figure 8.**
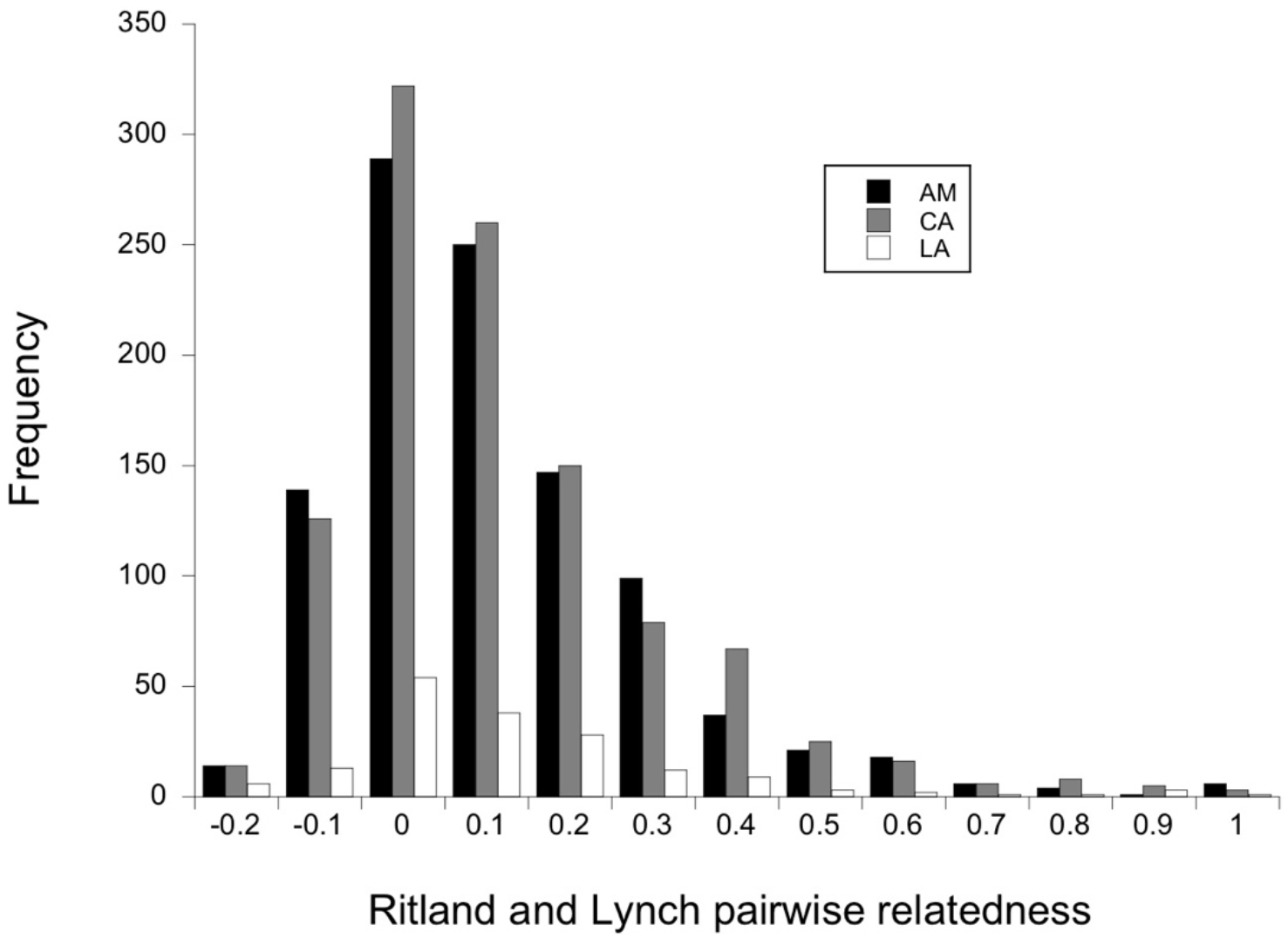
Ritland and Lynch pairwise relatedness values of rats within sampling sites. The x-axis is in increments of 0.1, with only the upper limit shown. At each site there is a right-hand skew to these distributions; Shapiro-Wilkes test for normality statistics W = 0.89 for site AM, W = 0.9. for site CA and W = 0.87 for site LA, P > 0.0001 in all cases.

**Supplementary Figure 9.**
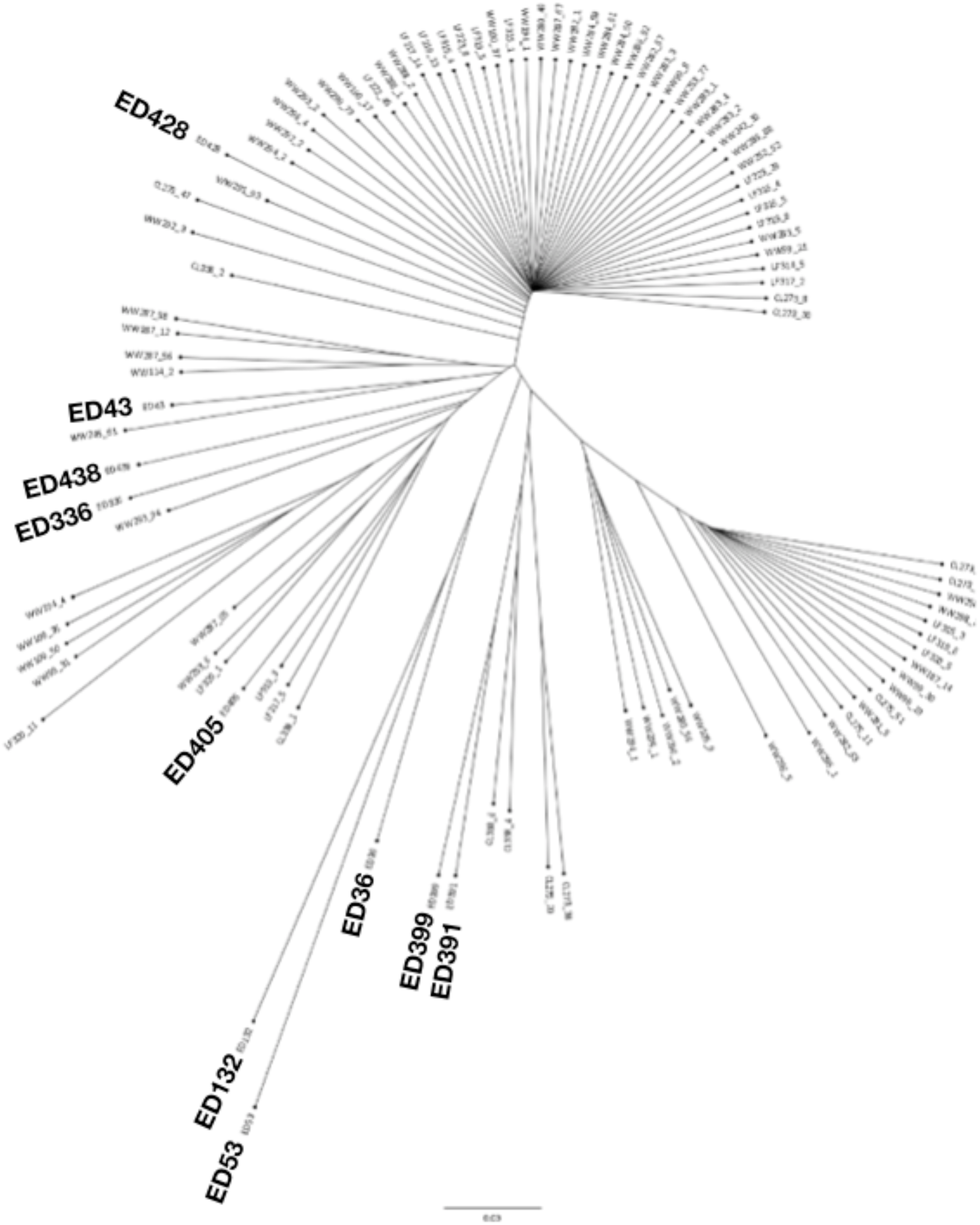
*S. ratti* neighbour-joining dendrograms of 10 isofemale lines and 90 larvae collected from the three sample sites. Isofemale lines are prefixed by ED (**Supplementary** Table 4). Five of the isofemale lines (derived from parasites collected from the southern UK in 1989-90, and Japan in 1990) are within clade 2b, which is the most diverse of the clades, but the other 5 isofemale lines are in clades 1, 2a and 4.

**Supplementary Figure 10.**
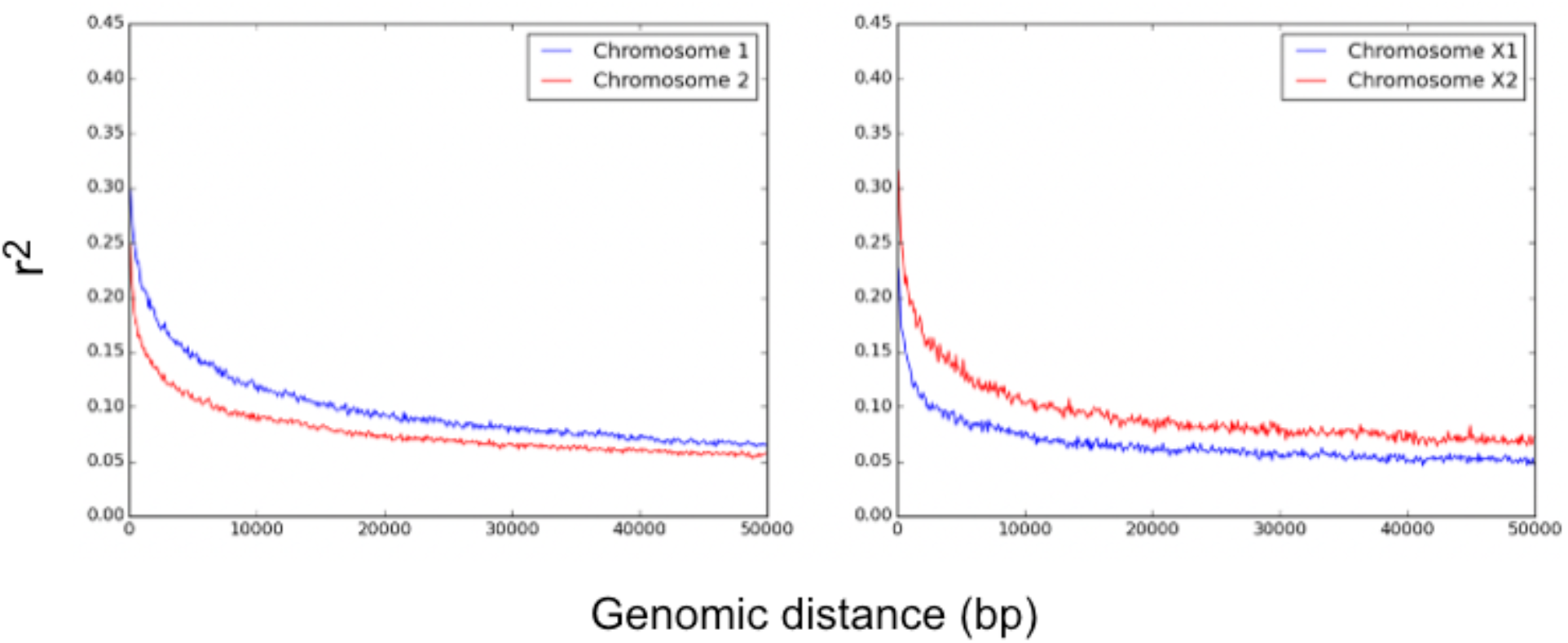
Linkage disequilibrium in the *S. ratti* genome. LD shown as r^2^ values. X1 and X2 are the largest and second largest X chromosome scaffolds, respectively.

**Supplementary Figure 11.**
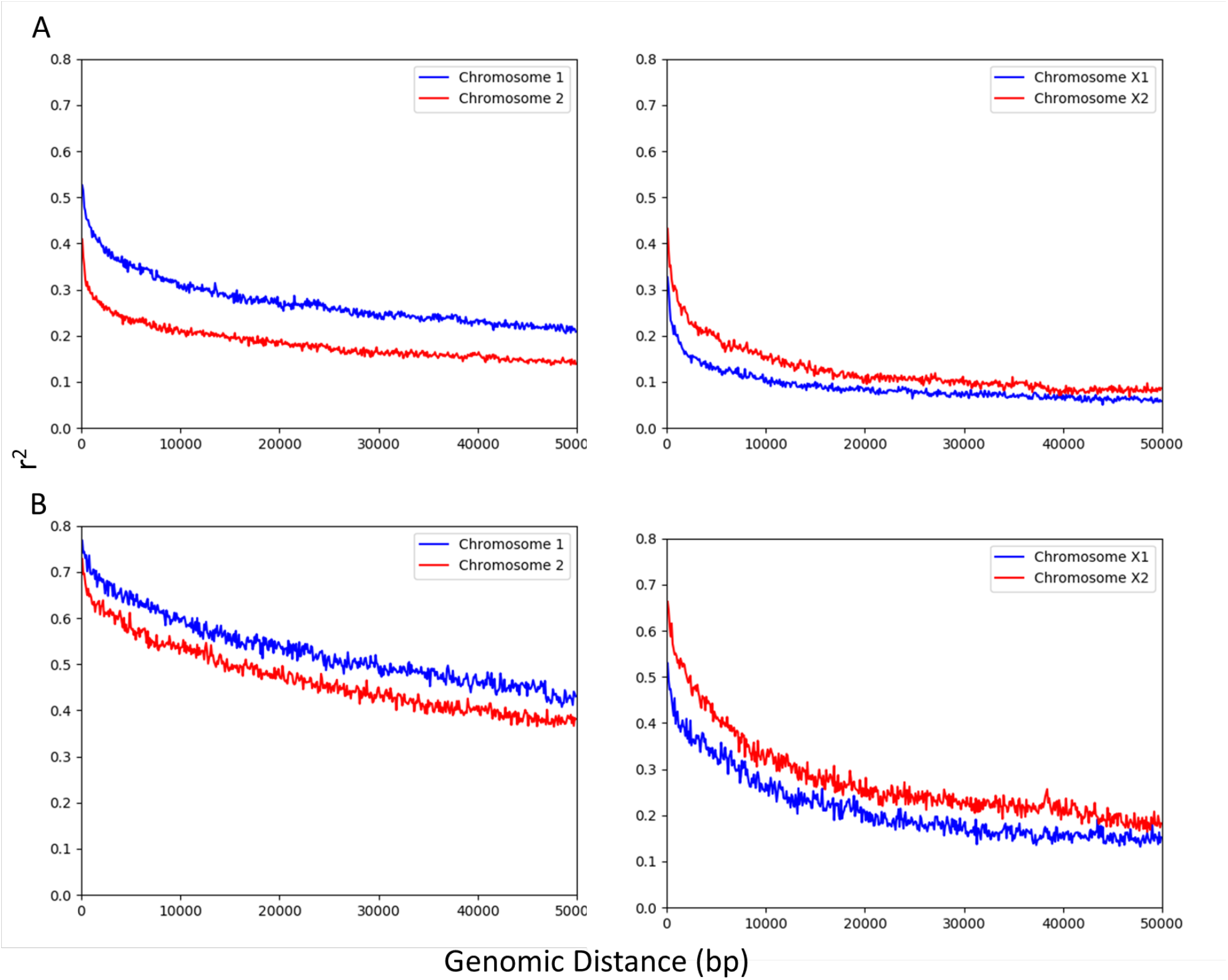
Linkage disequilibrium in the *S. ratti* genome for clade 1 and 3 parasites. LD is shown as r^2^ values for (A) clade 1 and (B) clade 3 parasites. X1 and X2 are the largest and second largest X chromosome scaffolds, respectively.

**Supplementary Figure 12.**
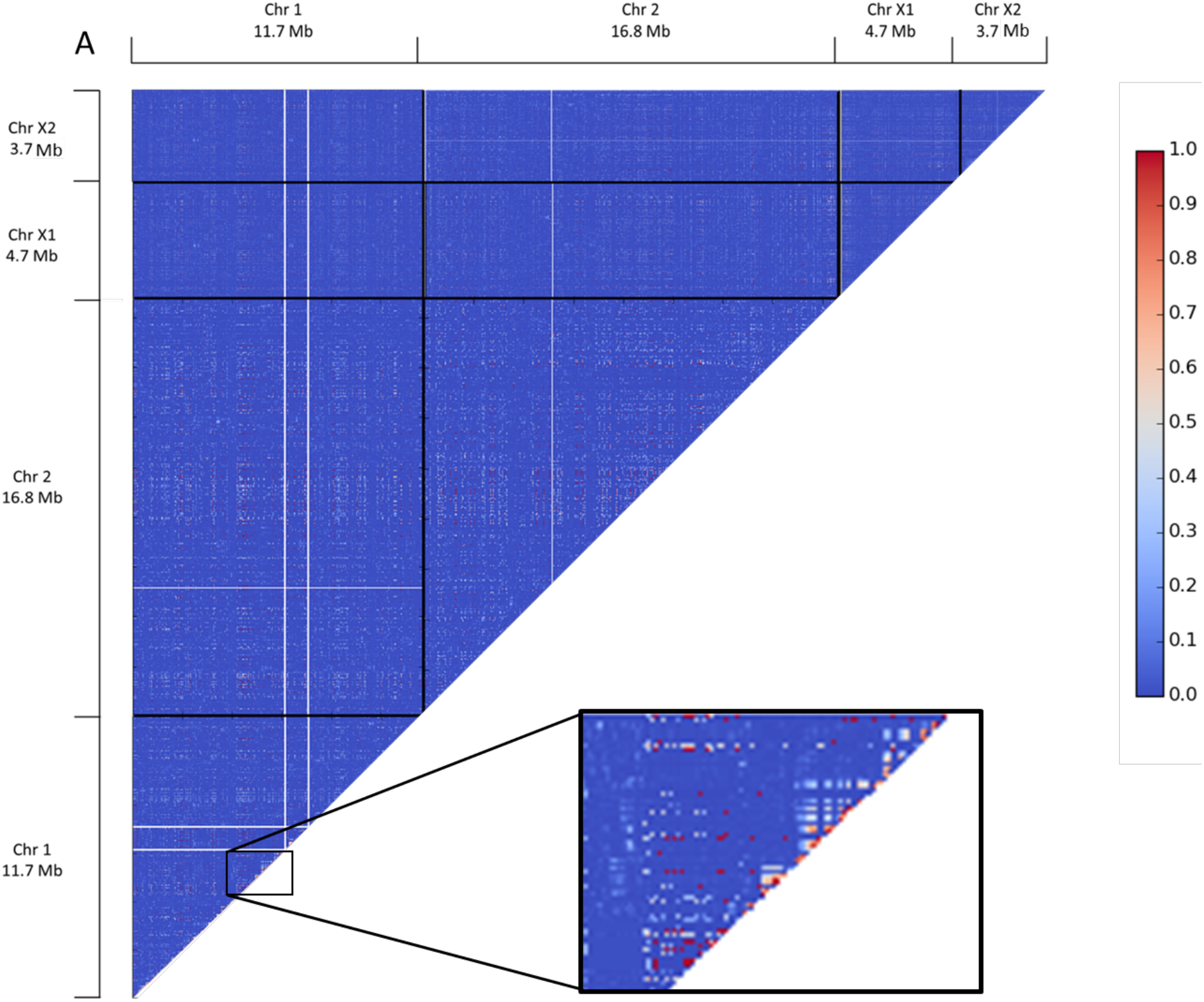
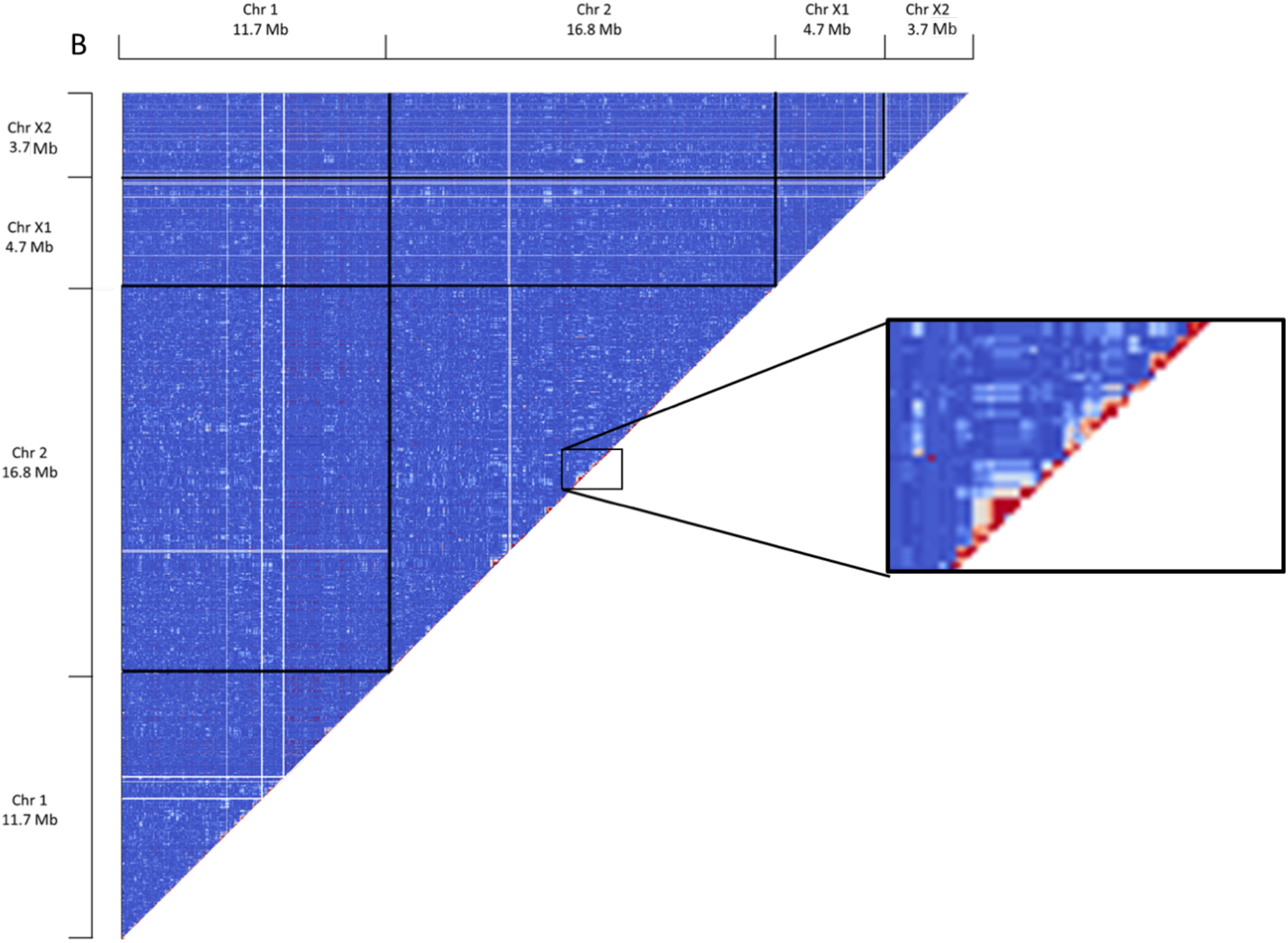
Heatmaps of linkage disequilibrium in the *S. ratti* genome for clade 1 and 3 parasites. LD is shown as r^2^ values on a coloured scale for (A) clade 1 and (B) clade 3. X1 and X2 refer to the largest and second largest scaffolds of the X chromosome, respectively. The inserts enlarge regions to show areas of high LD.

**Supplementary Figure 13.**
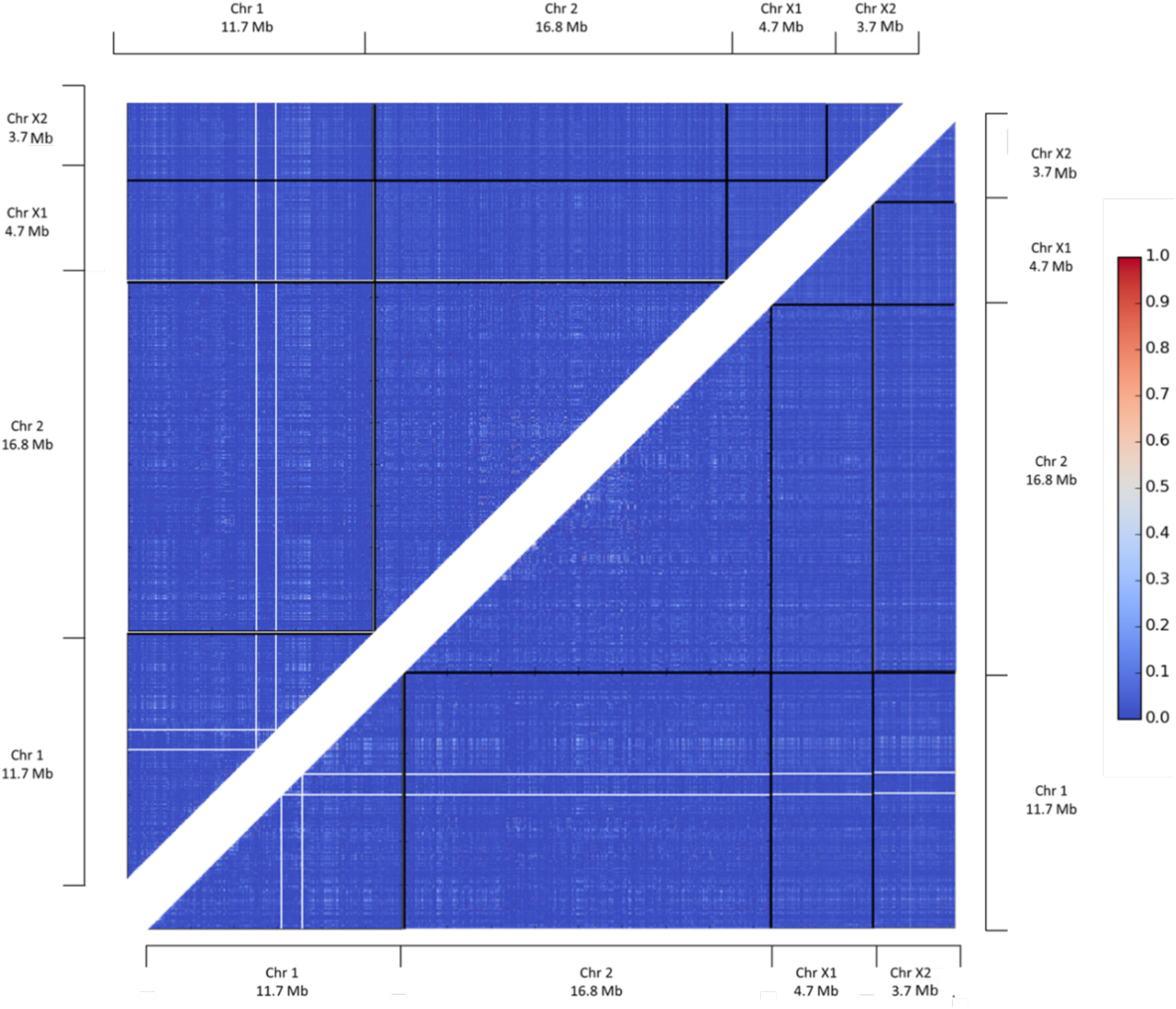
Heatmaps of linkage disequilibrium in the *S. ratti* genome. Phasing of genotypes was carried out by Beagle (above the diagonal) or Shapeit (below the diagonal). X1 and X2 refer to the largest and second largest scaffolds of the X chromosome, respectively. Vertical and horizontal white lines in chromosome 1 represent two megabase long tracts of Ns that separate the three X chromosome scaffolds, whose genomic order is not known.

**Supplementary Figure 14.**
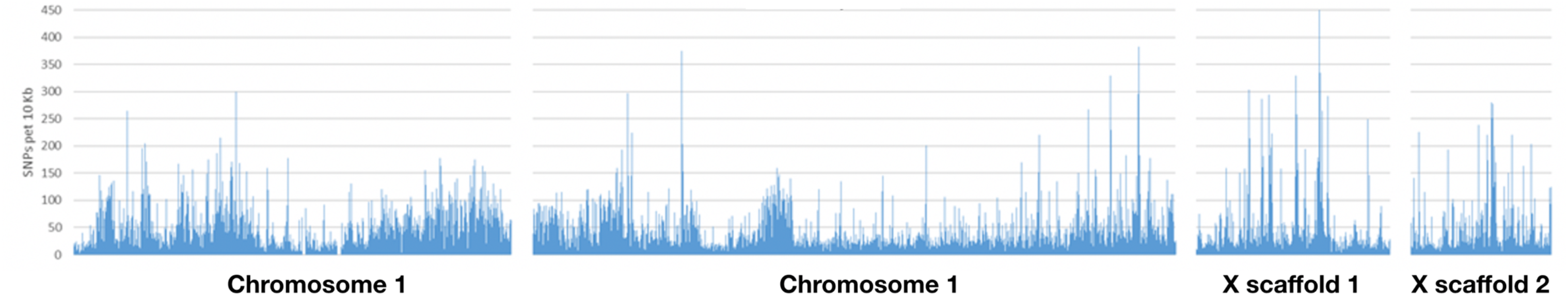
The distribution of SNPs across the *S. ratti* genome. The distribution of SNPs for all 90 parasites for discrete, 10 kb windows, each represented by a vertical bar, for chromosomes 1, 2 and the two scaffolds of the X chromosome.

**Supplementary Figure 15.**
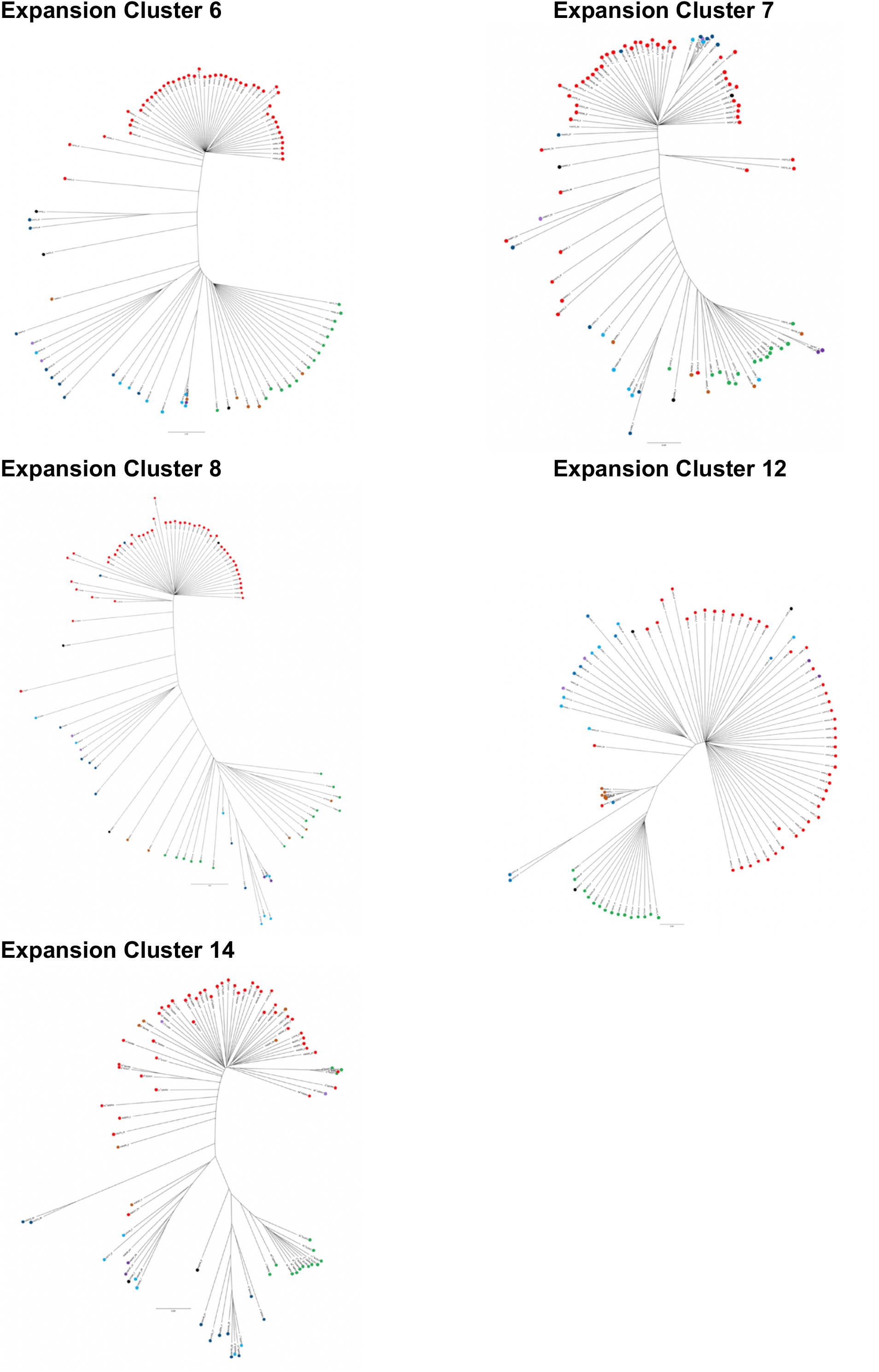
Neighbour-joining dendrograms based on five expansion clusters. Clusters, 6, 7, 8, 12 and 14 are the clusters from which no genes were excluded due to concerns about genome assembly. Each tree was calculated based on the entire sequence from the start to the end of each cluster (excluding flanking regions). Clades (1 –5) and sub-clades (2a and 2b, 4a and 4b) are defined based on the whole genome dendrogram (**Figure 3**). Individual parasites are marked with circles coloured according to their (sub-)clade in the whole genome dendrogram. Branch lengths are relative such that the distance between the two most distant individuals is 1, and so absolute distance differs among the panels.

**Supplementary Figure 16.**
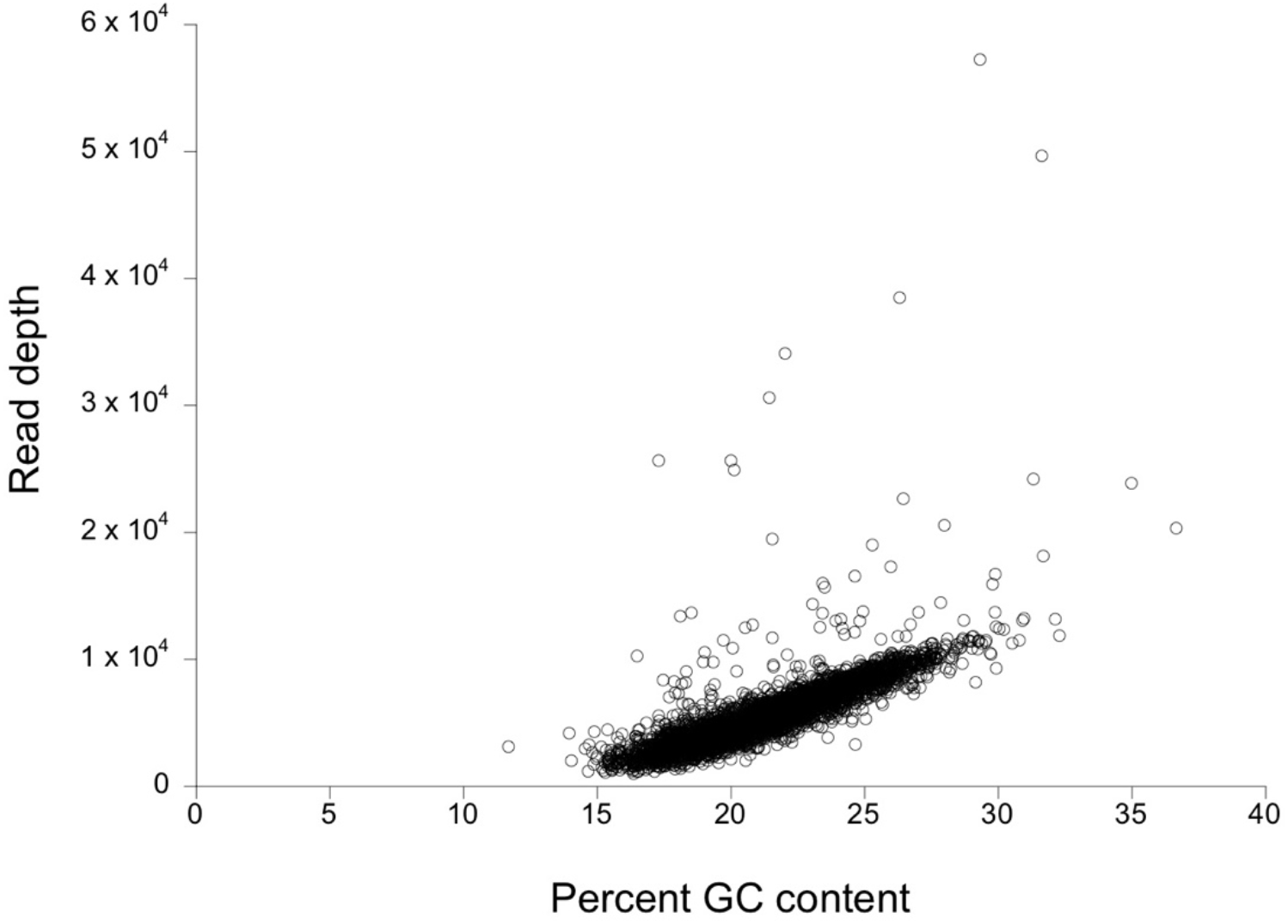
Correlation of read depth and GC content for 90 *S. ratti* larvae. GC content is measured in non-overlapping 10 kb regions. Correlation ρ = 0.783, *df* = 4,008, P < 0.00001.

